# Cohesin-dockerin code in cellulosomal dual binding modes and its allosteric regulation by proline isomerization

**DOI:** 10.1101/2019.12.19.882373

**Authors:** Andrés Manuel Vera, Albert Galera-Prat, Michał Wojciechowski, Bartosz Różycki, Douglas Vinson Laurents, Mariano Carrión-Vázquez, Marek Cieplak, Philip Tinnefeld

## Abstract

Cellulose is the most abundant organic molecule on Earth and represents a renewable and practically everlasting feedstock for the production of biofuels and chemicals. Self-assembled owing to the high-affinity cohesin-dockerin interaction, cellulosomes are huge multi-enzyme complexes with unmatched efficiency in the degradation of recalcitrant lignocellulosic substrates. The recruitment of diverse dockerin-borne enzymes into a multicohesin protein scaffold dictates the three-dimensional layout of the complex, and interestingly two alternative binding modes have been proposed. Using single-molecule Fluorescence Resonance Energy Transfer, molecular dynamics simulations and NMR measurements on a range of cohesin-dockerin pairs, we directly detect varying distributions between these binding modes that follow a built-in cohesin-dockerin code. Surprisingly, we uncover a prolyl isomerase-modulated allosteric control mechanism, mediated by the isomerization state of a single proline residue, which regulates the distribution and kinetics of binding modes. Overall, our data provide a novel mechanistic understanding of the structural plasticity and dynamics of cellulosomes.

## Introduction

The plant cell wall, featuring cellulose and hemicellulose, is the most abundant source of energy and carbon of the biosphere (*1*). Due to its structural complexity and chemical heterogeneity, this network of polysaccharides is highly resistant to enzymatic degradation. Some anaerobic bacteria, have developed the aptly named cellulosome, a huge multi-enzyme extracellular complex able to degrade it with high efficiency (*1, 2*). Present in diverse ecosystems like compost piles, vertebrates and invertebrates microbiota, hot spring pools, forest and pasture soil, cellulosome-producing organisms play a major role in the carbon turnover (*3, 4*).

The first-ever described cellulosome, that of *Clostridium thermocellum* (*5*), recapitulates well the main features of these enzymatic complexes. The primary scaffoldin CipA, a multimodular non-catalytic subunit, tethers the whole complex to the substrate and to the cell surface, while it serves as the binding platform for secreted cellulolytic enzymes. A series of type-I cohesin modules, separated by flexible linker regions, act as the anchor points to enzyme-borne dockerin type I modules. While the high affinity type-I cohesin-dockerin interaction dictates the supramolecular assembly of the cellulosome, the C-terminal type-II dockerin attaches the complex to the cell surface, and the carbohydrate-binding module (CBM) binds to the cellulosic substrate (*1, 6, 7*) (Fig 1A). Interestingly, the dockerin type-I modules from different enzymes display similar affinity to the different cohesins of the scaffoldin, potentially binding to any position with equivalent probability (*8*). Substrate targeting, and the spatiotemporal coordination of different enzymatic activities create a synergistic effect resulting in the highly efficient degradation of cellulosic material (*9, 10*).

**Figure 1.**
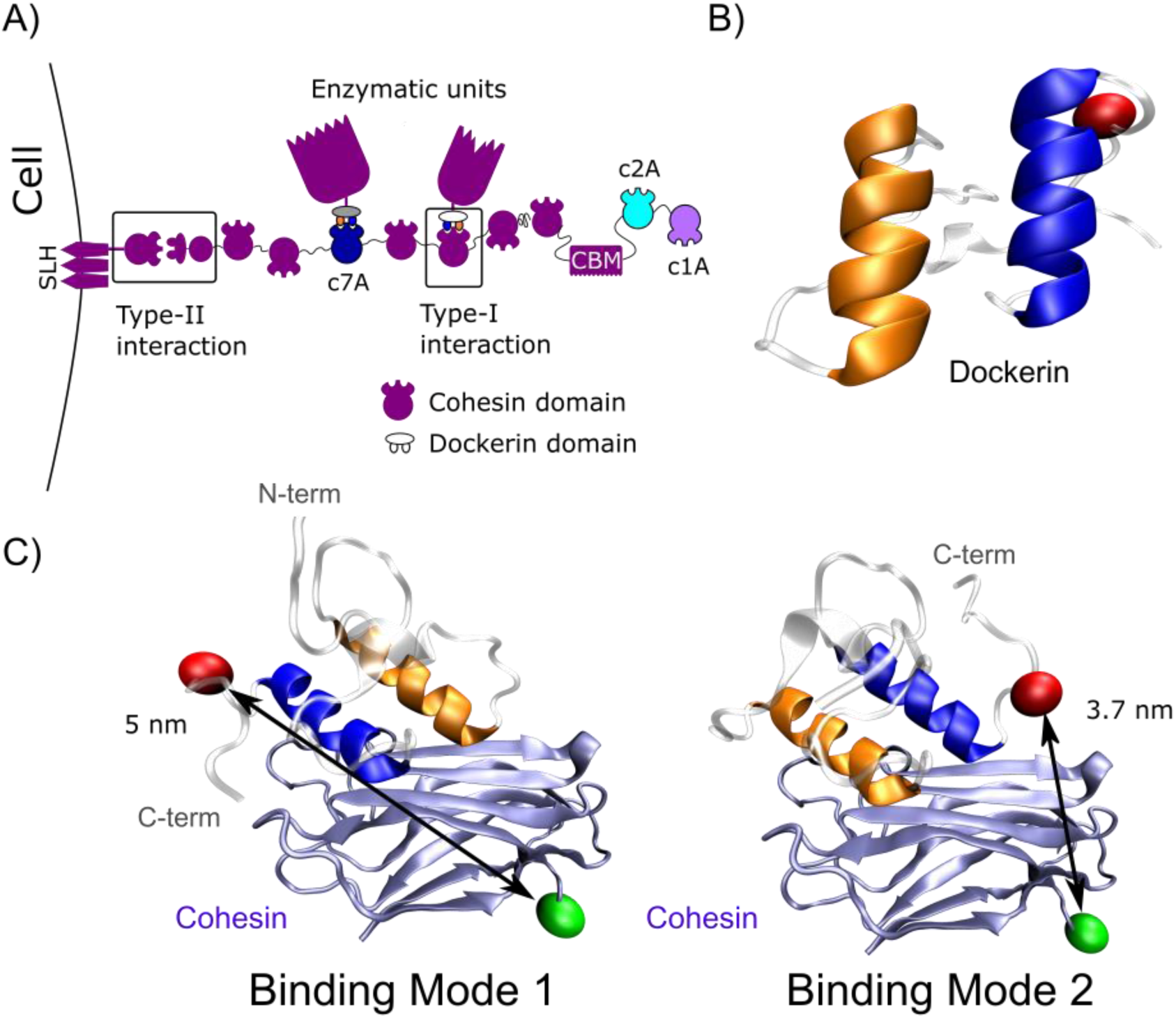
Dual binding mode of cohesin-dockerin complexes. **A)** Cartoon representation of the cellulosome of *C. thermocellum*. CBM domain on scaffoldin CipA mediates cellulose binding while the whole cellulosome is attached to the cell surface via a type-II cohesin-dockerin interaction. Type-I cohesin-dockerin interaction is essential to cellulosome self-assembly, and allows the incorporation of diverse dockerin-borne enzymes in any of the nine cohesin domains of Cip A scaffoldin protein. The cohesins studied, c2A (cyan), c1A (violet) and c7A (dark blue) are highlighted, and the color code used in the sketch followed in the rest of the figures. **B)** Helices symmetry in type I-dockerins. Structure of dockerin module from cellulase S (PDB 2MTE) with the helix H1 represented in orange and in the helix H3 in blue. The symmetry is evident, a 180° rotation around an axis perpendicular to the figure plane will produce essentially the same structure but with helices position exchanged. The red sphere represents the attachment point of the Alexa Fluor 647 dye. **C)** Dual binding modes of cohesin-dockerin complexes. On the left, model structure of a cohesin-dockerin complex in B1 mode, on the right in B2. The C-termini distances are different among the two binding modes, 5 nm between the red/green highlighted residues in the B1 mode and 3.7 nm in B2 mode. The red sphere represents the attachment point of the Alexa Fluor 647 dye (R63C), and the green sphere the Cy3B dye position (G144C).

The high catalytic efficiency of cellulosomes arises from the balanced combination of enzymatic synergy, structural plasticity and dynamic adaptation to the substrate. The flexible intermodular linkers, the vast diversity of enzymes, and the potential binding of enzymes to any cohesin in the scaffolding allow the complex to adjust to a substrate that is heterogeneous in nature and changes continuously during the degradation process (*9, 11-19*). On top of this, another level of structural plasticity has been proposed. Two repeated segments displaying high sequence and structural similarity, Helix 1 and Helix 3, make type-I dockerin a very symmetrical module (*20*) (Fig 1B). Mutational studies, and the module symmetry led to the idea of cohesin-dockerin complexes binding in two alternative conformations (*21-23*). Structures of mutant dockerins bound to cohesin in two binding modes have been obtained (*23-25*), and differs by a 180° rotation of the dockerin around an axis perpendicular to the helices plane (Fig 1C). In one of the binding modes, from now on referred as binding mode 1 (B1), the dockerin’s Helix 3 supports most of the contacts with the cohesin module, while in the alternative binding mode 2 (B2), a similar set of contacts was sustained by Helix 1. These alternative binding modes could allow cellulosomes to display different enzymatic arrangements avoiding steric hindrances, leveraging synergy or avoiding anti-synergistic effects, and enabling the enzymes to rapidly adopt an alternative conformation for interaction with intact parts of the substrate. Although extensive evidence for the dual binding mode has been presented (*21-26*), direct detection (*27*) and quantitative analysis on wild type complexes is still lacking, hindering its characterization and therefore the understanding of its role in cellulosome function and regulation.

Cellulosome composition is known to adapt to the substrate type and presence of extracellular polysaccharides, and temporal changes in its composition during cellulose degradation has been described (*28-30*). Although *de novo* synthesis of whole cellulosome components for adaptation is possible, it is clear that cellulosome-remodeling is the less metabolic-demanding solution to adapt the cellulosome to the changing environmental conditions. Currently, the way cellulosomes change their composition is unknown (*12, 16*), but any active mechanism will necessarily involve the modulation of the cohesin-dockerin interaction. As a key element in cellulosome self-assembly, it is crucial to understand the role of cohesin-dockerin interaction in its fine structure and regulation.

Here, we studied the dual binding mode phenomenon using single-molecule Förster resonance energy transfer (smFRET), molecular dynamics simulations and NMR measurements. Our single-molecule approach allows for direct and easy identification of two binding conformations in various type-I cohesin-dockerin complexes and the quantitative determination of the populations using photon distribution analysis (PDA). Our data reveal a cohesin-dockerin code for the size of the binding mode populations, an allosteric control mechanism as well as an unexpected dynamic behavior between binding modes. Furthermore, we report the control of the dual binding mode and its dynamic behavior by the isomerization state of a single proline, which can be externally modified by a prolyl isomerase. These results have a direct impact on our understanding of the structural organization and regulation of cellulosomes, as well as the role of the dual binding mode on these processes. Overall, we have unveiled a novel allosteric mechanism that could be exploited for cellulosome remodeling and dynamic regulation.

## Results

### smFRET enables direct dual binding mode detection

The starting point of our work are the PDB structures 1OHZ (*25*) and 2CCL (*23*) from *C. thermocellum*. In these structures, two different mutants of dockerin Xyn10B are interacting with their cognate CipA cohesin 2 module (c2A) in the two alternative binding modes (B1 and B2 respectively). After visual inspection, we realized that the distance between the C-termini of cohesin and dockerin modules were different between both binding modes. As a few C-terminal residues from the dockerin are missing in both structures, and in order to better estimate distance differences, we aligned the wild type full length dockerin CelS (PDB:2MTE (*20*)) with the dockerin in 1OHZ and 2CCL. Fig. 1C shows the dockerin CelS bound to c2A in the B1 (left) or the B2 mode (right). Using these structures, we estimated the distance between α-carbons of cohesin’s G144 and dockerin’s D63 (see methods for a discussion on the selection of residues) to be about 5 nm and 3.7 nm in the B1 and B2 mode, respectively. smFRET is ideal for studying biomolecules when the distance between acceptor and donor fluorophores are within this range, and due to the distance dependency of the energy transfer efficiency (*E*) (*31, 32*), 5 and 3.7 nm dye-separation should report lower and higher E values respectively. Thus, we exchanged these residues to cysteine and attached our donor (Cy3B in G144C) and acceptor dye (Alexa Fluor 647 in D63C) using maleimide chemistry (see Methods).

Firstly, we studied the complex between c2A and dockerin CelS (c2A-celS) by smFRET on freely diffusing molecules. The FRET histogram of the complex shows a bimodal distribution, with a subpopulation of low FRET efficiency separated from a second high-FRET subpopulation (see Fig. 2A, striped histogram), as expected for the B1 and the B2 modes. In order to confirm this assignment, we used single binding mode mutants (SB), which are only able to bind in the B1 (celS_SB1) or the B2 mode (celS_SB2). The FRET histogram of c2A-celS_SB1 mutant shows one discrete population with a FRET value that rightly matches that of the B1 subpopulation of the wild type protein (blue line, Fig. 2A). Similarly, the c2A-celS_SB2 complex presents a single population as well, and its maximum concurs with the wild type’s B2 subpopulation (orange line, Fig. 2A). The same type of bimodal distribution was also observed in surface-immobilized c2A-celS complexes (Figs. S1A and S1B).

**Figure 2.**
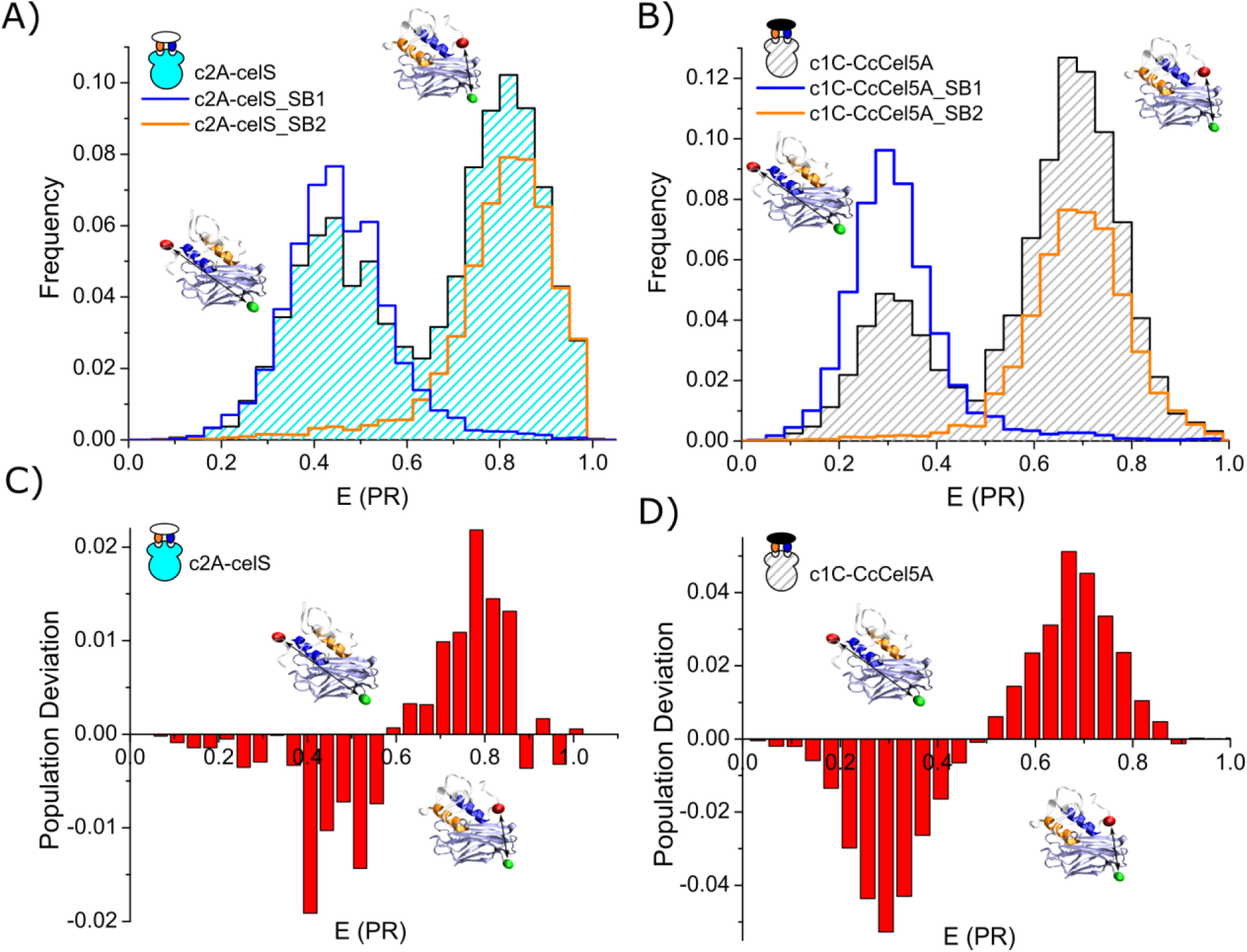
Direct detection of dual binding modes by smFRET. **A)** FRET efficiency histogram of the c2A-celS complex from *C. thermocellum*. smFRET data of free diffusing molecules shows two different population of molecules, one with higher E value that the other. We expect lower FRET values for the B1 mode than for the B2 due to the larger estimated C-termini distance. We double-check this assignment with single binding mode mutants (SB). SB1 in blue and SB2 in orange represent single binding mode mutants, only able to bind in B1 or B2 respectively. They effectively show a single population, which maxima coincides with the B1 and B2 peak of the wild type complex. **B)** E histogram of free diffusing c1C-CcCel5A complexes from *C. cellulolyticum*. The histogram shows the two different B1 and B2 modes as well, as confirmed by SB mutants (SB1 mutant in blue, and SB2 mutant in orange). All histograms were area-normalized to 1, for SB mutants, each mutant was normalized to 0.5 for comparison purposes. **C)** and **D)** In order to highlight the different distribution of B1 and B2 subpopulations in *C. thermocellum* and *C. cellulolyticum* complexes, we compared them with a population of equally-distributed binding modes. **D)** Plot showing the subtraction of a representative histogram of c2A-celS complex (B1 fraction of 0.43) and an artificial population of equally-represented single binding mode mutants (B1 fraction ≈ 0.50, see supplementary results for a detailed explanation on these artificial data sets). Similarly, **D)** shows the analog plot for c1C-Ccel5A *C. cellulolyticum* complex (B1 fraction of 0.26). All histograms were area-normalized before subtraction.

Type-I cohesin-dockerin complexes from *Clostridium cellulolyticum* have been proposed to display also a dual binding mode. Crystallographic structures of SB mutants are also available for this complex, and we took advantage of the high structural similarity between *C. thermocellum* and *C. cellulolyticum* complexes (*24*) and applied the same experimental approach (Fig. S2). Specifically, we studied the interaction between the cohesin 1 from scaffoldin CipC, and the dockerin from the cellulase 5A (c1C-CcCel5A complex). Similarly, the solution smFRET histogram of c1C-CcCel5A displays a double peak shape, featuring a high and low FRET population, which we confirmed as the B1 and B2 modes with the help of SB mutants (Fig. 2B). Finally, we also detected both binding modes in surface-immobilized molecules (Fig. S1A and S1B).

### Uncovering a Dual Binding Mode code

Visual inspection of the c1C-CcCel5A histogram shows that both binding modes are not equally populated, prevailing the B2 mode over B1 (Fig. 2B and 2D). This trend is also observed, although more subtly, in the c2A-celS complex. In the latter case, it is more difficult to notice from inspection of the FRET histograms (Fig. 2A), but it becomes obvious when compared with an histogram of SB1 and SB2 mutants with equal populations (Fig. 2C).

From the FRET histograms, it is clear that the binding modes are not only unevenly populated but that the size of the B1 and B2 populations is different among complexes. To get further insights into this phenomenon, we used PDA of smFRET data (*15, 31, 33, 34*) to quantify the distribution of binding modes of our cohesin-dockerin complexes. Firstly, we tested the ability of PDA to retrieve accurate values for the fraction of binding mode populations in our bimodal smFRET histograms. To this end, we used SB mutants data to create artificial data sets with controlled subpopulation ratios, and then used PDA as implemented in the free software PAM (*35*) to retrieve the fraction of binding modes. This shows that PDA can recover the fraction of subpopulations with a deviation of only 0.007 from the expected value (fraction values are normalized to 1) (see Supp. Results for full details). When analyzed by PDA, the asymmetric distribution of binding modes in c2A-celS was confirmed, with a fraction of 0.43 ± 0.02 of the complexes bound in the B1 state (since B1 and B2 fractions add up to 1, only B1 fraction is mentioned). These results are somewhat surprising because the canonical dual binding mode implicitly assumes a symmetrical binding based on both, the sequence conservation among dockerin H1 and H3 helices, and its structural symmetry (Fig 1B and Fig. S3A).

In structures of single binding mode mutants of *C. thermocellum* complexes, the residues at positions 19, 20 and 21 in helix H1 establish a set of important hydrophobic contacts with cohesin in the B1 mode, and the homologous positions 51, 52 and 53 are responsible for these contacts in the B2 mode (*23*). In the case of dockerin celS, sequence conservation between these residues in helix H1 and H3 is poor (Fig. S3A) and could explain the asymmetric distributions. To explore this scenario, we decided to study the dockerin from cellulase A (CelA) which displays a higher degree of similarity in this area (Fig. S3A). Surprisingly, asymmetric binding still occurs and it is even more pronounced (0.37 ± 0.04 of B1 fraction, Fig. 3A). It is noteworthy that these fractions, 0.43 for c2A-celS and 0.37 for c2A-celA are significantly different from the value 0.5 (*p* < 0.05), but also among themselves (*p* < 0.05).

**Figure 3.**
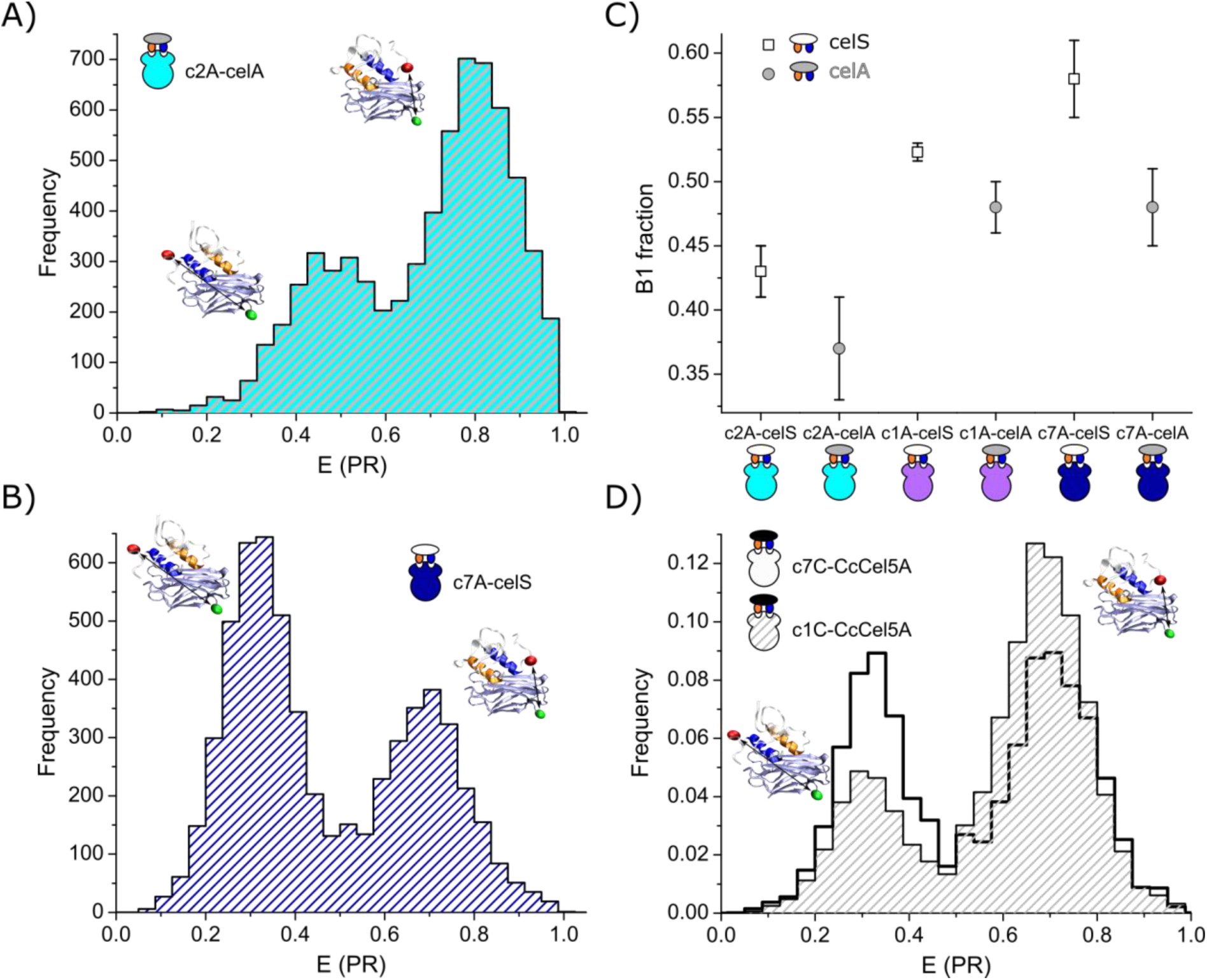
A cohesin-dockerin code. **A)** FRET histogram of the c2A-celA complex from *C. thermocellum*. The subpopulation of binding modes are not equally distributed, with a predominance of the B2 population. PDA analysis further confirmed this conclusion (fraction B1 population of 0.37 ± 0.04). This contrasts with the subpopulation distribution of another *C. thermocellum* complex, c7A-celS, shown in **B)**. Unlike the c2A-celA complex, the B1 mode is more abundant in the c7A-celS complex with a B1 fraction of 0.58 ± 0.03. **C)** Plot summarizing the binding mode distribution of all *C. thermocellum* complexes studied. With the exception of c1A-celA and c7A-celA all complexes display different distribution of binding modes (at p < 0.05). **D)** Comparison of c1C-Ccel5A and c7C-Ccel5A complexes from *C. cellulolyticum*. The difference of binding modes distribution between both complexes is obvious from the E histograms (B1 fraction of c1C-Ccel5A 0.26 ± 0.02 vs 0.45 ± 0.02 of c7C-Ccel5A).

Although crystallographic studies have shown that the network of hydrogen-bonds in both binding modes are be very similar, a few extra hydrogen bonds between the dockerin and cohesin are present in the PDB structure of the B2 mode mutant (*23*). These hydrogen bonds could explain the equilibrium shift toward B2 binding. Specific elimination of these hydrogen bonds without any side effects is virtually impossible, for this reason, we decided to probe this scenario indirectly, by studying the equilibrium population in the highly homologous cohesin 7 module from the same scaffoldin CipA (Fig. S3B). Cohesin 7 (c7A) displays a high degree of similarity with c2A (73% identity and 94% similarity) and, more importantly, shows a total conservation of the residues involved in the cohesin-dockerin interaction (*23, 25*) (Fig. S3B). We reasoned that the same set of hydrogen bonds would be very likely present in c7A and the trend toward B2 binding should remain if these extra hydrogen bonds account for the difference. Unexpectedly, our results suggest that these contacts are not the primary cause of the binding asymmetry, because the histograms of the c7A-celS complex showed an inversion of the subpopulations (Fig. 3B and Fig. S4A), with a prominence of B1 state at equilibrium. PDA analysis of c7A-celS and C7A-celA further confirm this conclusion with a B1 subpopulation fraction of 0.58 ± 0.03 and 0.48 ± 0.03 respectively (Fig. 3C). These changes among cohesin-dockerin complexes point to a scenario where, despite being very similar, different cohesin-dockerin complexes can show distinct distribution of binding modes, a sort of cohesin-dockerin code. This view was further confirmed when we included in our analysis the cohesin 1 module from the same scaffolding unit (c1A-celS and c1A-celA, Fig. 3C and S5). Furthermore, when comparing different cohesin-dockerin combinations, all the complexes display different subpopulation ratios (*p* < 0.05), with the only exception of the pair c1A-celA and c7A-celA (Fig. 3C). The binding modes were assigned in all cases using SB mutants (Fig. S5).

We found a very similar situation in *C. cellulolyticum* when we studied the cohesin-dockerin complex between the cohesin module 7 of scaffoldin CipC and the dockerin CcCel5A (c7C-CcCel5A). c1C and c7C cohesins are almost identical (81% identity and 96 % similarity, Fig. S3C), but the binding mode population distributions are radically different among c7C-CcCel5A and c1C-CcCel5A complexes (Fig. 3D and Fig.S4B), substantiating our finding of a cohesin-dockerin code that is not simply reflected in the immediate binding interface.

It is unlikely that small sequence differences between helices are the main source of asymmetry, as our results with dockerin celS and celA indicate. Besides, the remarkable structural resemblance between cohesins (Fig. S3D) and its high sequence identity (especially in the residues involved in the interaction, Fig. S3A, S3B and S3C) make them unlikely candidates as well. In order to rule out these possibilities in a single experiment, we designed a swapped version of dockerin celS (celS_Swap), in which the helix H1 and H3 was exchanged. If the asymmetry were a direct result of these two factors, one would expect an inversion of the subpopulation fractions. We did not observe such an inversion in any of the three cohesin-celS_Swap complexes studied (Fig. 4). The largest deviation from the expected value was observed for c2A-celS_Swap, where the B1 state takes over most of the population (B1 0.89 ± 0.04 *vs* the 0.57 expected, Fig. 4A). A predominance of the B2 state was expected for c1A-celS_Swap and c7A-celS_Swap, but instead we observed a higher abundance of the B1 population (B1 fraction of 0.59 ± 0.03 and 0.69 ± 0.03 respectively, see Fig. 4B and 4C).

**Figure 4.**
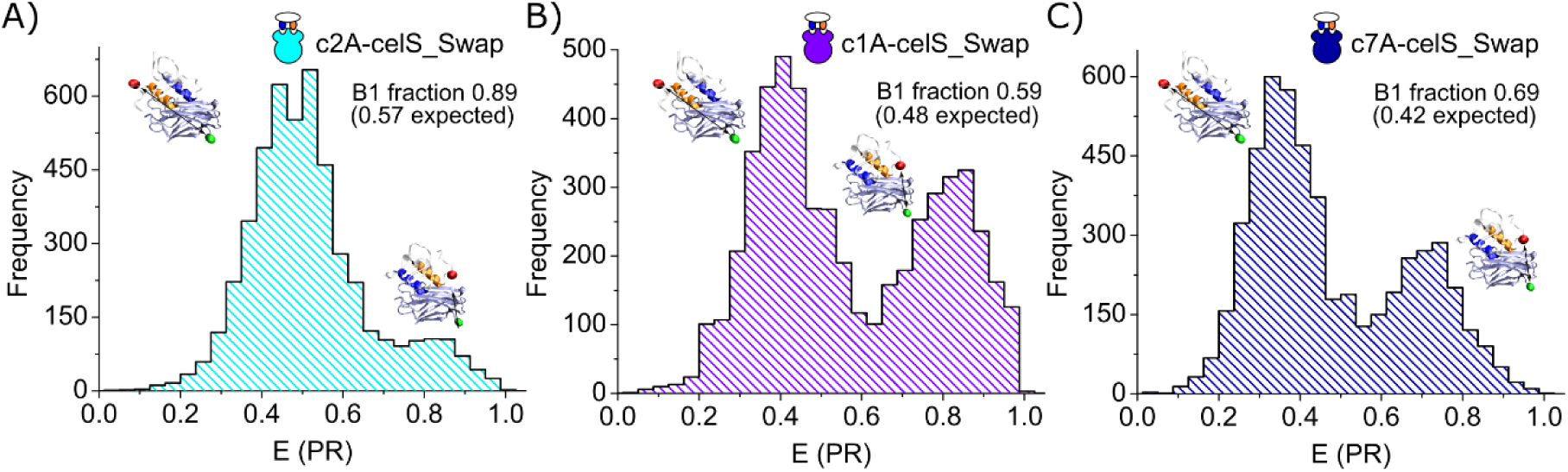
FRET histograms of cohesin-celS_Swap complexes. **A), B)**, and **C)** FRET histograms of c2A-celS_Swap, c1A-celS_Swap and c7A-celS_Swap complexes, respectively. In any of the three cases, the distribution of binding modes matches an inversion of subpopulation fractions relative to wild type complexes. For the c1A-celS_Swap and c7A-celS_Swap complexes a subpopulation inversion would predict a prominence of the B2 mode subpopulation (B1 fraction of 0.48 and 0.42 was expected, respectively), however as it can be seen in B) and C) the B1 mode prevails (B1 fraction of 0.59 ± 0.03 and 0.69 ± 0.03, respectively). In the case of c2A-celS_Swap complex, although a prevalence of B1 over B2 was expected, the deviation from the anticipated value was large (B1 fraction of 0.89 ± 0.04 vs the 0.57 expected).

Overall, our results suggest that the selection of binding mode is globally determined, probably by very small deviations throughout the cohesin-dockerin complex that impact the binding energies and the final populations of the binding modes.

### The dockerin clasp as a global regulator

The so-called dockerin clasp is a structure of unclear function found in several dockerins, and closes the dockerin by connecting its N and C termini (*20, 36*). Formed, in the case of dockerin celS, by the stacking of Tyr5 and Pro66, it is structurally distant from the cohesin-dockerin interface (Fig. 5A) but mutational studies have revealed that it plays a role in the stability of the cohesin-dockerin complex (*36*). In the context of our results, this makes the clasp a potential regulator of the dual binding modes.

**Figure 5.**
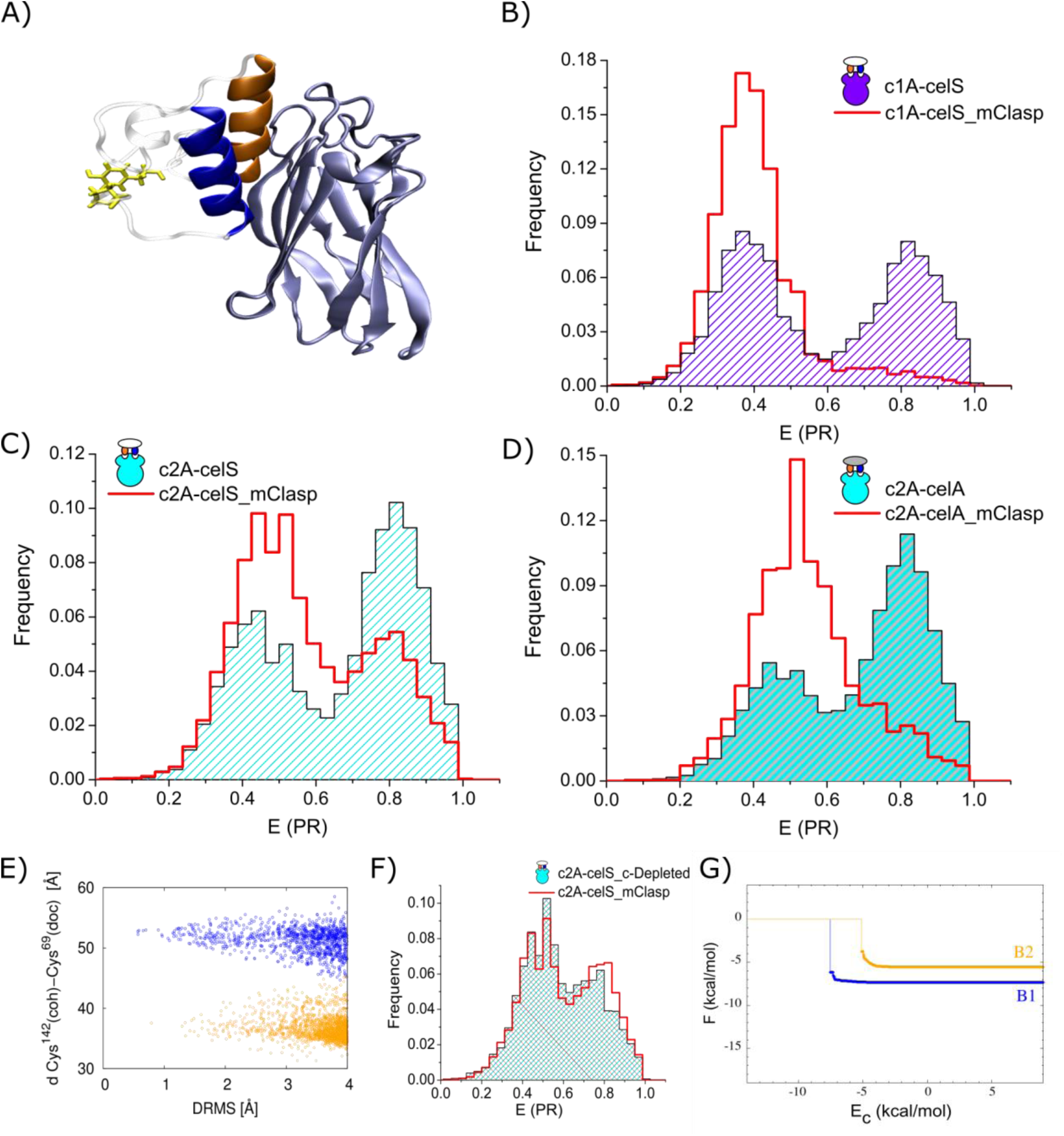
The clasp structure. **A)** A stacking interaction between dockerin residues Tyr5 and Pro66 creates an intramolecular clasp. As can be seen in the representation of c2A-celS complex, the clasp is located far away from the surface of interaction between cohesin and dockerin. **B), C)** and **D)** comparison of c1A-celS, c2A-celS, and c2A-celA wild type complexes with clasp mutant complexes (celS_mClasp in represented in red, wild type complexes in striped-colored histograms). Clasp mutants display a different distribution binding modes subpopulations, with a displacement of the equilibrium towards the B1 state. E) Coarse-grained simulation of the c2A-celS_mClasp complex. The distance between the α-C atoms of CYS^142^ in cohesin and CYS^69^ in dockerin CelS versus DRMS to B1 mode (DRMS(*S, S*_*I*_), blue data points) or B2 mode (DRMS(*S, S*_*II*_), orange data points) is shown. Each of the data points corresponds to a cohesin-dockerin structure obtained from the CG simulations. Unfortunately, the method fails to reproduce the shift toward the B1 mode, as can be notice from the similar distribution of structures in both binding modes (see Fig S7B). **F)** Effect in the dual binding mode of removing the last 6 C-terminal dockerin residues. smFRET histograms display the same shift toward the B1 state on both mutants, c2A-celS_mClasp mutant and c2A-celS_C-Depleted, indicating that the effect observed in the last is mainly due to the suppression of the clasp structure. **G)** Plot of the binding energies obtained with our Fold-X based simulations. Energy of binding in B1 mode (blue line) and B2 mode (orange line) quickly saturated as soon as the E_c_ value is large enough. The plot shows the results obtained for the complex c2A-celS_C-Depleted, which is unable to form the clasp structure. The simulation shows that the energy for binding in the B1 mode is lower by -3.7 kcal/mol for this complex.

In order to study the role of the clasp, we designed a clasp knockout dockerin mutant CelS_mClasp (Y5A, P66A) and we studied the complexes formed with c2A, c1A and c7A cohesins by smFRET. The mutation has a severe effect in all the complexes, especially in the complexes c1A-celS_mClasp and c7A-celS_mClasp where the B2 state was virtually eliminated (Fig. 5B and Fig. S6A). The effect is milder in c2A-celS_mClasp (Fig. 5C), but the shift towards the B1 state is consistent throughout the three complexes. The same shift toward B1 state was observed when we replicated the mutation in dockerin celA (Fig. 5D and S6A). These changes indeed make the clasp an important player in the dual binding mode, and suggest it is a key region contributing to the binding modes’ stability.

### Molecular dynamics simulations suggest that subtle structural differences alter binding mode populations

From a thermodynamic perspective, the shift toward the B1 state displayed by the clasp mutants points to two alternative effects of the clasp in the wild type complex, either a stabilization of the B2 state or a destabilization of B1. In order to get insight into the scenario that is taking place, we utilized two different molecular dynamics simulations approaches. The first involves a Monte Carlo (MC) sampling based on a coarse-grained (CG) model (*37*). The second method uses a previously described FoldX-based approach (*38*), but in a significantly improved manner that also involves a MC sampling (see Methods). Specifically, we applied them to study the c2A-celS complex.

The CG simulations predict the dual binding mode behavior in the wild type complex, with the probability of each mode close to 0.5 (Fig. S7A and S7B). Remarkably, they predict quite well the large change in B1 and B2 states in the case of SB mode mutants (Fig. S7A and S7B). However, they fail to predict the shift toward the B1 state observed experimentally in the clasp mutants. Fig 5E shows that the structures obtained during the MC sampling, assigned either to the B1 (blue dots) or the B2 mode (red dots), are distributed quite homogeneously between both modes, particularly where the density of structures is higher (see values of DRMS between 2-4 Å). In other words, the CG simulations predict that both binding modes are equally probable in the clasp mutant (Fig. S7B). This prediction is actually not unexpected. Since the proteins are represented as rigid bodies during the MC sampling, it is rather unlikely that the CG simulations would correctly capture the effect of charge-conserving mutations (Y5A, P66A) in regions that do not participate directly in the cohesin-dockerin binding. This seems to explain the discrepancy between the simulations and experiments, and also backs the idea that the distribution of binding modes cannot be understood without considering contributions from amino acid residues that are not directly at the binding interface.

On the other hand, the FoldX-based method is an all atom approach where the side chains of the residues are considered explicitly. In the case of the wild type complex, this method predicts an energy difference between B1 and B2 of about -1.4 kcal/mol (supplementary results, Fig. SR6), which indicates a slight preference for B1, but within the range of our error (±2 kcal/mol). For the clasp mutant, this energy difference is larger (−1.8 kcal/mol) but still within the error range (Fig. SR6). Our observations indicate that the method is able to catch the trend observed experimentally, although the energy difference is too small to be considered conclusive. In order to perturb the system toward a situation where our simulation can predict significantly the energy differences, we removed the last six residues from the dokerin’s C-*terminus*. The selection is not accidental, in the first solved structure of a cohesin-dockerin complex (*25*), those residues including Pro66 of the clasp were removed. We hypothesized that the authors were actually able to obtain a crystal because, by removing the proline, clasp formation was prevented and therefore the equilibrium was shifted toward the B1 mode. Indeed, the FRET histogram of this c-depleted mutant recapitulates the shift toward the B1 state observed in the c2A-celS_mClasp mutant (see Fig. 5F), and therefore indicates that the effect originates from the removal of the clasp. The energy difference calculated between the B1 and B2 state for this c-depleted mutant is -3.7 kcal/mol (Fig. 5G), above our error and demonstrating a destabilization of the B2 mode in the mutant. Overall, these results suggest that the changes observed experimentally for the clasp mutant are due to a clasp-mediated stabilization of the B2 state in the wild type complex.

### The clasp as a dual binding mode allosteric regulator

Due to its role in the equilibrium of binding states, it is tempting to speculate about a role of the clasp in the regulation of the dual binding mode. Single amino acid mutants of this structure (either Y5A or P66A mutants) show that Pro66 is key for the response observed in the clasp mutant (Fig. S6B). In the PDB 2MTE, Leu65-Pro66 peptide bond has the particularity of being in the unusual *cis* conformation (*20*). Although the NMR solution structure of the dockerin CelS (2MTE) was solved in the absence of cohesin, our own NMR data show that this bond stays in the *cis* conformation upon cohesin binding (see Supplementary results). Our smFRET data and simulations suggest that the net of contacts of Pro66 and its neighboring residues are involved in the modulation of the cohesin-dockerin interaction, and since this region is well-ordered, a change toward the *trans* conformation would severely affect this net of contacts (*20*).

A particular behavior of *C. cellulolyticum* complexes allowed us to test the previous hypothesis. In contrast to *C. thermocellum* complexes, the samples of the c1C-CcCel5A wild type complex were not at equilibrium at the beginning of the measurement (Fig S8). Instead, we observed slow subpopulation dynamics. Starting from a roughly equally distributed population of binding modes, the system evolved to a situation where the B2 state was the predominant species (Fig. 6A, black symbols and Fig S8). Our data strongly suggest that this process is not due to a direct interconversion between binding modes, but rather a slow re-equilibration process between binding modes that occurs in the diluted conditions of the measurement (see discussion in Fig S9 and S10).

**Figure 6.**
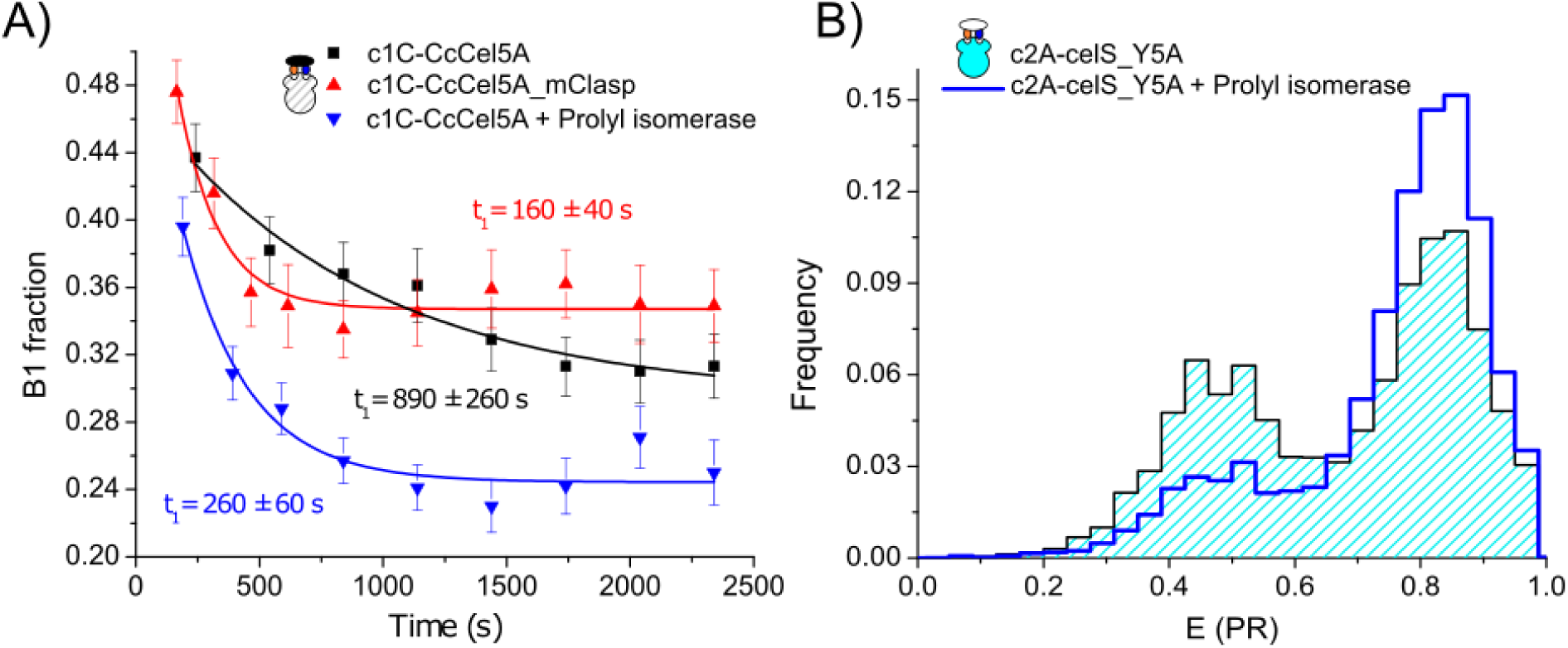
Regulation of dual binding modes by the intramolecular clasp. **A)** Proline *cis*/*trans* isomerization regulates the kinetics towards the steady state. The *C. cellulolyticum* c1C-cCCel5A complex (black symbols) shows a slow kinetics in the distribution of binding modes, from a similarly distributed B1 and B2 states at t_0_, the system evolves to a B2-dominated scenario. c1C-CcCel5A-mClasp mutant (red symbols) shows accelerated kinetics towards the steady state, indicating a regulatory role of the clasp in the process. Actually, alteration of the isomerization state of the clasp’s proline by a prolyl isomerase (cyclophilin A) accelerates the kinetics in a similar way as in the clasp mutant (blue symbol). t_1_ stands or decay times of the kinetic process. **B)** Prolyl isomerase activity is able to change the distribution of binding modes in *C. thermocellum* complexes. In blue, the histogram of a cyclophilin-treated c2A-celS_Y5A complex shows the change in the distribution of binding modes when compared to an untreated sample (striped histogram). An increase in the B2 mode population is observed for the treated sample.

Although the clasp mutant of *C. cellulolyticum* c1C-CcCel5A complex shows only mild enrichment in the B1 state when compared to wild type complexes (Fig. S8), the dynamic process driving towards a B2-dominated population is still present, and remarkably the kinetics of this process is greatly accelerated (Fig. 6A, red symbols). Furthermore, if the clasp’s proline in CcCel5A dockerin shares the same isomerization bias as in CelS, we hypothesized that a change in the isomerization state of the clasp’s proline should have a similar effect on the kinetics, as most of the contacts established by the proline in the clasp will be affected. With this idea in mind, we treated the c1C-CcCel5A samples with cyclophilin A, a prolyl isomerase from *E.coli* (*39*) and observed faster kinetics toward the final equilibrium conditions, almost comparable to that found in the c1C-CcCel5A_mClasp complex (Fig. 8A, blue symbols). Furthermore, this effect was not observed when we used a cyclophilin A mutant with hampered isomerase activity that still binds substrate with high affinity (*40*) (Fig. S11A). It is worth highlighting that, besides the accelerated kinetics during the binding modes equilibration, a general destabilization of the complex occurred upon proline isomerization. This is suggested from the accelerated decay over time in the total number of cohesin-dockerin complexes observed in cyclophilin-treated samples which is not observed in the wild type (Fig. S10A and S10B).

In the case of *C. thermocellum* complexes, the effect observed for the clasp mutant complexes led us to expect an effect on the subpopulation fraction upon P66 isomerization. Even though a small enrichment in B2 state was observed for cyclophilin-treated c2A-celS samples, the effect was too mild to be conclusive (Fig S11C). In order to obtain a larger response, we used the single residue clasp’s mutant Y5A (c2A-celS_Y5A), which displays the same subpopulation ratios as the wild type complex (Fig. 2A and S6B). We predicted that the suppression of contacts between the Pro66 and the Tyr5 would weaken the network of contacts that keep the former in the *cis* conformation and therefore would make it more accessible to enzymatic isomerization. The response to enzymatic isomerization was drastically greater for the c2A-celS_Y5A complex (Fig. 6B), with a significant shift (*p* < 0.01) of the population toward the B2 mode (B1 fraction of 0.420 ± 0.016 for the untreated samples *vs* 0.20 ± 0.01 for the cyclophilin-treated). Again, cyclophilin A mutant did not produce such a shift (Fig. S11B). We noticed that this shift toward the B2 state is also observed for cyclophilin-treated c1C-CcCel5A complexes (see Fig. 6A, black *vs* blue symbols).

## Discussion

Although previous detection of dual binding modes using single-molecule force spectroscopy (*26*) had been recently questioned (*41*), our present work undoubtedly demonstrates that type-I cohesin-dockerin complexes bind in two alternative binding modes. Our smFRET approach makes the identification of the two binding modes trivial, mere visual inspection of the FRET efficiency histograms already suffices to identify both binding modes. Most importantly, it opens the door to quantitatively study cellulosome assembly and dynamics at an unprecedented detail.

Interestingly, most cohesin-dockerin complexes show unequal population of the two binding modes. This contrasts with the classical idea of a dual binding mode for cohesin-dockerin complexes, since the internal symmetry of the dockerin and the high degree of sequence similarity between Helix 1 and 3 suggest identical binding modes. Furthermore, the actual distribution of binding modes depends on both the cohesin and the dockerin. Although the structure of whole cellulosomes remains unknown, this cohesin-dockerin code can provide hints on how cellulosomes are structurally arranged, as the code predicts that particular cellulosome conformations are more likely to occur. But, on the other hand, the existence of dual binding modes provides the system with enough structural flexibility to accommodate enzymes in the alternative binding mode in a sterically-constrained environment.

The cohesin-dockerin code, the helices-swapped mutants, and our coarse-grained simulations suggest that although the dual binding mode phenomenon is grounded on the dockerin’s helical symmetry, the final distribution of binding modes is determined globally, with small structural differences throughout the cohesin-dockerin complex contributing to the final binding energies. In this regard, our results with the clasp show that the contributions of the well-ordered residues around the dockerin’s Leu65-Pro66 bond are particularly important. In addition to the clasp’s stacking interaction between Pro66 and Tyr5, Tyr67 is involved in hydrophobic interactions with Ile53 and Ile62 and a hydrogen bond with Leu40. Besides, Leu65 packs on Tyr57’s aromatic ring and a salt bridge between Glu42 and Lys68 is observed (*20*). These residues are expected to be important actors in the allosteric effect observed, since Ile53 and Tyr57 are part of dockerin’s Helix 3, and Leu40 establishes contacts with Asp50 and Leu54, and Ile62 with Tyr57 and Arg56, all of them part of the Helix 3.

Remarkably, we have found that the dockerin intramolecular clasp, a structure without a clear function to date, plays an active role on the regulation of the dual binding mode regulating the equilibrium population of the binding modes, the kinetics toward the final distribution of binding modes and even the global stability of the interaction. Most importantly, we have shown that all the effects of the clasp can be enzymatically regulated by a prolyl-isomerase catalyzing the *cis*/*trans* isomerization of its constitutive proline. Overall, our results have great implications on cellulosome physiology as they provide a mechanistic insight into their structural regulation and organization. The regulation of the distribution of binding modes is a game changer in our understanding of the cellulosome plasticity, since we shift from a paradigm of passive elements that generate plasticity, to a view in which the plasticity can be actively actuated and regulated. Furthermore, the regulated-kinetics toward the steady state can be exploited to reach a particular cellulosome configuration more quickly, leveraging efficiency in a particular substrate micro-environment. Besides, the general complex destabilization upon proline isomerization provides a simple mechanism to achieve cellulosome remodeling (by faster exchange of cellulosomal enzymes). Although we used a prolyl isomerase from *E. coli*, a Blast search (https://blast.ncbi.nlm.nih.gov/Blast.cgi) in *C. thermocellum* and *C. cellulolyticum* genomes retrieves a match with a predicted prolyl isomerase in both cases. Interestingly, the sequences display a signal peptide for extracytoplasmic export (GenBank: ABN51309.1 and ACL77371.1) and in the case of *C. thermocellum*, an upregulation of this gene has been experimentally observed during the stationary phase (*29*). Future studies should address the identification and characterization of prolyl isomerases from cellulosome-producing species, as well as their activity in cohesin-dockerin complexes.

Besides, our findings are not only relevant to cohesin-dockerin type-I complexes of *Clostridium* species. Clasp structures have been found also in type-II complexes, and in several other cellulosome-producing species as well (*20, 36*). Furthermore, the chemical nature of the clasp is completely different in some cases, featuring electrostatic interaction between charged residues (*36*). It is tempting to speculate that this salt bridge-based clasp could confer a simple mechanism for cellulosomes remodeling upon pH changes, since acidification takes place during the fermentation carried out by these organisms (*42*).

Finally, our results open the gate to the incorporation of dynamics features for the design of improved designer cellulosomes, which are a promising and affordable alternative for bioethanol production from lignocellulosic waste (*10*).

## Materials and methods

A full description of methods and materials is provided in the supplementary material section.

## Acknowledgements

This project has received funding from the European Union’s Horizon 2020 research and innovation programme under the Marie Skłodowska-Curie grant agreement No 746635. Support by the DFG excellence cluster CIPSM is acknowledged. This work was supported by a Seventh Framework Programme in Nanosciences, Nanotechnologies, Materials & New Production Technologies (7PM - NMP 2013-17, 604530-2, *CellulosomePlus*) and the ERA-IB-ERANET-2013-16 (EIB.12.022, *FiberFuel*) through the Spanish MINECO (PCIN-2013-011-C02-01) to MCV and by a SAF2016-76678-C2-2-R grant (DVL). NMR experiments were performed in the “Manuel Rico” NMR laboratory (LMR) of the Spanish National Research Council (CSIC), a node of the Spanish Large-Scale National Facility (ICTS R-LRB). The Warsaw group has received support from the National Science Centre (NCN) under grant Nos. 2018/31/B/NZ1/00047 (MC and MW) and 2016/21/B/NZ1/00006 (BR and MC).

AMV thanks Angelika Kardinal for her assistance and support in all the molecular biology work, and to Javier Oroz and Meltem Aygüler for critical reading of the manuscript.

## Author contributions

AMV designed the research, performed the smFRET experiments, analyzed the data and wrote the paper. AG-P and DVL performed the NMR experiments, analyzed the data and wrote the paper. MW, BR and MC performed the molecular dynamics simulations and wrote the paper. MC-V and PT wrote the paper

## References

1. S. P. Smith, E. A. Bayer, Insights into cellulosome assembly and dynamics: from dissection to reconstruction of the supramolecular enzyme complex. Current Opinion in Structural Biology 23, 686–694 (2013).

2. E. A. Bayer, J. P. Belaich, Y. Shoham, R. Lamed, The cellulosomes: multienzyme machines for degradation of plant cell wall polysaccharides. Annu Rev Microbiol 58, 521–554 (2004).

3. C. M. Fontes, H. J. Gilbert, Cellulosomes: highly efficient nanomachines designed to deconstruct plant cell wall complex carbohydrates. Annu Rev Biochem 79, 655–681 (2010).

4. L. Artzi, E. A. Bayer, S. Morais, Cellulosomes: bacterial nanomachines for dismantling plant polysaccharides. Nat Rev Microbiol 15, 83–95 (2017).

5. R. Lamed, E. Setter, E. A. Bayer, Characterization of a cellulose-binding, cellulase-containing complex in Clostridium thermocellum. J Bacteriol 156, 828–836 (1983).

6. M. A. Currie et al., Scaffoldin conformation and dynamics revealed by a ternary complex from the Clostridium thermocellum cellulosome. The Journal of biological chemistry 287, 26953–26961 (2012).

7. J. L. A. Brás et al., Diverse specificity of cellulosome attachment to the bacterial cell surface. Scientific Reports 6, 38292 (2016).

8. S. Yaron, E. Morag, E. A. Bayer, R. Lamed, Y. Shoham, Expression, purification and subunit-binding properties of cohesins 2 and 3 of the Clostridium thermocellum cellulosome. FEBS Letters 360, 121–124 (1995).

9. Y. Vazana et al., A synthetic biology approach for evaluating the functional contribution of designer cellulosome components to deconstruction of cellulosic substrates. Biotechnology for Biofuels 6, 182 (2013).

10. J. Stern, S. Moraïs, R. Lamed, E. A. Bayer, Adaptor Scaffoldins: An Original Strategy for Extended Designer Cellulosomes, Inspired from Nature. mBio 7, e00083–00016 (2016).

11. F. Mingardon, A. Chanal, C. Tardif, E. A. Bayer, H.-P. Fierobe, Exploration of New Geometries in Cellulosome-Like Chimeras. Appl Environ Microbiol 73, 7138 (2007).

12. Y. J. Bomble et al., Modeling the Self-assembly of the Cellulosome Enzyme Complex. Journal of Biological Chemistry 286, 5614–5623 (2011).

13. M. Hammel et al., Structural basis of cellulosome efficiency explored by small angle X-ray scattering. J Biol Chem 280, 38562–38568 (2005).

14. A. L. Molinier et al., Synergy, structure and conformational flexibility of hybrid cellulosomes displaying various inter-cohesins linkers. J Mol Biol 405, 143–157 (2011).

15. A. Barth et al., Dynamic interactions of type I cohesin modules fine-tune the structure of the cellulosome of <em>Clostridium thermocellum</em>. Proceedings of the National Academy of Sciences 115, E11274–E11283 (2018).

16. R. Borne, E. A. Bayer, S. Pages, S. Perret, H. P. Fierobe, Unraveling enzyme discrimination during cellulosome assembly independent of cohesin-dockerin affinity. Febs j 280, 5764–5779 (2013).

17. B. Garcia-Alvarez et al., Molecular architecture and structural transitions of a Clostridium thermocellum mini-cellulosome. J Mol Biol 407, 571–580 (2011).

18. K. Hirano et al., Enzymatic diversity of the Clostridium thermocellum cellulosome is crucial for the degradation of crystalline cellulose and plant biomass. Scientific reports 6, 35709–35709 (2016).

19. S. Moraïs et al., Deconstruction of Lignocellulose into Soluble Sugars by Native and Designer Cellulosomes. mBio 3, e00508–00512 (2012).

20. C. Chen et al., Revisiting the NMR solution structure of the Cel48S type-I dockerin module from Clostridium thermocellum reveals a cohesin-primed conformation. J Struct Biol 188, 188–193 (2014).

21. F. Schaeffer et al., Duplicated Dockerin Subdomains of Clostridium thermocellum Endoglucanase CelD Bind to a Cohesin Domain of the Scaffolding Protein CipA with Distinct Thermodynamic Parameters and a Negative Cooperativity. Biochemistry 41, 2106–2114 (2002).

22. A. Karpol, Y. Barak, R. Lamed, Y. Shoham, E. A. Bayer, Functional asymmetry in cohesin binding belies inherent symmetry of the dockerin module: insight into cellulosome assembly revealed by systematic mutagenesis. Biochem J 410, 331–338 (2008).

23. A. L. Carvalho et al., Evidence for a dual binding mode of dockerin modules to cohesins. Proceedings of the National Academy of Sciences 104, 3089 (2007).

24. B. A. Pinheiro et al., The Clostridium cellulolyticum dockerin displays a dual binding mode for its cohesin partner. J Biol Chem 283, 18422–18430 (2008).

25. A. L. Carvalho et al., Cellulosome assembly revealed by the crystal structure of the cohesin– dockerin complex. Proceedings of the National Academy of Sciences 100, 13809 (2003).

26. M. A. Jobst et al., Resolving dual binding conformations of cellulosome cohesin-dockerin complexes using single-molecule force spectroscopy. eLife 4, e10319 (2015).

27. M. A. Nash, S. P. Smith, C. M. Fontes, E. A. Bayer, Single versus dual-binding conformations in cellulosomal cohesin-dockerin complexes. Curr Opin Struct Biol 40, 89–96 (2016).

28. Y. Nataf et al., Clostridium thermocellum cellulosomal genes are regulated by extracytoplasmic polysaccharides via alternative sigma factors. Proc Natl Acad Sci U S A 107, 18646–18651 (2010).

29. B. Raman, C. K. McKeown, M. Rodriguez, Jr., S. D. Brown, J. R. Mielenz, Transcriptomic analysis of Clostridium thermocellum ATCC 27405 cellulose fermentation. BMC Microbiol 11, 134 (2011).

30. S. Yoav et al., How does cellulosome composition influence deconstruction of lignocellulosic substrates in Clostridium (Ruminiclostridium) thermocellum DSM 1313? Biotechnology for biofuels 10, 222–222 (2017).

31. E. Sisamakis, A. Valeri, S. Kalinin, P. J. Rothwell, C. A. Seidel, Accurate single-molecule FRET studies using multiparameter fluorescence detection. Methods Enzymol 475, 455–514 (2010).

32. B. Hellenkamp et al., Precision and accuracy of single-molecule FRET measurements—a multi-laboratory benchmark study. Nature Methods 15, 669–676 (2018).

33. M. Antonik, S. Felekyan, A. Gaiduk, C. A. Seidel, Separating structural heterogeneities from stochastic variations in fluorescence resonance energy transfer distributions via photon distribution analysis. J Phys Chem B 110, 6970–6978 (2006).

34. S. Kalinin, S. Felekyan, A. Valeri, C. A. M. Seidel, Characterizing Multiple Molecular States in Single-Molecule Multiparameter Fluorescence Detection by Probability Distribution Analysis. The Journal of Physical Chemistry B 112, 8361–8374 (2008).

35. W. Schrimpf, A. Barth, J. Hendrix, D. C. Lamb, PAM: A Framework for Integrated Analysis of Imaging, Single-Molecule, and Ensemble Fluorescence Data. Biophysical Journal 114, 1518–1528 (2018).

36. M. Slutzki et al., Intramolecular clasp of the cellulosomal Ruminococcus flavefaciens ScaA dockerin module confers structural stability. FEBS Open Bio 3, 398–405 (2013).

37. Y. C. Kim, G. Hummer, Coarse-grained models for simulations of multiprotein complexes: application to ubiquitin binding. J Mol Biol 375, 1416–1433 (2008).

38. M. Wojciechowski et al., Dual binding in cohesin-dockerin complexes: the energy landscape and the role of short, terminal segments of the dockerin module. Scientific Reports 8, 5051 (2018).

39. J. Liu, C. T. Walsh, Peptidyl-prolyl cis-trans-isomerase from Escherichia coli: a periplasmic homolog of cyclophilin that is not inhibited by cyclosporin A. Proc Natl Acad Sci U S A 87, 4028–4032 (1990).

40. C. Camilloni et al., Cyclophilin A catalyzes proline isomerization by an electrostatic handle mechanism. Proceedings of the National Academy of Sciences 111, 10203–10208 (2014).

41. M. Wojciechowski, M. Cieplak, Dual binding mode in cohesin-dockerin complexes as assessed through stretching studies. The Journal of Chemical Physics 145, 134102 (2016).

42. P. J. Weimer, J. G. Zeikus, Fermentation of cellulose and cellobiose by Clostridium thermocellum in the absence of Methanobacterium thermoautotrophicum. Appl Environ Microbiol 33, 289–297 (1977).

