## Supplementary information for "Cohesin-dockerin code in cellulosomal dual binding modes and its allosteric regulation by proline isomerization"

#### *Materials and Methods*

##### Protein design and cloning

All constructs used in the present work were produced by PCR cloning or direct subcloning using standard cloning techniques(*1*). When PCR-cloned, genes were amplified from existing plasmids or genomic DNA; in addition, several genes were commercially purchased from gene synthesis services (Eurofins Genomics, Germany). Purchased genes were codon-optimized for expression in *Escherichia coli* using a web server (http://genomes.urv.es/OPTIMIZER/). For a detailed list of the used PCR templates, plasmids and restriction enzymes see Table 1. Point mutations were introduced in the cloning oligos, since most of the mutated residues were located near the N- or C-terminus of the protein fragment. When mutations were located somewhere in the middle of the gene, overlapping PCR was used to introduce the mutation. All dockerins were fused to the C-terminus of a cysteine-free xylanase 6 from *Bacillus stearothermophilus*. Xylanase 6 has been used previously for dockerin production since it provides both, high solubility and protein yield(*2*). The plasmid XynT6-DocS carrying the xylanase 6 from *B. stearothermophilus* fused to the dockerin celS from *C. thermocellum* was used as a template and PCR subcloned between the sites NcoI and XhoI in the vector pET 24d (Novagen, Merck Millipore, Darmstadt, Germany). There is a BamHI restriction site between the xylanase 6 and the dockerin that was used to exchange the different dockerin genes in a cassette fashion. This pET 24-d derived plasmid will be called pXyn from now on. The plasmid XynT6-DocS was a generous gift of Prof. Edward Bayer (Weizmann Institute, Israel). Plasmid encoding for hetero-polyproteins, described elsewhere (*3*), were used for PCR cloning of cohesin c2A, c7A and c1C.

For NMR studies, the dockerin sequence was fused to xyn6 and TEV cleavage site was introduced in between. The construct was generated by independently amplifying xyn6 (adding NheI site at 5' and TEV restriction site and part of the sequence from 5' celS dockerin) and dockerin celS (adding 2 stop codons and XhoI restriction site at its 3'). The fragments were then fused using overlapping PCR and the product cloned into pET28 vector (Novagen, Merck Millipore, Darmstadt, Germany) using NheI and XhoI sites. The final construct consisted of His6-Xyn6-TEV-DockCelS.

Point mutations to cysteine have been introduced in the proteins in order to dye-label them for FRET measurements. The position of the cysteines were chosen in order to maximize the difference of the distance between the dyes in the both binding modes while minimizing the impact on the proteins and their interaction. As for the cohesin modules, cysteines were introduced at the C-terminus just after the folded domain and at the beginning of the natural linker between cohesins. Cysteine was introduced in the position Asp63 of dockerin CelS since it is a poor conserved position next to the dockerin C-terminus (*4*), and its side chain does not participate in any contacts. The analogous position to Asp63 was selected for the rest of dockerins studied (see protein list below).

In the case of *C. thermocellum* dockerins, four point mutations were introduced in order to generate single binding mode mutants (SB). CelS dockerin residues S16, T17, K23 and R24 in H1 and their corresponding residues in H3 (S48, T49, 55K, 56R) are expected to sustain an extensive net of contacts with the cohesin in B1 or B2 mode respectively (*5, 6*). We used here a tetra-mutant of helix 1 (S16D, T17E, K23A and R24A) with severe impairment in B2 interaction that we called celS_SB1 (single binding mode 1). CelS_SB2 mutant carries analogous mutations that hamper interaction in the B1 mode(*2*). In addition, we generated a second version of single binding mode mutant 2 (SB2.1) in which only two mutations where made (S48A, T49A). The same kind of mutant was used by the crystallographic study that revealed the first structure of a *C. thermocellum* cohesin-dockerin complex in the B2 state. We used CelS_B2.1 to generate the cohesin 1 artificial data set analyzed by PDA (see Sup. Results). CelA_SB1 and CelA_SB2 carry the same kind of quadruple mutations described for CelS dockerin (see protein list below).

*C. cellulolyticum* SB mutants implement the same two point mutations used in(*5*) to generate cohesin-dockerin complexes that interact either in the SB1 or the SB2. (see protein list below)

*C. thermocellum* CipA cohesin 5 clone for NMR studies was described previously (*7*)

| Construct | Gene origin | Restriction Sites | Expression vector |
|---|---|---|---|
| Cel S | PCR amplified from XynT6-DocS plasmid | NcoI/XhoI | pXyn |
| Cel A | PCR amplified from *C. thermocellum* genomic DNA | BamHI/XhoI | pXyn |
| CcCel5A | PCR amplified from genomic DNA | BamHI/XhoI | pXyn |
| CelS_mClasp | PCR amplified from plasmid | BamHI/XhoI | pXyn |
| CelS_C-depleted | PCR amplified from plasmid | BamHI/XhoI | pXyn |
| CelS_5YA | PCR amplified from XynT6-DocS plasmid | BamHI/XhoI | pXyn |
| CelS_P66A | PCR amplified from XynT6-DocS plasmid | BamHI/XhoI | pXyn |
| CelS_Swap | Gene synthesis services | BamHI/XhoI | pXyn |
| CelS SB1 mutant | PCR amplified from XynT6-DocS plasmid | BamHI/XhoI | pXyn |
| CelS SB2 mutant | PCR amplified from XynT6-DocS plasmid | BamHI/XhoI | pXyn |
| CelS SB2.1 mutant | PCR amplified from XynT6-DocS plasmid | BamHI/XhoI | pXyn |
| CelA_mClasp | PCR amplified from *C. thermocellum* genomic DNA | BamHI/XhoI | pXyn |
| CelA SB1 mutant | PCR amplified from *C. thermocellum* genomic | BamHI/XhoI | pXyn |

| | | | |
|---|---|---|---|
| | DNA | | |
| CelA SB2 mutant | PCR amplified from *C. thermocellum* genomic DNA | BamHI/XhoI | pXyn |
| Xyn10B | Gene synthesis services | BamHI/XhoI | pXyn |
| c2A | PCR cloning from plasmid | NcoI/XhoI | pET 24-d |
| AviTag-c2A | PCR cloning from plasmid | NcoI/XhoI | pET 24-d |
| c1A | Gene synthesis services | NcoI/XhoI | pET 24-d |
| c7A | PCR cloning from plasmid | NcoI/XhoI | pET 24-d |
| c1C | PCR cloning from plasmid | NcoI/XhoI | pET 24-d |
| AviTag-c1C | PCR cloning from plasmid | NcoI/XhoI | pET 24-d |
| c7C | Gene synthesis services | NcoI/XhoI | pET 24-d |
| Cyclosphilin A | PCR amplified from *E. coli* genomic DNA | NdeI/XhoI | pET 21-a |
| Cyclophilin A R50A mutant | PCR amplified from *E. coli* genomic DNA | NdeI/XhoI | pET 21-a |
| Bir A | PCR amplified from *E. coli* genomic DNA | NcoI/XhoI | pET 24-d |
| Xyn6-TEV-CelS | PCR cloning from plasmids | NheI/XhoI | pET 28 |

**Table 1**. List of protein constructs used in this work. The specific plasmid used for PCR amplification are described in the text. Please note that celS, celA, CcCel5A and Xyn10A refers to the dockerin of these enzymes, as described in the main text.

**Protein production and purification.**

*E. coli* BL21 Star strain was used for recombinant protein production. Bacterial cultures were grown in LB broth at 37 °C with vigorous agitation and protein production was induced at an OD 0.8-1.2 by addition of 0.5 mM IPTG. Expression took place at 16 °C overnight for cohesin constructs and for 4 hours at 25 °C for the rest of the samples. Cell pellets were resuspended in 50 mM phosphate buffer pH 7.4, 500 mM NaCl, 50 mM Imidazole (Buffer A), and lysed by chemical methods. Proteins were purified by $Ni^{2+}$ -affinity chromatography using Histrap HP columns (GE Healthcare, IL, USA) on an FPLC apparatus (ÄKTA Purifier, GE Healthcare) and eluted from the column with a 500mM imidazole containing buffer (buffer B). When necessary, a second purification step by size exclusion chromatography was performed (Superdex 200 10/300 GL, GE Healthcare). The purity of proteins was monitored by SDS-PAGE and the molar concentration estimated spectrophotometrically using the theoretical extinction coefficient at 280 nm.

In the case of cohesin 5 construct used for NMR studies, cells were grown in LB media at 37 °C and then expression was induced for 16 h at 16 °C by addition of 0.1 mM IPTG. $^{15}N$ and $^{13}C$ labelled dockerin was obtained following a method described elsewhere (*8*). Briefly, cells were grown in LB media until $OD_{595\ nm}$ ~ 0.6, then they were harvested by centrifugation and resuspended in M9 media, collected again by centrifugation and resuspended in one fourth of original volume of M9 media containing $^{15}NH_4Cl$ and $^{13}C$-glucose and 1 mM $CaCl_2$. Cells were incubated at 16 °C for 1 h and then expression was induced for 16 h by adding 1 mM IPTG. Cell pellets were resuspended and lysated in 50 mM Tris pH 7.5, 500 mM NaCl, 1mM $CaCl_2$, 50 mM Imidazole and 1 mM DTT. Proteins were purified using $Ni^{2+}$ NTA columns and eluted in 500mM imidazole containing buffer. Fractions containing protein were then buffer exchanged to 50 mM Tris, 300 mM NaCl, 1 mM $CaCl_2$, 3 mM DTT and cleaved using TEV protease overnight at 37 °C and re-purified by reverse IMAC. Clean proteins were concentrated and buffer exchanged to ammonium bicarbonate pH7.5, frozen in liquid nitrogen and lyophilized overnight.

**Protein labelling with fluorescent dyes**

In order to achieve site-specific labeling, cysteine residues were introduced in the proteins (see protein list) and maleimide-modified dyes used for labeling. The labeling reaction takes place in 20 mM MOPS, 100 mM NaCl, 0.2 mM EDTA pH 7, and a pre-incubation step with 5 mM TCEP (30 min at RT) was performed in order to reduce the cysteine sulfhydryl groups. Usually, a 20 fold excess of maleimide-dye was used, typical reaction conditions were 50 µM of protein *versus* 1 mM of maleimide-dye for 90 min, at RT, in light-tight 1.5 ml tubes. Dye excess was removed using a 1 ml FF affinity Histrap column (GE Healthcare Life Sciences, IL, USA). Protein-dye solution was manually loaded in the column and washed with at least 10 volumes of Buffer A. Finally, protein was eluted from the column with buffer B and buffer exchanged to 20 mM Tris pH 7.4, 100 mM NaCl, 1 mM $CaCl_2$ using Amicon Ultra Centrifugal Filters (Merck Millipore, Darmstadt, Germany).

**Biotinylation of recombinant proteins**

Cohesin modules carrying an N-terminal Avitag peptide sequence were enzymatically biotinylated using the Bir A enzyme from *E.coli* (*9, 10*). Biotinylation was carried out at 30 °C for 1 hour in 50 mM Tris pH 8.4 under the following conditions: 30 µM Avitag-protein, 0.9 µM Bir A enzyme, 1.4 mM ATP, 3.5 mM $MgCl_2$, 105 µM D-Biotin. Prior to large scale biotinylation, small reactions were performed and the degree of modification checked by SDS-PAGE retardation assay upon streptavidin binding as described elsewhere (*11*).

After biotinylation, proteins were further purified by size exclusion chromatography in order to separate the biotinylated protein from the Bir A enzyme and free biotin molecules.

**Labtek chambers passivation and functionalization**

For solution measurements, Labtek chambers (Lab-Tek II chambered coverglass sytem, ThermoFisher Scientific, MA, USA) were passivated with 1 mg/ml BSA (PAA laboratories GmbH, Germany). BSA solution was kept in the chamber for at least 1 hour and only removed just before the chamber was used. After removal of the BSA solution, chamber was washed out at least 3 times with PBS and finally with measurement buffer (50 mM Tris, 100 mM NaCl, 2mM Trolox/Trolox quinone, 1% glucose, 1 mM $CaCl_2$, pH 8).

In the case of measurements with surface-immobilized molecules, chambers were first etched for 3 hours in 1 M KOH followed by 3 washing steps with PBS buffer. Chambers were functionalized with a solution of 0.5 mg/ml biotin-labeled Albumin (Merck KGaA, Darmstadt, Germany) overnight at 4°C. After incubation, biotinylated albumin was washed out and the chamber incubated with 0.25 mg/ml Neutravidin (ThermoFisher Scientific) solution in PBS for 15 min. Finally, the chamber was washed 3 times with PBS.

**smFRET experiments**

Confocal fluorescence measurements on surface-immobilized molecules were performed using a home-built confocal setup based on an Olympus IX-83 inverted microscope and a 78 MHz-pulsed laser beam (SuperK Extreme, NKT Photonics). Wavelength selection was achieved *via* an acousto-optically tunable filter (AOTF, SuperK Dual AOTF, NKT Photonics, Denmark) and a digital controller (AODS 20160 8R, Crystal Technology, Inc., USA). *Via* computer software (AODS 20160 Control Panel, Crystal Technology, Inc. USA) selection of wavelegths in a range between 400 and 700nm is possible. For ALEX-FRET measurements(*12*) wavelengths of 532nm and 640nm are chosen to excite the dyes. A second AOTF (AA.AOTF.ns: TN, AA-Opto-Electronic, France) is used to alternate between the two wavelengths. Furthermore, the second AOTF, controlled *via* LabVIEW software, is used to control laser intensity as well as spectrally cleaning the laser beam. A neutral density filter (ndF, OD 0-2, Thorlabs, Germany) is used to regulate the laser intensity manually. 1 µW and 3 µW of excitation power are used for the green and red laser (measured in front of the entrance of the microscope). The height difference between excitation path and microscope body is overcome by coupling the laser into a polarization maintaining fiber (PM-Faser, P1-488PM-FC-2, Thorlabs, Germany). The laser is then coupled into a linear polarizer (LPVISE100-A, Thorlabs, Germany), followed by a quarter-wave plate (λ/4, AQWP05M-600, Thorlabs, Germany). The laser is focused onto the sample with an oil-immersion objective (UPlanSApo100x/1.4 NA, WD=0:12 mm, Olympus). Positioning of the sample is performed with a piezo stage (E-501.00, Physik Instrumente GmbH & Co. KG). Excitation light is separated from the emitted light through the dichroic beam splitter and then focused on a 50 µm pinhole (Linos). The emission channels for red and green are spectrally filtered (red: Razo-rEdge 647, Semrock, USA and green: Brightline HC582/75, AHF, Germany). The light is then detected by a Single-Photon Avalanche Diode (SPCM, AQR 14, PerkinElmer, MA, United States) and registered by a TCSPC system (HydraHarp 400, PicoQuant GmbH, Berlin, Germany). A custom-made LabVIEW software (National Instruments) is used to process the acquired data.

Cohesin-dockerin samples were incubated for a few seconds at a concentration of 2 µM, at a cohesin:dockerin molar ratio of 1:1 for c2A-celS and 1:2 for c1C-CcCel5A samples. Samples were diluted to a concentration of $\sim$ 20 pM in 20 mM Tris, 100 mM NaCl, 1 mM $CaCl_2$, pH 8 and incubated for 5-10 minutes in neutravidin-functionalized chambers. The solution was then removed and the chamber washed out once with the same buffer. Measurements were carried out in sealed chambers in 50 mM

Tris, 100 mM NaCl, 2mM Trolox/Trolox quinone, 1% glucose, 1 mM $CaCl_2$, pH 8, supplemented with 100 U/mL glucose oxidase, and 160 U/mL catalase for oxygen removal (*13*).

Solution smFRET experiments were performed on a PIE-based(*14*) home built confocal microscope based on an Olympus IX-71 inverted microscope. Two pulsed lasers (639 nm, 80 MHz, LDH-D-C-640; 532 nm, 80 MHz, LDH-P- FA-530B; both PicoQuant GmbH) were altered on the nanosecond timescale by a multichannel picosecond diode laser driver (PDL 828 “Sepia II”, PicoQuant GmbH, Berlin, Germany) with an oscillator module (SOM 828, PicoQuant GmbH). The lasers were coupled into a single mode fiber (P3-488PM-FC, Thorlabs GmbH, Dachau, Germany) to obtain a Gaussian beam profile. Circular polarized light was obtained by a linear polarizer (LPVISE100-A, Thorlabs GmbH) and a quarter-wave plate (AQWP05M-600, Thorlabs GmbH, Dachau, Germany). The light was focused by an oil-immersion objective (UPLSAPO100XO, NA 1.40, Olympus Deutschland GmbH) onto the sample. The sample was moved by a piezo stage (P-517.3CD, Physik Instrumente (PI) GmbH & Co. KG, Karlsruhe, Germany) controlled by a E-727.3CDA piezo controller (Physik Instrumente (PI) GmbH & Co. KG, Karlsruhe, Germany). The emission was separated from the excitation beam by a dichroic beam splitter (z532/633, AHF analysentechnik AG, Tübingen, Germany) and focused onto a 50 µm pinhole (Thorlabs GmbH). The emission light was split by a dichroic beam splitter (640DCXR, AHF analysentechnik AG) into a green (Brightline HC582/75, AHF analysentechnik AG; RazorEdge LP 532, Laser 2000 GmbH, Weßling, Germany) and red (Shortpass 750, AHF Analysentechnik AG; RazorEdge LP 647, Laser 2000 GmbH) detection channel. Emission was focused onto avalanche photo diodes (SPCM-AQRH-14-TR, Excelitas Technoligies GmbH & Co. KG, Wiesbaden, Germany) and signals were registered by a time-correlated single photon counting (TCSPC)-unit (HydraHarp400, PicoQuant GmbH, Berlin, Germany). The setup was controlled by a commercial software package (SymPhoTime64, Picoquant GmbH, Berlin, Germany). Excitation powers of 36 µW and 25 µW were used for donor and acceptor lasers (as measured in front of the entrance of the microscope).

Cohesin-dockerin samples were incubated for a few seconds at a concentration of cohesin of 1 µM, at a cohesin:dockerin molar ratio of 1:1 for *C. thermocellum* samples and 1:2 for *C. cellulolyticum*. Samples were finally diluted to a concentration of ∼ 200-400 pM in 50 mM Tris, 100 mM NaCl, 2mM Trolox/Trolox quinone, 1% glucose, 1 mM $CaCl_2$, pH 8 for smFRET measurements.

In the case of the kinetics experiments performed with *C. cellulolyticum* complexes, samples were incubated at a concentration of cohesin of 1 µM and a cohesin:dockerin ratio of 1:2. We diluted the sample to 444 pM, and data acquisition was started 90 seconds after the proteins were mixed at micromolar concentration. In order to obtain significant statistics for every time point, the measurements were repeated 4-7 times for each sample and combined together. For cyclophilin A treatment, dockerin was first incubated in the presence of 20 µM of cyclophilin A for 10 minutes, after that, cohesin was added. Besides, the measurement was performed in the presence of 14 µM of cyclophilin A in the chamber. All measurement were measured at room temperature.

**smFRET analysis**

For the case of smFRET experiments in surface-immobilized molecules, FRET efficiencies (*E*) were calculated with the following simplified formula(*15*):

$$E = 1 - \frac{I_{DA}}{I_D}$$

(1)

Where $I_{DA}$ stands for intensity in the acceptor channel after donor excitation (FRET) before the acceptor dye bleaches, and $I_D$ for intensity in the donor channel after donor excitation after bleaching of acceptor dye. All intensities are background-corrected

Burst selection in smFRET solution experiments was performed using a sliding time window burst search, with a time window of 500 µs, a minimum of 4 photon *per* time window, and threshold for burst detection of 40 photons(*16*). These detection parameters are quite loose; they were chosen to avoid detection bias toward possibly more bright population of molecules. ALEX-2CDE (*17*) and lTDX-TAAl filters (*18*) were applied to sort out photobleaching and blinking events. Doubled-labeled molecules where further selected by keeping the stoichiometry parameter between 0.2 and 0.8. Accurate FRET efficiencies(*19, 20*) were calculated from fluorescence intensities as:

$$E = \frac{I_{DA} - \alpha I_{DD} - \delta I_{AA}}{\gamma I_{DD} + I_{DA} - \alpha I_{DD} - \delta I_{AA}}$$

(2)

where $I_{AA}$, $I_{DD}$ and $I_{DA}$ are the background-corrected photon counts in the acceptor channel after acceptor excitation, the donor channel after donor excitation and the acceptor channel after donor excitation. $\alpha$ and $\delta$ correction parameters are calculated from donor only and acceptor only subpopulations and account for spectral cross talk and direct excitation of the donor dye. The different detection efficiencies and quantum yields of fluorophores are corrected with the $\gamma$ correction factor obtained from global fittings of 1/S *vs* E plots(*19, 20*) . When proximity ratios are calculated, E(PR), these parameters are set to $\alpha$ = 0, $\delta$ = 0, and $\gamma$ = 1. In order to present the data as raw as possible, we choose to show FRET efficiencies as E(PR) histograms in the main text and supplementary figures, nevertheless all data set where fully corrected for PDA analysis, and a set of corrected data is shown in Fig. S5b as an example.

**PDA analysis**

Quantitative PDA analysis(*21-23*) was carried out using the free software PAM. As described in the supplementary results section, and in order to reduce the impact of the elevated number of free fitting parameters, the structural parameter *R* and $\sigma$ were fitted globally for the same type of samples. Besides, low FRET and high FRET subpopulations were included as samples in the fitting (see Supplementary results for a detailed discussion of why this protocol was followed). Förster Radius ($R_0$ = 69.12 Å) was calculated using the overlap integrals from donor emission spectra (Cy3B) and acceptor absorption spectra (Alexa Fluor 647). The time binning was set to 0.8 ms. Fitting parameter error represent confidence intervals at 95%. Fitting parameters for all analyzed samples can be found in Supplementary Table 1.

For the kinetics analysis of c1C-CcCel5A samples, the data set was divided into intervals of 150-300 seconds, the subpopulations ratios were then estimated by PDA and the obtained ratio was considered as representative of the middle point of the interval. Please note that we have no data of the first 90s of kinetics (see smFRET measurements).

We have shown (see Supplementary results) that PDA can estimate the population with a deviation of 0.007 from the expected value. When we report the population fraction of a single sample, the confidence interval at 95% was used as error, since they were always larger than this lower limit. When different populations where compared, the standard deviation of the population was used as error as long as they were larger than the lower limit of 0.007, otherwise the confidence interval at 95% was used.

**Kinetic analysis**

We analyzed the kinetics process of c1C-CcCel5A samples as a first order kinetics process with pseudo-direct interconversion between the B1 and the B2 state. Please note that, very likely, the process will not involve direct interconversion between the states, since it will require separation of the proteins for a few nanometers and also rotation of 180 $^{o}$ of the dockerin. It will be very probable that the proteins would diffuse apart during the whole process. More likely, the real process involves exchange of the dockerin from one binding mode to the pool of free dockerins and then the binding in the alternative mode. Our data back this scenario, in Fig. S10 can be seen that the depopulation of B1 state is not instantaneously associated with an increase in the B2 state (see for example the first 500 seconds in Fig. S10A) as it would be expected from a direct interconversion between states. This is even more obvious in cyclophilin treated samples, where a release of cohesins to the free-cohesin pool (yellow, Fig. S10B) is concomitant to the re-equilibration process between binding modes. Besides, burst variance analysis (*24*) and the FRET-2CDE filter (*17*) do not show dynamics during the burst duration (Fig S12), and we did not observe interconversion between binding modes in the range of seconds in our measurements with surface-immobilized molecules.

The kinetic data were fit with a monoexponential curve of the type:

$$B_1 = B_{eq} + Be^{-t/t_1} \tag{3}$$

where $B_1$ stands for the fraction of the population in the $B_1$ mode, $B_{eq}$ for the fraction at the final equilibrium, and $t_1$ is the decay time. In the context of our first order kinetics model with pseudo-direct interconversion of states, $t_1 = (1/(K_{12} + k_{21}))$, where $k_{12}$ and $k_{21}$ are the pseudo-kinetics constants of interconversion between B1 and B2.

**NMR measurements**

$^{15}$N-labeled Dockerin was dissolved to a final concentration of 0.68 mM in aqueous buffer containing 10 mM $KH_2PO_4$, 10 mM $CaCl_2$, 1 mM $NaN_3$ (sodium azide) to prevent microbial growth and 0.050 mM 4,4-dimethyl-4-silapentane-1-sulfonic acid (DSS), as the internal chemical shift $^1$H reference. The $^{15}$N chemical shift reference was calculated from the $^{1}H$ reference using the nuclei's gyromagnetic ratios. The pH of the solution was 6.23. NMR spectra were recorded at 25.0 °C using a Bruker 800 MHz spectrometer equipped with a triple resonance cryoprobe and Z-gradients. After recording 1D $^{1}H$ NMR, 2D $^{1}H$-$^{15}N$ HSQC and 3D $^{1}H$-$^{15}N$ HSQC-NOESY spectra on the $^{15}N$-labeled Dockerin sample alone, it was titrated with unlabeled Cohesin protein which had previously been dissolved in the same buffer. 1D $^{1}H$ and 2D $^{1}H$-$^{15}N$ HSQC spectra were recorded on the mixture at Dockerin:Cohesin ratios of 1.0 : 0.5, 1.0 : 0.8, and 1.0 : 1.3. At the final ratio, an additional 3D $^{1}H$-$^{15}N$ HSQC NOESY spectrum was also recorded. Spectra were transformed using Bruker Topspin 2.1 software and were analyzed using NMRFAM 1.4/Sparky 3.1. No linear prediction, which may affect relative peak intensities, was used in the data processing. The spectral acquisition parameters are listed in Table 2.

| Spectrum Type | # of Scans per Transient | Sweep Width (ppm) | Matrix Size | NOESY Mixing Time (ms) |
|---|---|---|---|---|
| 1D, $^{1}H$ | 32 | 14 | 32 | -- -- |
| 2D $^{1}H$-$^{15}N$ HSQC | 8 | 12 ($^{1}H$) x 28 ($^{15}N$) | 2k ($^{1}H$) x 256 ($^{15}N$) | -- -- |
| 3D $^{1}H$-$^{15}N$ HSQC-NOESY | 8 | 12 ($^{1}H$) x 28 ($^{15}N$) x 12 ($^{1}H$) | 2k ($^{1}H$, direct) x 48 ($^{15}N$) x 128 ($^{1}H$ indirect) | 100 |

**Table 2**: Acquisition Parameters for NMR Spectroscopy

### Molecular dynamics simulations

We use two methods to explore the energy landscape of the cohesin-dockerin complex between cohesin 2 and dockerin S (PDB code 2MTE) from *C. thermocellum.* The first method uses a FoldX-based approach(*25*) but in a significantly improved manner that involves a Monte Carlo sampling. The FoldX procedure optimizes the conformations of the sidechains while keeping the backbone geometry fixed. The second method involves a coarse-grained model introduced by Kim and Hummer (*26*) in which the molecules interact through the residues at the inter-molecular interface, but themselves are considered to be rigid bodies.

#### *The energy landscape of cohesin-dockerin complexes*

First, we compared our new Monte Carlo (MC) sampling FoldX procedure with our previous method(*25*). In order to do so, we analyzed the same cohesin-dockerin type I complex studied in that work and we benchmark our new results with those obtained previously. We investigated the complex formed between *C. thermocellum* CipA cohesin 2 and the dockerin from Xylanase 10B (c2A-Xyn10B complex).

The structure of this complex has been solved in both binding modes, and their structures can be found under the PDB code 1OHZ(*6*) for B1 mode, and 2CLL(*5*) for B2 state. Unfortunately, neither of the two structures carries the C-terminal stretch of the last 6 residues and these segments have to be reconstructed from selected templates (we used PDB: 4DH2, 1OHZ and 2CCL) by employing a set of structure-prediction servers. We used for that purpose Swiss Model(*27*) and iTASER(*28*).

The determination of the free-energy landscape is facilitated by first establishing the direction of a symmetry axis, *Z*, such that a rotation of dockerin around *Z* by an angle $\varphi$ transforms the cohesin-dockerin structure between the two binding modes. The landscape is described in terms of the variables *Z* and $\varphi$. The coordinates of cohesin-dockerin at *Z*=0 and $\varphi$=0 correspond to the conformation of PDB:1OHZ. Similarly, $Z \approx 0$ and $\varphi \approx \pi$ corresponds to the conformation of PDB:2CCL. We use the convention in which positive values of *Z* correspond to shifting the two molecules closer together and negative values to shifting them further away. The determination of the free-energy landscape is facilitated by first establishing the direction of the symmetry axis, *Z*, such that a rotation of the dockerin around *Z* by an angle transforms the cohesin-dockerin structure between the two binding modes.

The program FoldX (*29, 30*) has been designed primarily for predicting free energy differences between a WT (wild type) protein and its mutant. FoldX employs an energy function that consists of ten terms and takes the entropy into account in an empirical way. The free energy is optimized with respect to the side-chain conformations while the backbone atoms are kept at fixed positions. Our calculations are performed at temperature T =298 K and the calcium ions are taken into account (*31*).

The resulting free energy *ΔG* depends not only on the coordinates *Z* and $\varphi$ but also on the structural model considered. The model involves substantial uncertainty, and therefore we used the 12 predicted WT structures for our Monte Carlo runs (see bellow for a description of these 12 predicted structures). Furthermore, the binding energy, $F_I$, involved in attaching in B1 is obtained by summing over all values of *Z* and $\varphi$ with the condition that $\varphi$ is between −π/2 and about π/2.

$$F_I = -k_B T \ \log\left[\sum_i \sum_Z \sum_{-\frac{\pi}{2}<\varphi<\frac{\pi}{2}} \exp\left(\frac{-\Delta G_i(Z,\varphi)}{k_B T}\right)\theta(E_c - \Delta G_i)\right] \quad (4)$$

The sum over index $i$ corresponds to averaging over the structures considered from the Monte Carlo sampling. $\Delta G_i (Z, \varphi)$ denotes $\Delta G(Z, \varphi)$ computed for the input structure with index $i$. The unit step function, $\theta$, is defined as follows: $\theta$(x) = 1 for x > 0 and $\theta$(x) = 0 for x < 0. The structures corresponding to the B1 mode are those with $-\pi/2 < \varphi < \pi/2$ and free energies $\Delta G_i$ smaller than a cut-off value $E_c$. Our results show that $F_I$ does not depend on the choice of the cut-off as long as $E_c$ is large enough (see Fig. 5D and Sup. Results). Correspondingly, the binding energy in B2, $F_{II}$ incorporates $\varphi$ between $\pi/2$ and $3\pi/2$. The ratio of probabilities of binding in modes I and II is then equal to $p = pI/pII = exp[-(FI - FII)/k_BT]$.

We performed a Monte Carlo run in which the energies *ΔG* were calculated for various values of *Z* and $\varphi$ using FoldX. The maximal allowed value of rotation in a single step was 2° and of translation 0.2 Å. We then introduced a grid in which the nodes along the (global or local) $Z$-axis are $\delta Z$=0.1 Å apart and the angle $\varphi$ changes in steps of $\delta\varphi$ =1°. The lowest $\Delta G$ in the bin was associated with the grid site. A sufficiently long run should allow for switches between the basins corresponding to the B1 and B2 state situations.

We performed many MC simulation runs. For each initial cohesin-dockerin structure, we performed 10 simulation runs. It was done to effectively probe possible configurations of the cohesin-dockerin complex. Moreover, we used many different starting configurations. In the case of 1OHZ/2CCL (c2A-Xyn10B), we had six independent modeled structures of dockerin (since the terminal tails were absent in the original PDB structures). We modeled the tails using two PDB entries: 1OHZ and 2CCL, for each one we generated three structures, one coming from Swiss Model, and two coming from iTASER. For details please check Table 2 in(*25*) index from 9-14, where there is a typo and *d* should be read as *D*. For each of these six initial structures, three in the B1 (1OHZ-derived structures) and three in the B2 state (2CCL-derived structures), we generated another six initial states in the alternative binding mode by rotating the dockerin around 180°. This summed up for a total of twelve initial structures, six in each binding mode. The diversity of the initial configurations allowed us to better sample the configurational space of the cohesin-dockerin complex.

To further improve sampling of the configurational space, we generated an even larger set of initial configurations by rotating the dockerin around the $Z$-axis by nine starting $\varphi$ angles (−40°, −30°, −20°, −10°, 0°, 10°, 20°, 30°, 40°). In this way we produced 12*10*9 simulation trajectories, each of them consisting of N = 12000 MC steps. We thus obtained 12*10*9*12000 values of the cohesin-dockerin interaction energy. Dissociative events come with a high energy so they do not affect the simulations. Importantly, in MC procedure, we are not interested in one continuous trajectory. We merge all the MC simulation data into a single pool of conformations. Next, for given values of angle $\varphi$ and separation $Z$ of the grid, we select one conformation that has the lowest energy. In this way, each point in the ($Z$, $\varphi$) space is assigned the lowest energy of the cohesin-dockerin interactions, which is then used to compute the $F_I$ and $F_{II}$ values according to Eq. (4).

Please note that in our previous approach only six cohesin-dockerin structures were sampled in order to calculate the binding energy $F_I$ and $F_{II}$, while in the new Monte Carlo approach the $\Delta G$ of millions of cohesin/dockerin structures are explored, providing a thorough sampling of the energy landscape. The results are discussed in the Sup. Results section.

In the case of the c2A-celS complex, the whole-sequence dockerin structure is available (2MTE). Since the conformations were determined by NMR, there are 20 versions of this structure. In our calculations we take them all into account (in an analogy to considering the 6 predicted structures derived for 1OHZ/2CCL). In this case, we don't have structure of the cohesin-dockerin complex and therefore we take the coordinates of cohesin from either 1OHZ or 2CCL. We did not consider initial configurations generated by the rotations around the Z-axis. The case of 1OHZ/2CLL actually showed that in almost all simulations started from $\varphi$ not equal to 0, the system evolved quickly towards configurations with $\varphi$ close to 0. For the 20 starting structures we start the MC runs from either the B1 mode ($Z$ = 0, $\varphi = 0°$), or the B2 mode ($Z \approx 0$, $\varphi \approx 180°$).

We have considered two possible calculational schemes. In the first one, we take the *Z*-axis to as determined before for the 1OHZ/2CCL system (*25*) also for the 2MTE case (c2A-CelS complex) even though the symmetries for the new system may be distinct. This scheme will be referred to as uniaxial. Another scheme, referred to as multiaxial, involves many axes of rotations that are chosen randomly.

We generated the structures of c2A-celS_SB1 and c2A-celS_mClasp mutants using FoldX by starting from PDB:2MTE. All original sidechains of the mutated residues were deleted and the software generated the structures of the sidechains of the residues introduced by mutation. In the case of the C-terminal depletion, we simply cut off the C-terminal amino acids and did not do any backbone remodeling. The trajectories are 8,000 steps long of the WT complex, 4,000 for the c2A-celS_B1 complex, 8,000 in the case of the clasp mutant, and 20,000 steps for the C-terminal depleted dockerin.

When calculating the RMSD for a given simulation, we take the starting structure as a reference for the version of the complex that is being simulated.

*The coarse-grained model*

To efficiently sample conformations of the c2A-CelS complex, we used an implicit-solvent, coarse-grained model introduced by Kim and Hummer (*26*). This model is equipped with a transferable energy function and devised for simulating conformational ensembles of multi-protein complexes. It has been successfully applied to systems ranging from the ESCRT membrane-protein trafficking machinery (*32, 33*), to lipid kinases (*34, 35*) and complexes of cell adhesion proteins involved in immunological responses (*36*).

In the framework of this model, amino-acid residues are represented as spherical beads centered at the α-C atoms. The interactions between the beads are described by statistical amino-acid dependent Lennard-Jones-type potentials and Debye-Huckel-type electrostatics. Folded protein domains are simulated as rigid bodies formed of the amino-acid beads. Here, the cohesin and dockerin modules are treated as two separate rigid bodies. Detailed description of this model together with its parameterization is provided in(*26*).

Within the framework of this model, we performed replica exchange Monte Carlo simulations of the c2A-CelS system using in-house software with replicas at 12 temperatures $T_i$ given by $T/T_i$ = 0.6,0.64,...,1.0,1.04 relative to the room temperature $T$ =300 K (which is nearly the same as in the FoldX-based calculations). The basic MC steps were rigid body translational and rotational moves on the cohesin and dockerin domains. The probabilities of the MC moves were governed by the Metropolis criterion. The simulations were done in a cubic box with dimensions $L_x = L_y = L_z$ = 20 nm and with periodic boundary conditions. The simulations were started from an unbound configuration in which the centers of the cohesin and dockerin modules were separated by about 5 nm. After the initial $10^6$ MC moves for equilibration, $10^8$ MC moves were performed for the data acquisition. The protein conformations were saved every $10^4$ MC moves at $T_i = T$, which gave us an ensemble of $10^4$ conformations for further analysis. We assumed that a cohesin-dockerin complex was formed if the protein-pair interaction energy was smaller than -2 $k_BT$, as in(*26*).

Our task is to predict how a general cohesin-dockerin complex should be arranged. Let $S_I$ denote the proposed cohesin-dockerin structure that is bound in B1 mode and $S_{II}$ the structure that is bound in B2 mode. For each structure S obtained during simulations in a coarse-grained model based on the dynamics of the α-C atoms, we can define the distance root mean square (DRMS) to the proposed structures $S_I$ and $S_{II}$. They are defined as follows:

$$DMRS\ (S, S_I) = \left[ \frac{1}{N_2} \sum_{ij} \left( d_{ij}^{S} - d_{ij}^{S_I} \right)^2 \right]^{1/2} \tag{5}$$

And

$$DMRS\ (S, S_{II}) = \left[ \frac{1}{N_2} \sum_{ij} \left( d_{ij}^{S} - d_{ij}^{S_{II}} \right)^2 \right]^{1/2} \tag{6}$$

Here, $d_{ij}^{(S_I)}$ is the Cartesian distance between the $\alpha$-C atoms of residues $i$ and $j$ in two different modules of structure $S_I$; $d_{ij}^{(S_{II})}$ is the Cartesian distance between the $\alpha$-C atoms of residues $i$ and $j$ in two different modules of structure $S_{II}$; and $N_2$ is the number of residue pairs over which the sum is performed. If a typical value of $DRMS(S_I,S_{II})$ is $d_{I,II}$ then the structures with $DRMS(S,S_I) < d_c$, where $d_c = d_{I,II}/2$, are plausibly assigned to B1 mode. Similarly, the structures with $DRMS(S,S_{II}) < d_c$ are assigned to B2 mode.

When applied to the complex c2A-CelS we get $d_{I,II}$ ≈8 Å so $d_c$ is 4 Å. We also compute the Cartesian distance between the $\alpha$-C atoms of $Cys^{142}$ in cohesin and $Cys^{69}$ in dockerin. These are the Cys residues to which the fluorescent labels are attached in the single-molecule FRET experiments.

For each of the simulated 2MTE containing systems, we determined the number $n_I(d)$ of structures $S$ found in the binding mode 1 with DRMS$(S,S_I) < d$. By analogy, we also computed the number $n_{II}(d)$ of structures S belonging to the binding mode 2 with DRMS$(S,S_{II}) < d$. Fig. S7B shows the ratios $p_I = n_I/(n_I + n_{II})$ (blue lines) and $p_{II} = n_{II}/(n_I + n_{II}) = 1 - p_I$ (red lines) as functions of $d$.

**List of proteins:**

Code:

Blue xylanase T6

Red residue mutated to cysteine for dye labeling

Green Aa residue from restriction enzyme (Aa residue which come from cloning)

Yellow : histag

**celS:**

MASKNADSYAKKPHISALNAPQLDQRYKNEFTIGAAVEPYQLQNEKDVQMLKRHFNSIVAENVMKPISIQPEEGKFNF
EQADRIVKFAKANGMDIRFHTLVWHSQVPQWFFLDKEGKPMVNETDPVKREQNKQLLLKRLETHIKTIVERYKDDIKY
WDVVNEVVGDDGKLRNSPWYQIAGIDYIKVAFQAARKYGGDNIKLYMNDYNTEVEPKRTALYNLVKQLKEEGVPIDGI
GHQSHIQIGWPSEAEIEKTINMFAALGLDNQITELDVSMYGWPPRAYPTYDAIPKQKFLDQAARYDRLFKLYEKLSDKI
SNVTFWGIADNHTWLDSRADVYYDANGNVVVDPNAPYAKVEKGKGKDAPFVFGPDYKVKPAYWAIIDHKGSVVPGT
PSTKLYGDVNDDGKVNSTDAVALKRYVLRSGISINTDNADLNEDGRVNSTDLGILKRYILKEICTLPYKNLEHHHHHH

**celS_SB1:**

MASKNADSYAKKPHISALNAPQLDQRYKNEFTIGAAVEPYQLQNEKDVQMLKRHFNSIVAENVMKPISIQPEEGKFNF
EQADRIVKFAKANGMDIRFHTLVWHSQVPQWFFLDKEGKPMVNETDPVKREQNKQLLLKRLETHIKTIVERYKDDIKY
WDVVNEVVGDDGKLRNSPWYQIAGIDYIKVAFQAARKYGGDNIKLYMNDYNTEVEPKRTALYNLVKQLKEEGVPIDGI
GHQSHIQIGWPSEAEIEKTINMFAALGLDNQITELDVSMYGWPPRAYPTYDAIPKQKFLDQAARYDRLFKLYEKLSDKI
SNVTFWGIADNHTWLDSRADVYYDANGNVVVDPNAPYAKVEKGKGKDAPFVFGPDYKVKPAYWAIIDHKGSVVPGT
PSTKLYGDVNDDGKVNDEDAVALAAYVLRSGISINTDNADLNEDGRVNSTDLGILKRYILKEICTLPYKNLEHHHHHH

**celS_SB1.1:**

MASKNADSYAKKPHISALNAPQLDQRYKNEFTIGAAVEPYQLQNEKDVQMLKRHFNSIVAENVMKPISIQPEEGKFNF
EQADRIVKFAKANGMDIRFHTLVWHSQVPQWFFLDKEGKPMVNETDPVKREQNKQLLLKRLETHIKTIVERYKDDIKY
WDVVNEVVGDDGKLRNSPWYQIAGIDYIKVAFQAARKYGGDNIKLYMNDYNTEVEPKRTALYNLVKQLKEEGVPIDGI
GHQSHIQIGWPSEAEIEKTINMFAALGLDNQITELDVSMYGWPPRAYPTYDAIPKQKFLDQAARYDRLFKLYEKLSDKI
SNVTFWGIADNHTWLDSRADVYYDANGNVVVDPNAPYAKVEKGKGKDAPFVFGPDYKVKPAYWAIIDHKGSVVPGT
PSTKLYGDVNDDGKVNSTDAVALKRYVLRSGISINTDNADLNEDGRVNAADLGILKRYILKEICTLPYKNLEHHHHHH

**celS_SB2:**

MASKNADSYAKKPHISALNAPQLDQRYKNEFTIGAAVEPYQLQNEKDVQMLKRHFNSIVAENVMKPISIQPEEGKFNF
EQADRIVKFAKANGMDIRFHTLVWHSQVPQWFFLDKEGKPMVNETDPVKREQNKQLLLKRLETHIKTIVERYKDDIKY
WDVVNEVVGDDGKLRNSPWYQIAGIDYIKVAFQAARKYGGDNIKLYMNDYNTEVEPKRTALYNLVKQLKEEGVPIDGI
GHQSHIQIGWPSEAEIEKTINMFAALGLDNQITELDVSMYGWPPRAYPTYDAIPKQKFLDQAARYDRLFKLYEKLSDKI
SNVTFWGIADNHTWLDSRADVYYDANGNVVVDPNAPYAKVEKGKGKDAPFVFGPDYKVKPAYWAIIDHKGSVVPGT
PSTKLYGDVNDDGKVNSTDAVALKRYVLRSGISINTDNADLNEDGRVNDEDLGILAAYILKEICTLPYKNLEHHHHHH

**CelS_mClasp:**

MASKNADSYAKKPHISALNAPQLDQRYKNEFTIGAAVEPYQLQNEKDVQMLKRHFNSIVAENVMKPISIQPEEGKFNF
EQADRIVKFAKANGMDIRFHTLVWHSQVPQWFFLDKEGKPMVNETDPVKREQNKQLLLKRLETHIKTIVERYKDDIKY
WDVVNEVVGDDGKLRNSPWYQIAGIDYIKVAFQAARKYGGDNIKLYMNDYNTEVEPKRTALYNLVKQLKEEGVPIDGI
GHQSHIQIGWPSEAEIEKTINMFAALGLDNQITELDVSMYGWPPRAYPTYDAIPKQKFLDQAARYDRLFKLYEKLSDKI
SNVTFWGIADNHTWLDSRADVYYDANGNVVVDPNAPYAKVEKGKGKDAPFVFGPDYKVKPAYWAIIDHKGSVVPGT
PSTKLAGDVNDDGKVNSTDAVALKRYVLRSGISINTDNADLNEDGRVNSTDLGILKRYILKEICTLAYKNLEHHHHHH

**CelS_c-Depleted:**

MASKNADSYAKKPHISALNAPQLDQRYKNEFTIGAAVEPYQLQNEKDVQMLKRHFNSIVAENVMKPISIQPEEGKFNF
EQADRIVKFAKANGMDIRFHTLVWHSQVPQWFFLDKEGKPMVNETDPVKREQNKQLLLKRLETHIKTIVERYKDDIKY
WDVVNEVVGDDGKLRNSPWYQIAGIDYIKVAFQAARKYGGDNIKLYMNDYNTEVEPKRTALYNLVKQLKEEGVPIDGI
GHQSHIQIGWPSEAEIEKTINMFAALGLDNQITELDVSMYGWPPRAYPTYDAIPKQKFLDQAARYDRLFKLYEKLSDKI
SNVTFWGIADNHTWLDSRADVYYDANGNVVVDPNAPYAKVEKGKGKDAPFVFGPDYKVKPAYWAIIDHKGSVVPGT
PSTKLYGDVNDDGKVNSTDAVALKRYVLRSGISINTDNADLNEDGRVNSTDLGILKRYILKEICGGLEHHHHHH

**CelS_mSwap:**

MASKNADSYAKKPHISALNAPQLDQRYKNEFTIGAAVEPYQLQNEKDVQMLKRHFNSIVAENVMKPISIQPEEGKFNF
EQADRIVKFAKANGMDIRFHTLVWHSQVPQWFFLDKEGKPMVNETDPVKREQNKQLLLKRLETHIKTIVERYKDDIKY
WDVVNEVVGDDGKLRNSPWYQIAGIDYIKVAFQAARKYGGDNIKLYMNDYNTEVEPKRTALYNLVKQLKEEGVPIDGI
GHQSHIQIGWPSEAEIEKTINMFAALGLDNQITELDVSMYGWPPRAYPTYDAIPKQKFLDQAARYDRLFKLYEKLSDKI
SNVTFWGIADNHTWLDSRADVYYDANGNVVVDPNAPYAKVEKGKGKDAPFVFGPDYKVKPAYWAIIDHKGSVVPGT
PSTKLYGDVNDDGKVNSTDLGILKRYILKSGISINTDNADLNEDGRVNSTDAVALKRYVLREICTLPYKNLEHHHHHH

**celA:**

MASKNADSYAKKPHISALNAPQLDQRYKNEFTIGAAVEPYQLQNEKDVQMLKRHFNSIVAENVMKPISIQPEEGKFNF
EQADRIVKFAKANGMDIRFHTLVWHSQVPQWFFLDKEGKPMVNETDPVKREQNKQLLLKRLETHIKTIVERYKDDIKY
WDVVNEVVGDDGKLRNSPWYQIAGIDYIKVAFQAARKYGGDNIKLYMNDYNTEVEPKRTALYNLVKQLKEEGVPIDGI
GHQSHIQIGWPSEAEIEKTINMFAALGLDNQITELDVSMYGWPPRAYPTYDAIPKQKFLDQAARYDRLFKLYEKLSDKI
SNVTFWGIADNHTWLDSRADVYYDANGNVVVDPNAPYAKVEKGKGKDAPFVFGPDYKVKPAYWAIIDHKGSPTPSL
PPQVVYGDVNGDGNVNSTDLTMLKRYLLKSVTNINREAADVNRDGAINSSDMTILKRYLIKSICHLPYLEHHHHHH

**celA_SB1:**

MASKNADSYAKKPHISALNAPQLDQRYKNEFTIGAAVEPYQLQNEKDVQMLKRHFNSIVAENVMKPISIQPEEGKFNF
EQADRIVKFAKANGMDIRFHTLVWHSQVPQWFFLDKEGKPMVNETDPVKREQNKQLLLKRLETHIKTIVERYKDDIKY
WDVVNEVVGDDGKLRNSPWYQIAGIDYIKVAFQAARKYGGDNIKLYMNDYNTEVEPKRTALYNLVKQLKEEGVPIDGI
GHQSHIQIGWPSEAEIEKTINMFAALGLDNQITELDVSMYGWPPRAYPTYDAIPKQKFLDQAARYDRLFKLYEKLSDKI
SNVTFWGIADNHTWLDSRADVYYDANGNVVVDPNAPYAKVEKGKGKDAPFVFGPDYKVKPAYWAIIDHKGSPTPSL
PPQVVYGDVNGDGNVNDEDLTMLAAYLLKSVTNINREAADVNRDGAINSSDMTILKRYLIKSICHLPYLEHHHHHH

**celA_SB2:**

MASKNADSYAKKPHISALNAPQLDQRYKNEFTIGAAVEPYQLQNEKDVQMLKRHFNSIVAENVMKPISIQPEEGKFNF
EQADRIVKFAKANGMDIRFHTLVWHSQVPQWFFLDKEGKPMVNETDPVKREQNKQLLLKRLETHIKTIVERYKDDIKY
WDVVNEVVGDDGKLRNSPWYQIAGIDYIKVAFQAARKYGGDNIKLYMNDYNTEVEPKRTALYNLVKQLKEEGVPIDGI
GHQSHIQIGWPSEAEIEKTINMFAALGLDNQITELDVSMYGWPPRAYPTYDAIPKQKFLDQAARYDRLFKLYEKLSDKI
SNVTFWGIADNHTWLDSRADVYYDANGNVVVDPNAPYAKVEKGKGKDAPFVFGPDYKVKPAYWAIIDHKGSPTPSL
PPQVVYGDVNGDGNVNSTDLTMLKRYLLKSVTNINREAADVNRDGAINDEDMTILAAYLIKSICHLPYLEHHHHHH

**celA_mClasp:**

MASKNADSYAKKPHISALNAPQLDQRYKNEFTIGAAVEPYQLQNEKDVQMLKRHFNSIVAENVMKPISIQPEEGKFNF
EQADRIVKFAKANGMDIRFHTLVWHSQVPQWFFLDKEGKPMVNETDPVKREQNKQLLLKRLETHIKTIVERYKDDIKY
WDVVNEVVGDDGKLRNSPWYQIAGIDYIKVAFQAARKYGGDNIKLYMNDYNTEVEPKRTALYNLVKQLKEEGVPIDGI
GHQSHIQIGWPSEAEIEKTINMFAALGLDNQITELDVSMYGWPPRAYPTYDAIPKQKFLDQAARYDRLFKLYEKLSDKI
SNVTFWGIADNHTWLDSRADVYYDANGNVVVDPNAPYAKVEKGKGKDAPFVFGPDYKVKPAYWAIIDHKGSPTPSL
PPQVVAGDVNGDGNVNSTDLTMLKRYLLKSVTNINREAADVNRDGAINSSDMTILKRYLIKSICHLAYLEHHHHHH

**Ccel5A:**

MASKNADSYAKKPHISALNAPQLDQRYKNEFTIGAAVEPYQLQNEKDVQMLKRHFNSIVAENVMKPISIQPEEGKFNF
EQADRIVKFAKANGMDIRFHTLVWHSQVPQWFFLDKEGKPMVNETDPVKREQNKQLLLKRLETHIKTIVERYKDDIKY
WDVVNEVVGDDGKLRNSPWYQIAGIDYIKVAFQAARKYGGDNIKLYMNDYNTEVEPKRTALYNLVKQLKEEGVPIDGI
GHQSHIQIGWPSEAEIEKTINMFAALGLDNQITELDVSMYGWPPRAYPTYDAIPKQKFLDQAARYDRLFKLYEKLSDKI
SNVTFWGIADNHTWLDSRADVYYDANGNVVVDPNAPYAKVEKGKGKDAPFVFGPDYKVKPAYWAIIDHKGSPVIVYG
DYNNDGNVDALDFAGLKKYIMAADHAYVKNLDVNLDNEVNAFDLAILKKYLLGMVCKLPSNLEHHHHHH

**Ccel5A_SB1:**

MASKNADSYAKKPHISALNAPQLDQRYKNEFTIGAAVEPYQLQNEKDVQMLKRHFNSIVAENVMKPISIQPEEGKFNF
EQADRIVKFAKANGMDIRFHTLVWHSQVPQWFFLDKEGKPMVNETDPVKREQNKQLLLKRLETHIKTIVERYKDDIKY
WDVVNEVVGDDGKLRNSPWYQIAGIDYIKVAFQAARKYGGDNIKLYMNDYNTEVEPKRTALYNLVKQLKEEGVPIDGI
GHQSHIQIGWPSEAEIEKTINMFAALGLDNQITELDVSMYGWPPRAYPTYDAIPKQKFLDQAARYDRLFKLYEKLSDKI
SNVTFWGIADNHTWLDSRADVYYDANGNVVVDPNAPYAKVEKGKGKDAPFVFGPDYKVKPAYWAIIDHKGSPVIVYG
DYNNDGNVDSTDFAGLKKYIMAADHAYVKNLDVNLDNEVNAFDLAILKKYLLGMVCKLPSNLEHHHHHH

**Ccel5A_SB2:**

MASKNADSYAKKPHISALNAPQLDQRYKNEFTIGAAVEPYQLQNEKDVQMLKRHFNSIVAENVMKPISIQPEEGKFNF
EQADRIVKFAKANGMDIRFHTLVWHSQVPQWFFLDKEGKPMVNETDPVKREQNKQLLLKRLETHIKTIVERYKDDIKY

WDVVNEVVGDDGKLRNSPWYQIAGIDYIKVAFQAARKYGGDNIKLYMNDYNTEVEPKRTALYNLVKQLKEEGVPIDGI
GHQSHIQIGWPSEAEIEKTINMFAALGLDNQITELDVSMYGWPPRAYPTYDAIPKQKFLDQAARYDRLFKLYEKLSDKI
SNVTFWGIADNHTWLDSRADVYYDANGNVVVDPNAPYAKVEKGKGKDAPFVFGPDYKVKPAYWAIIDHKGSPVIVYG
DYNNDGNVDALDFAGLKKYIMAADHAYVKNLDVNLDNEVNSTDLAILKKYLLGMVCKLPSNLEHHHHHH

**Ccel5A_mClasp:**

MASKNADSYAKKPHISALNAPQLDQRYKNEFTIGAAVEPYQLQNEKDVQMLKRHFNSIVAENVMKPISIQPEEGKFNF
EQADRIVKFAKANGMDIRFHTLVWHSQVPQWFFLDKEGKPMVNETDPVKREQNKQLLLKRLETHIKTIVERYKDDIKY
WDVVNEVVGDDGKLRNSPWYQIAGIDYIKVAFQAARKYGGDNIKLYMNDYNTEVEPKRTALYNLVKQLKEEGVPIDGI
GHQSHIQIGWPSEAEIEKTINMFAALGLDNQITELDVSMYGWPPRAYPTYDAIPKQKFLDQAARYDRLFKLYEKLSDKI
SNVTFWGIADNHTWLDSRADVYYDANGNVVVDPNAPYAKVEKGKGKDAPFVFGPDYKVKPAYWAIIDHKGSPVIVYG
DANNDGNVDALDFAGLKKYIMAADHAYVKNLDVNLDNEVNAFDLAILKKYLLGMVCKLASNLEHHHHHH

**c2A:**

MDGVVVEIGKVTGSVGTTVEIPVYFRGVPSKGIANCDFVFRYDPNVLEIIGIDPGDIIVDPNPTKSFDTAIYPDRKIIVFLF
AEDSGTGAYAITKDGVFAKIRATVKSSAPGYITFDEVGGFADNDLVEQKVSFIDGGVNVCNALEHHHHHH

**c7A:**

MGAVRIKVDTVNAKPGDTVRIPVRFSGIPSKGIANCDFVYSYDPNVLEIIEIEPGELIVDPNPTKSFDTAVYPDRKMIVFL
FAEDSGTGAYAITEDGVFATIVAKVKSGAPNGLSVIKFVEVGGFANNDLVEQKTQFFDGGVNVCLEHHHHHH

**c1A:**

MGATMTVEIGKVTAAVGSKVEIPITLKGVPSKGMANCDFVLGYDPNVLEVTEVKPGSIIKDPDPSKSFDSAIYPDRKMIV
FLFAEDSGRGTYAITQDGVFATIVATVKSAAAAPITLLEVGAFADNDLVEISTTFVAGGVNLCLEHHHHHH

**c1C:**

MDSLKVTVGTANGKPGDTVTVPVTFADVAKMKNVGTCNFYLGYDASLLEVVSVDAGPIVKNAAVNFSSSASNGTISFL
FLDNTITDELITADGVFANIKFKLKSVTAKTTTPVTFKDGGAFGDGTMSKIASVTKTNGSVTIDPCTQLEHHHHHH

**c7C:**

MGKELKVAVGTASGKAGDTVTVPVTFADVATVGNVGTCNFYVTYDTNLLEVASVTPGSIVTNAAVNFSSSTSNGTISF
LFLDNTITDQLIKTDGTFAEIKFKLKSVTAKTTTPVAFKDGGAFGDGTMAKIATVTKTNGSVTIDVCLEHHHHHH

**Cyclophilin:**

MAAKGDPHVLLTTSAGNIELELDKQKAPVSVQNFVDYVNSGFYNNTTFHRVIPGFMIQGGGFTEQMQQKKPNPPIKN
EADNGLRNTRGTIAMARTADKDSATSQFFINVADNAFLDHGQRDFGYAVFGKVVKGMDVADKISQVPTHDVGPYQN
VPSKPVVILSAKVLPLEHHHHHH

**Cyclophilin R50A:**

MAAKGDPHVLLTTSAGNIELELDKQKAPVSVQNFVDYVNSGFYNNTTFHAVIPGFMIQGGGFTEQMQQKKPNPPIKN
EADNGLRNTRGTIAMARTADKDSATSQFFINVADNAFLDHGQRDFGYAVFGKVVKGMDVADKISQVPTHDVGPYQN
VPSKPVVILSAKVLPLEHHHHHH

Methods references

1. J. Sambrook, *Molecular cloning : a laboratory manual*. (Third edition. Cold Spring Harbor, N.Y. : Cold Spring Harbor Laboratory Press, [2001] ©2001, 2001).
2. S. W. Stahl *et al.*, Single-molecule dissection of the high-affinity cohesin–dockerin complex. *Proceedings of the National Academy of Sciences* **109**, 20431-20436 (2012).
3. A. Valbuena *et al.*, On the remarkable mechanostability of scaffoldins and the mechanical clamp motif. *Proc Natl Acad Sci U S A* **106**, 13791-13796 (2009).
4. C. Chen *et al.*, Revisiting the NMR solution structure of the Cel48S type-I dockerin module from Clostridium thermocellum reveals a cohesin-primed conformation. *J Struct Biol* **188**, 188-193 (2014).
5. A. L. Carvalho *et al.*, Evidence for a dual binding mode of dockerin modules to cohesins. *Proceedings of the National Academy of Sciences* **104**, 3089 (2007).
6. A. L. Carvalho *et al.*, Cellulosome assembly revealed by the crystal structure of the cohesin–dockerin complex. *Proceedings of the National Academy of Sciences* **100**, 13809 (2003).
7. A. Galera-Prat, D. Pantoja-Uceda, D. V. Laurents, M. Carrion-Vazquez, Solution conformation of a cohesin module and its scaffoldin linker from a prototypical cellulosome. *Archives of biochemistry and biophysics* **644**, 1-7 (2018).
8. J. Marley, M. Lu, C. Bracken, A method for efficient isotopic labeling of recombinant proteins. *J Biomol NMR* **20**, 71-75 (2001).
9. M. G. Cull, P. J. Schatz, in *Methods in Enzymology*. (Academic Press, 2000), vol. 326, pp. 430-440.
10. M. Fairhead, M. Howarth, in *Site-Specific Protein Labeling: Methods and Protocols,* A. Gautier, M. J. Hinner, Eds. (Springer New York, New York, NY, 2015), pp. 171-184.
11. M. Fairhead, M. Howarth, Site-specific biotinylation of purified proteins using BirA. *Methods in molecular biology (Clifton, N.J.)* **1266**, 171–184 (2015).
12. A. N. Kapanidis *et al.*, Fluorescence-aided molecule sorting: Analysis of structure and interactions by alternating-laser excitation of single molecules. **101**, 8936-8941 (2004).
13. T. Ha, P. Tinnefeld, Photophysics of Fluorescent Probes for Single-Molecule Biophysics and Super-Resolution Imaging. **63**, 595-617 (2012).
14. B. K. Müller, E. Zaychikov, C. Bräuchle, D. C. Lamb, Pulsed interleaved excitation. *Biophysical journal* **89**, 3508-3522 (2005).
15. J. Bohlen *et al.*, Plasmon-assisted Förster resonance energy transfer at the single-molecule level in the moderate quenching regime. *Nanoscale* **11**, 7674-7681 (2019).
16. E. Nir *et al.*, Shot-noise limited single-molecule FRET histograms: comparison between theory and experiments. *J Phys Chem B* **110**, 22103-22124 (2006).
17. T. E. Tomov *et al.*, Disentangling subpopulations in single-molecule FRET and ALEX experiments with photon distribution analysis. *Biophys J* **102**, 1163-1173 (2012).
18. V. Kudryavtsev *et al.*, Combining MFD and PIE for accurate single-pair Forster resonance energy transfer measurements. *Chemphyschem : a European journal of chemical physics and physical chemistry* **13**, 1060-1078 (2012).
19. B. Hellenkamp *et al.*, Precision and accuracy of single-molecule FRET measurements—a multi-laboratory benchmark study. *Nature Methods* **15**, 669-676 (2018).
20. N. K. Lee *et al.*, Accurate FRET Measurements within Single Diffusing Biomolecules Using Alternating-Laser Excitation. *Biophysical Journal* **88**, 2939-2953 (2005).
21. M. Antonik, S. Felekyan, A. Gaiduk, C. A. Seidel, Separating structural heterogeneities from stochastic variations in fluorescence resonance energy transfer distributions via photon distribution analysis. *J Phys Chem B* **110**, 6970-6978 (2006).
22. S. Kalinin, S. Felekyan, A. Valeri, C. A. M. Seidel, Characterizing Multiple Molecular States in Single-Molecule Multiparameter Fluorescence Detection by Probability Distribution Analysis. *The Journal of Physical Chemistry B* **112**, 8361-8374 (2008).

23. E. Sisamakis, A. Valeri, S. Kalinin, P. J. Rothwell, C. A. Seidel, Accurate single-molecule FRET studies using multiparameter fluorescence detection. *Methods Enzymol* **475**, 455-514 (2010).
24. J. P. Torella, S. J. Holden, Y. Santoso, J. Hohlbein, A. N. Kapanidis, Identifying molecular dynamics in single-molecule FRET experiments with burst variance analysis. *Biophysical journal* **100**, 1568-1577 (2011).
25. M. Wojciechowski *et al.*, Dual binding in cohesin-dockerin complexes: the energy landscape and the role of short, terminal segments of the dockerin module. *Scientific Reports* **8**, 5051 (2018).
26. Y. C. Kim, G. Hummer, Coarse-grained models for simulations of multiprotein complexes: application to ubiquitin binding. *J Mol Biol* **375**, 1416-1433 (2008).
27. M. Biasini *et al.*, SWISS-MODEL: modelling protein tertiary and quaternary structure using evolutionary information. *Nucleic Acids Res* **42**, W252-258 (2014).
28. J. Yang *et al.*, The I-TASSER Suite: protein structure and function prediction. *Nat Methods* **12**, 7-8 (2015).
29. R. Guerois, J. E. Nielsen, L. Serrano, Predicting changes in the stability of proteins and protein complexes: a study of more than 1000 mutations. *J Mol Biol* **320**, 369-387 (2002).
30. J. Schymkowitz *et al.*, The FoldX web server: an online force field. *Nucleic Acids Res* **33**, W382-388 (2005).
31. J. W. Schymkowitz *et al.*, Prediction of water and metal binding sites and their affinities by using the Fold-X force field. *Proc Natl Acad Sci U S A* **102**, 10147-10152 (2005).
32. E. Boura *et al.*, Solution structure of the ESCRT-I and -II supercomplex: implications for membrane budding and scission. *Structure* **20**, 874-886 (2012).
33. E. Boura *et al.*, Solution structure of the ESCRT-I complex by small-angle X-ray scattering, EPR, and FRET spectroscopy. *Proceedings of the National Academy of Sciences of the United States of America* **108**, 9437-9442 (2011).
34. D. Chalupska *et al.*, Structural analysis of phosphatidylinositol 4-kinase IIIbeta (PI4KB) - 14-3-3 protein complex reveals internal flexibility and explains 14-3-3 mediated protection from degradation in vitro. *J Struct Biol* **200**, 36-44 (2017).
35. D. Chalupska *et al.*, Phosphatidylinositol 4-kinase IIIβ (PI4KB) forms highly flexible heterocomplexes that include ACBD3, 14-3-3, and Rab11 proteins. *Scientific Reports* **9**, 567 (2019).
36. J. Steinkuhler *et al.*, Membrane fluctuations and acidosis regulate cooperative binding of 'marker of self' protein CD47 with the macrophage checkpoint receptor SIRPalpha. *J Cell Sci* **132**, (2018).

### *Supplementary Results*

**Photon distribution analysis (PDA) for estimation of population sizes.**

In order to validate the PDA analysis as a way to estimate binding modes population fractions, we created artificial data sets using single binding mode mutants data. Since the mutants only bind in one of the two possible conformations, artificial data sets with known B1 and B2 fractions can be generated using experimental data. For example, 1000 bursts of M1 mutant can be mixed with 2000 bursts of M2, creating an artificial population in which 1/3 of the whole population is in the binding mode 1. As in the main text, we use here the fraction of binding mode 1 population to describe the population (the rest of the population is in the B2 state)

Using real data from mutants SB1 and SB2 of dockerin celS, we created an artificial data set *per* each cohesin from *C. thermocellum* (cohesin 1, 2 and 7) and analyzed them by PDA to estimate the fraction of each subpopulation (*1-4*). Since each set includes between 12-14 artificial populations with known subpopulation fractions, we can address the suitability of the PDA analysis to recover the subpopulation fractions of our samples.

Due to the stochastic nature of fluorescence emission, any fluorescence signal shows a distribution around its mean value, that is what is called the shot noise. PDA takes into account this stochastic nature and it is able to fit the experimental FRET distribution. For a single FRET population, the shape, mean and width of the distribution can be fitted using the distance between dyes (R) as the only fitting parameter. Generally, the FRET distribution is better fitted using a Gaussian distribution of distances centered at R and with standard deviation σ. Here, we used the implementation of PDA included in the free software package PAM (*3, 4*), and the correction parameters for detection efficiency, cross talk, direct excitation and background were included in every fit.

Three fitting parameters are required *per* each FRET state, a distance between dyes (R), a standard deviation of distances around R (σ), and finally an amplitude. Firstly, we fitted the original experimental populations of single binding mutants to obtain the best fit of R and σ values (Fig. SR1). In order to account for residual binding of the mutants in the alternative binding mode, SB1 and SB2 data were fitted together with R and σ acting as global parameters for both distributions. The retrieved values of R and σ were used as fixed values for the fitting of the artificial data sets, in such a way, only the fractions of subpopulations are the free fitting parameters.

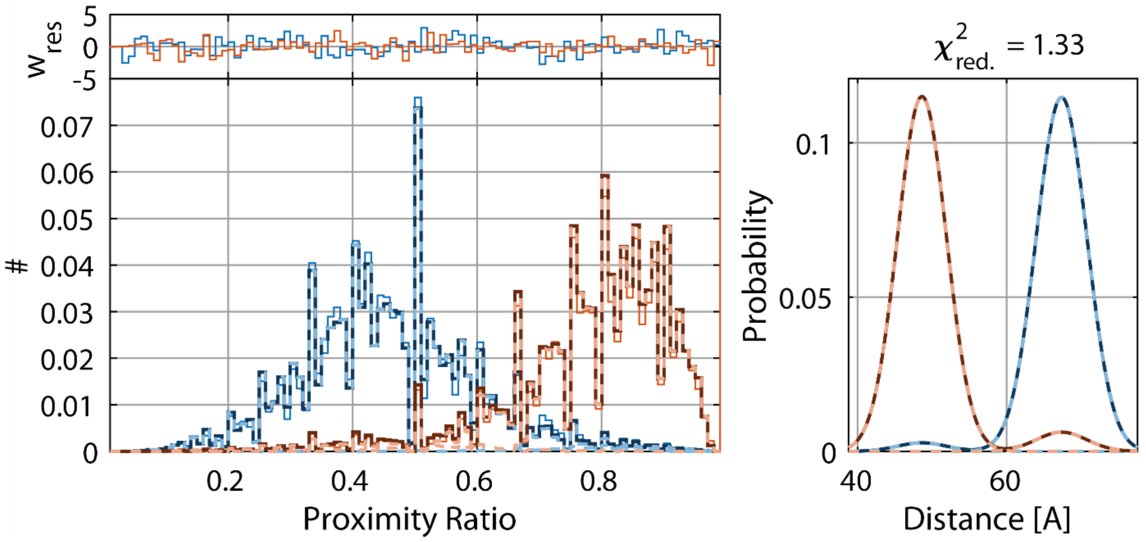

**Figure SR1.** PDA fitting of c2A-CelS_SB1 and c2A-CelS_SB2. In the left panel the predicted proximity ratio distribution in dashed line is superimposed to the experimental proximity ratio distribution. On the right panel, the distribution of fitting distances is shown.

As an example, we describe here the case of c2A artificial data set (fig SR2). A plot of the recovered fractions against expected values shown a very strong linear correlation (Pearson's r 0.999) but with a significant deviation from a line of slope 1 (fitting model y= a + b*x, a=0.908 ± 0.08, b= 0.048 ± 0.004, errors represented by standard error of the fitting). From the PDA fit and the FRET histograms, we noticed that the mutants retain residual binding in the alternative conformation, 2.3% for M1 mutant binding in B2 conformation and 5.3% for M2 binding in B1 conformation (values obtained in the initial PDA fitting of the mutants). We concluded that this might cause the observed deviation and we proceeded to correct the expected fractions for the residual binding. As an example, a population made out of 1506 bursts of SB1 and 3351 bursts of SB2 was corrected as: Nº of Bursts of SB1= (1506 - 1506*0.023 + 3351 *0.053), Nº of Bursts of SB2= (3351 -3351*0.053 + 1506 *0.023). Linear fitting of the PDA's recovered fractions *vs* the corrected fractions show a very good agreement with a line of slope 1 (y= a + b*x, a=0.984 ± 0.008, b= -0.004 ± 0.04, Pearson's r 0.9996). We repeated the procedure for all the three artificial data sets and the results are compiled in table S1. Excellent linear correlation was observed for all data sets (fig. SR3 and table SR1) and the mean difference between the PDA´s recovered values and the expected fraction range between: 0.004-0.012. The mean value of deviation for all the data sets was 0.007.

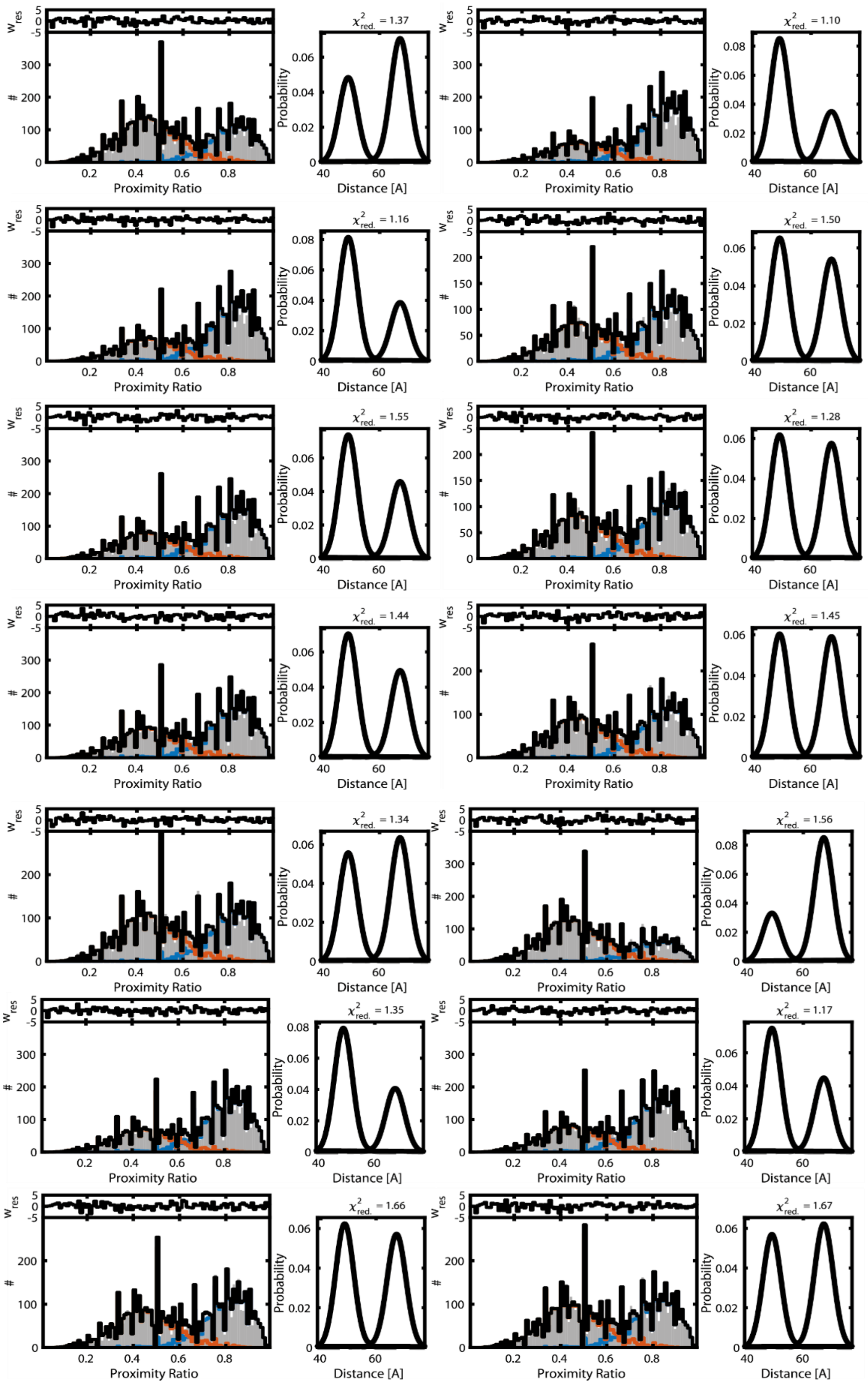

**Figure SR2**. Fitting of the whole artificial c2A data set. Fourteen artificial samples with different subpopulation ratios were fitted simultaneously. For every subpanel, the predicted proximity ratio distribution (dashed)

superimposed to the experimental distribution (solid) can be found on the left while the distribution of fitting distances are shown on the right.

We also used these data sets to study the impact of the elevated degrees of freedom introduced by six free fitting parameters on the determination of subpopulation fractions. Our main concern was to avoid significant deviations in the analysis of our real samples. In our previous analysis, we eluded the problem by fixing the R and σ value to those retrieved in the fitting of SB1 and SB2 mutants (see above). We decided not to use the same approach for the wild type CelS and CelA dockerin samples. Although good fitting was achieved in the case of the dockerin CelS for all the three cohesin-dockerin complexes studied, we got less accurate fitting when CelA wild type dockerin was fitted with the R and σ value of the mutants, likely because of small structural deviations from wild type.

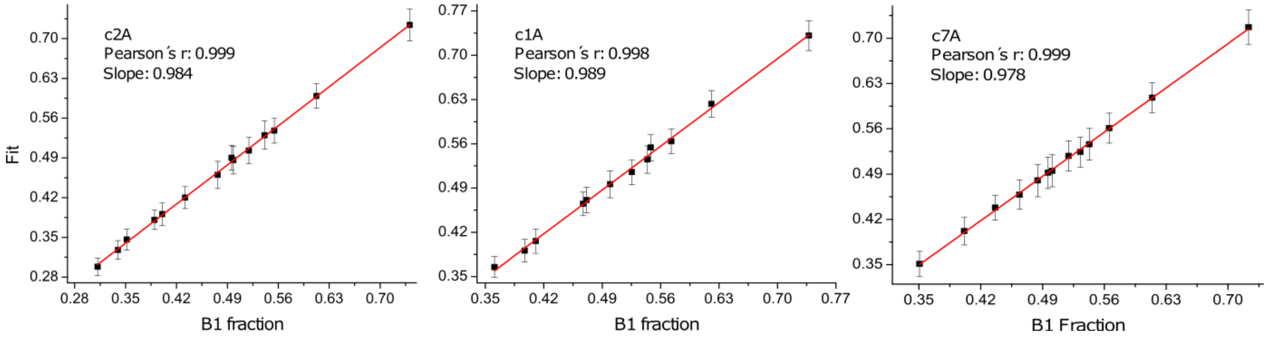

**Figure SR3** Plot of expected B1 fraction *vs* fraction recovered by PDA fit. Excellent linear correlation is observed for the three artificial data set.

We addressed the issue by fitting globally the six parameters for the whole artificial data set. This strategy gave good results since the linear correlations and mean deviation from the expected fraction values are very similar to those observed above (see Table SR2). Nevertheless, we observed that the obtained R and σ deviated slightly in some cases (see for example the case of the c1A cohesin data set, table SR2), and the deviation from the expected subpopulation fraction is generally larger. These small deviations were probably introduced by the extra degrees of freedom. We decided to incorporate the constrained fitting provided by the mutants in an alternative way. Starting with a population of roughly equal fraction of subpopulations, we split the FRET histogram in two, right in the point where the minimum between the high and low FRET was located. These high and low FRET populations were also included in the fitting among with the data set and the parameter fitted globally. With this approach, we reached better agreement with the R and σ values previously retrieved and a slightly smaller deviation respect to the expected values (see table SR2).

| Artificial set | slope | Error | intercept | Error | Pearson's r | Mean deviation from expected value |
|---|---|---|---|---|---|---|
| c2A Dock celS | 0.984 | 0.008 | -0.004 | 0.004 | 0.999 | 0.012 |
| c1A Dock celS | 0.989 | 0.018 | 0.0009 | 0.009 | 0.998 | 0.006 |
| c7A Dock celS | 0.978 | 0.008 | 0.007 | 0.004 | 0.999 | 0.004 |

**Table SR1.** Linear fit of expected B1 fraction *vs* PDA fitted fractions. Error stands for 95% confidence intervals.

Overall, and regardless of the strategy followed, the PDA analysis seems to be robust enough. Provided they are fitted globally, the flexibility introduced by the six free parameters in the fitting doesn´t affect significantly the recovered subpopulations fractions. Nevertheless and since the restrained fit provide better result, we systematically used it in the PDA analysis of our cohesin-dockerin complexes.

| Data Set | Method | R1 | Sigma1 | R2 | Sigma2 | Mean $\chi^2_{red}$ | Mean deviation |
|---|---|---|---|---|---|---|---|
| c2A Dock celS | Mutant values | 67.38± 0.54 | 3.39 ± 1.35 | 48.66 ± 0.48 | 3.28 ± 0.72 | 1.40 | 0.012 |
| | Free fitting | 67.65 ± 0.53 | 2.31 ± 1.44 | 48.88 ± 0.47 | 3.43 ± 0.72 | 1.34 | 0.027 |
| | Restrained fit | 67.35 ± 0.54 | 3.11 ± 1.38 | 48.59 ± 0.47 | 2.97 ± 0.72 | 1.49 | 0.012 |
| c1A Dock celS | Mutant values | 70.66 ± 0.35 | 3.33 ± 1.05 | 48.08 ± 0.32 | 3.24 ± 0.50 | 1.19 | 0.006 |
| | Free fitting | 70.76 ± 0.35 | 2.06 ± 1.1 | 48.33 ± 0.31 | 3.57 ± 0.49 | 1.12 | 0.014 |
| | Restrained fit | 70.61 ± 0.35 | 2.83 ± 1.08 | 48.09 ± 0.32 | 3.13 ± 0.50 | 1.22 | 0.005 |
| c7A Dock celS | Mutant values | 74.64 ± 0.55 | 3.16 ± 1.45 | 53.47 ± 0.39 | 2.63 ± 0.59 | 1.49 | 0.004 |
| | Free fitting | 74.47 ± 0.55 | 2.99 ± 1.45 | 53.32 ± 0.39 | 2.77 ± 0.59 | 1.46 | 0.003 |
| | Restrained fit | 74.39 ± 0.55 | 3.46 ± 1.45 | 53.22 ± 0.39 | 2.55 ± 0.59 | 1.57 | 0.004 |

**Table SR2**. Comparison of the three different approaches used to recover the subpopulation ratios by PDA analysis. Two FRET states were used for fitting. The fitting parameter, mean reduced $\chi^2$ and mean deviation of the whole data set from expected B1 ratio are shown.

**Molecular dynamics simulations**

*The Fold-X method*

As we stated in the methods section, we benchmarked our new MC Fold-X scheme studying first the cohesin-dockerin pair c2A-Xyn10B, analyzed in (*5*) by our previous Fold-X approach. An example of a Monte Carlo trajectory is shown in Fig. SR4. It shows *ΔG* and RMSD as a function of the number of the Monte Carlo steps. The RMSD is calculated either from the B1 structure used in a given simulation (the red color) or the B2 one (the blue color). The lowest *ΔG* found is -41.9 kcal/mol and -41.6 kcal/mol for B1 and B1 modes respectively, indicating a similar character of the corresponding basins. No transitions from B1 to B2 are observed despite several events of dissociation of the cohesin-dockerin complex.

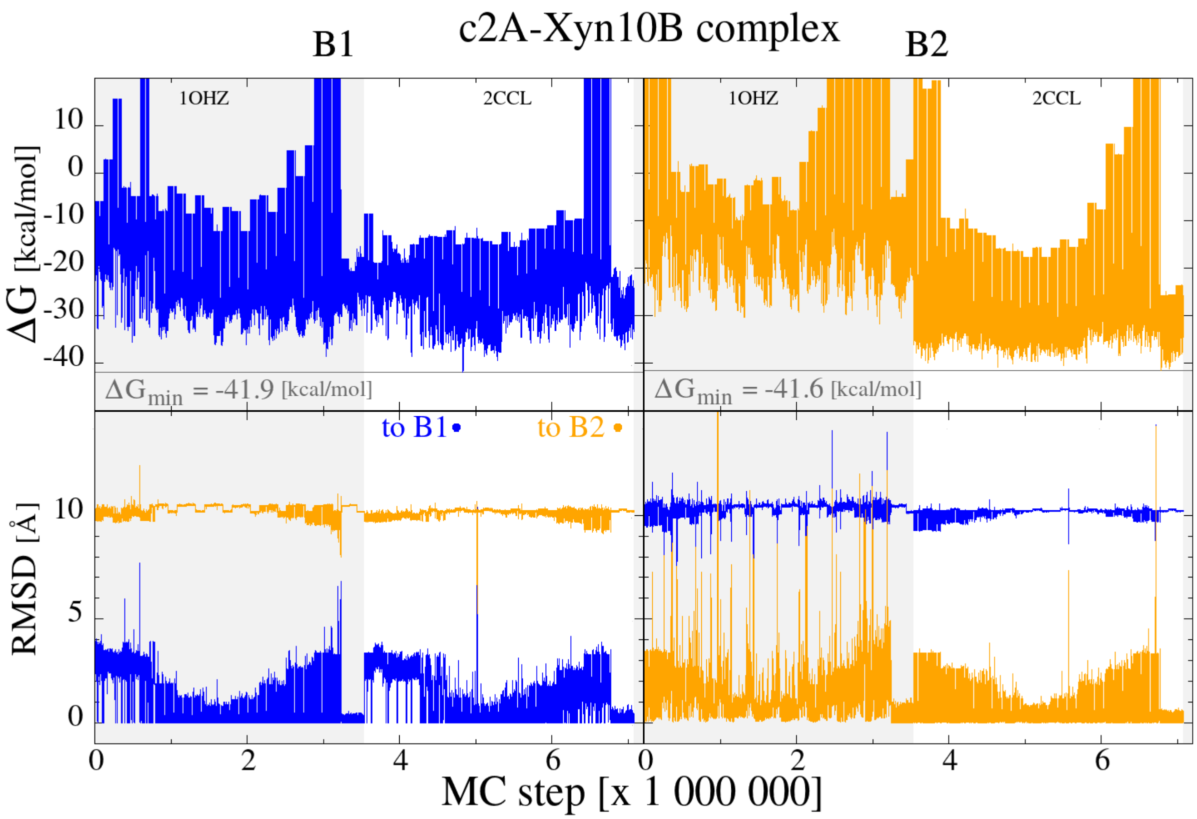

**Figure SR4** Example of a Monte Carlo trajectory for the c2A-Xyn10B complex. The top panels show the FoldX-derived free energy *ΔG* and the bottom panel – the RMSD distance to the B1 (in blue) and B2 (in orange) conformations (i.e. to 1OHZ and 2CCL). The left and right parts of the panels show structures in the B1 and B2 mode, respectively. The gray background on each graph shows simulations for cohesin-dcokerin complexes in which for 1OHZ serves as a template. The part without this background indicates simulations in which 2CCL is the template**.**

The results for the c2A-Xyn10B complex on $E_c$ dependence of the angle-, *Z*-, and structure-integrated $F_I$ and $F_{II}$, are shown in Figure SR5. The saturation values of *F* indicate that there is not much difference between the modes: -41.9 kcal/mol for B1 and -41.6 kcal/mol for B2. However, we estimate that the error of our method is of order 2 kcal/mol. Still, our simulations predict that the dual binding is present in this system. It should be noted that the new method is much more thorough and exhaustive. It yields $F_I$ and $F_{II}$ some 10 kcal/mol lower than those listed in the simplified previous approach (*5*). The free-energy minimum for B2 appears to be broader than the one for B1 in the sense that we get more points in the vicinity of the minimum for B2 than for B1.

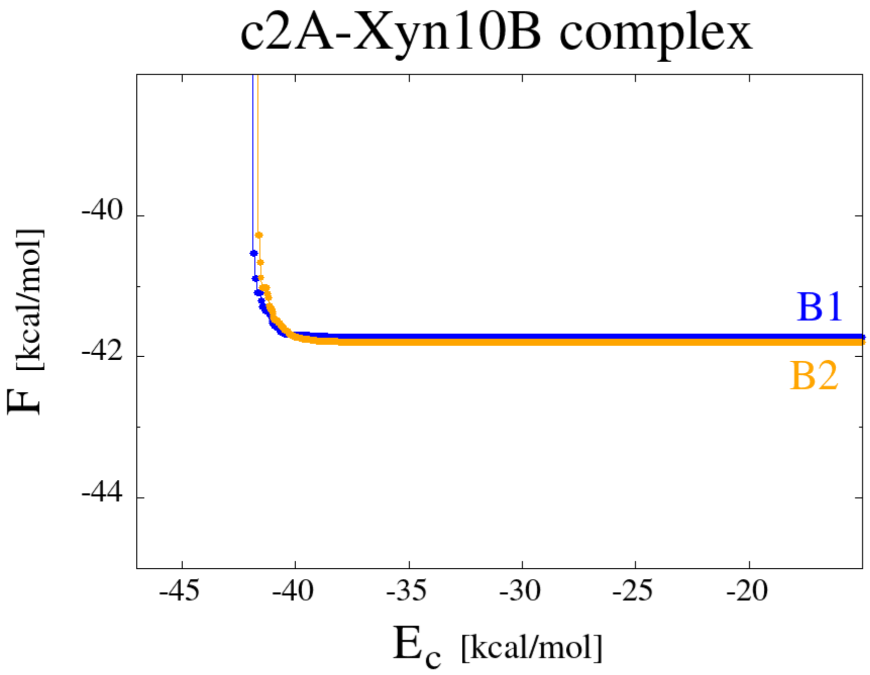

**Figure SR5** The dependence of the Monte-Carlo derived free energies on the cutoff energy $E_c$. The blue line corresponds to binding in B1 mode and the orange line to the B2 mode.

When we applied the MC Fold-X method to study the effect of the clasp mutation in the c2A-celS complex, we also analyzed the energy landscape of the WT and SB1 mutant complexes because they are good reference points. The first provides a reference to compare the effect of the removal of the clasp structure, and the second a situation where the methods should report a huge deviation toward binding in the B1 mode. Figure SR6 shows $F_I$ and $F_{II}$ as functions of $E_c$ for the three sequences. The results have been obtained within the uniaxial scheme. The binding energies appear to be a factor of 4 weaker than for the case of c2A-Xyn10B. However, they are, in fact, comparable, because these energies are measured relative to the energies in the dissociated state (about 20 kcal/mol for 1OHZ/2CCL and 10-15 kcal/mol for the 2MTE-case). The saturation values of $F_I$ and $F_{II}$ suggest existence of the dual binding mode for the WT but not for the B1 mutant, as expected. For the WT, the energy difference, $F_I - F_{II}$ is about −1.4 kcal/mol which indicates a slight preference for B1, but is still within the statistical error of ±2 kcal/mol of our method. The results obtained with the clasp mutant are thoroughly discussed in the main text.

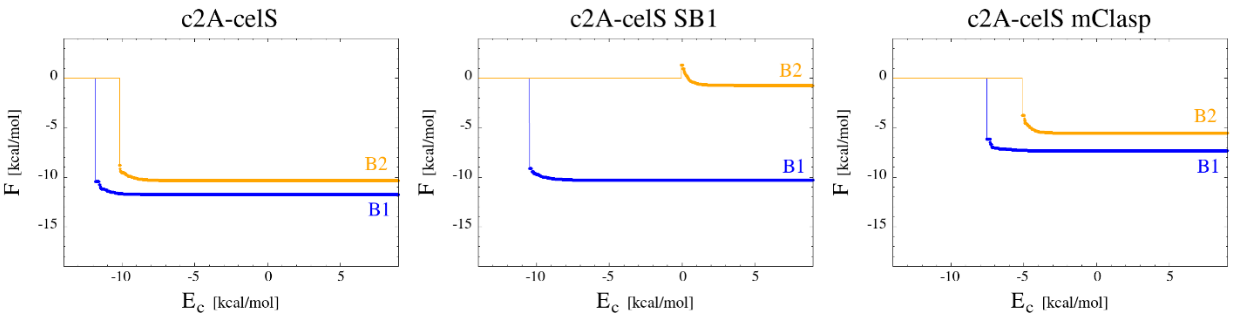

**Figure SR6** The $E_c$-dependence of the (angle-, *Z*-, and structure-integrated) free-energy for the c2A-CelS complex. The blue lines correspond to B1 and the orange lines to B2 mode. The panels from the left to the right correspond to the wild type complex, the single binding mode mutant SB1, and the clasp mutation respectively.

On the other hand, the problem with the uniaxial calculation is that it assumes that the proper axis of rotation of the dockerin with respect to the cohesin is the same as in the previous system. In a multi-axial calculation one can keep trying out new directions of the axis and making rotations around these various axes. Figure SR7 compares Monte-Carlo examples of trajectories obtained within the two schemes. The characteristic energies appear to be lowered by 3 – 4 kcal/mol when using the multiaxial approach. Nevertheless, the improved scheme would require a substantially larger computational effort and was abandoned.

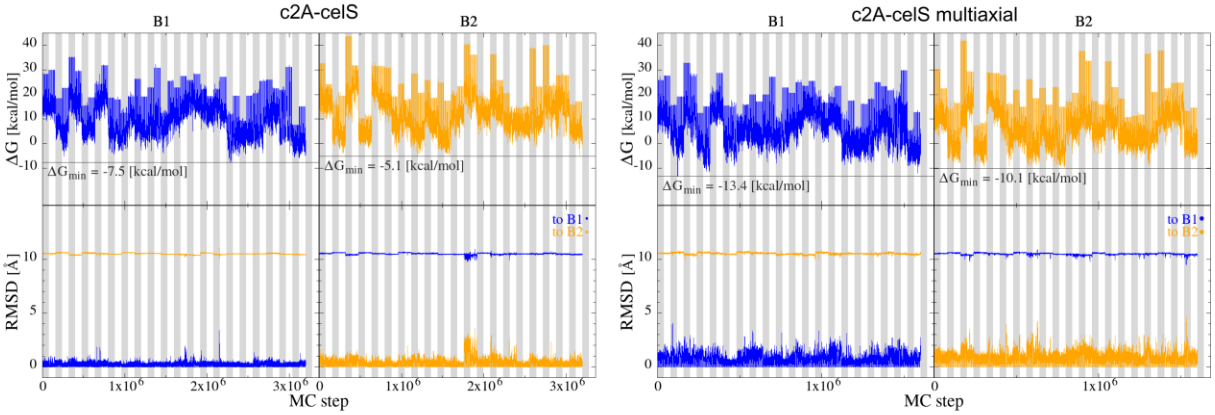

**Figure SR7** Monte Carlo trajectory for c2A-celS. The top panels: the energies *ΔG vs*. the Monte Carlo step count. The bottom panels: the corresponding values of RMSD counted either with respect B1 mode (in blue) or B2 (in orange). The two panels on the left are for the uniaxial simulation. Those on the right – for the multiaxial simulation.

**NMR experiments.**

We performed solution NMR experiments in order to check if the dockerin´s Leu 65 - Pro 66 peptide bond is in the *cis* conformation in the cohesin-dockerin complex. Although in the dockerin solution structure PDB:2MTE this peptide bond is in the *cis* conformation, this structure was solved in the absence of cohesin and therefore it remains unclear whether this bond remains in this conformation in the complex. As described below, we did not solve the solution structure of the dockerin bound to the cohesin, instead we used the exquisite sensitivity of the NMR chemical shifts to the local environment to probe that the Leu 65 - Pro 66 bond stays in the cis conformation in the cohesin-dockerin complex.

The solution structure of dockerin was elucidated previously by(*6*) under conditions similar to those employed here. In their structure (PDB: **2MTE)**, it can be seen that the Leu 65 - Pro 66 peptide bond is in the *cis* conformation. Moreover, the structure shows that Leu 65 packs on Tyr 57's aromatic ring and that Pro66's pyrrolidine ring is stacked on the aromatic ring of Tyr 5:

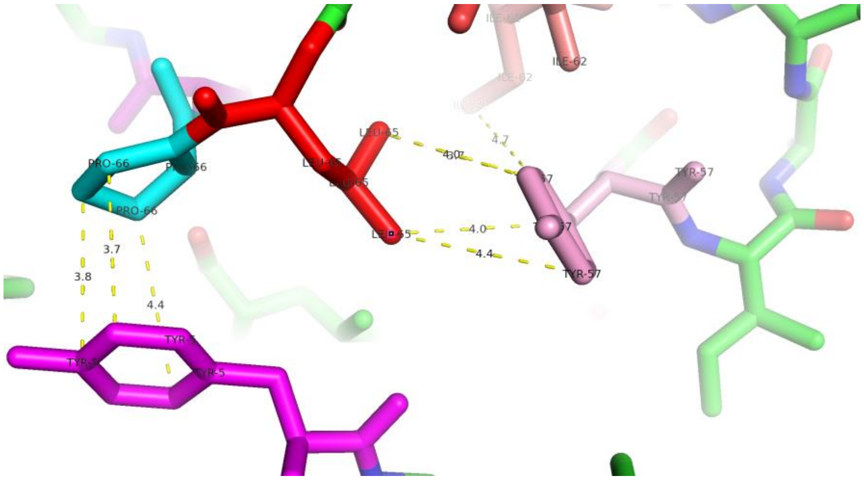

**Figure SR8**. Detail of the dockerin solution structure (PDB: 2MTE) showing Tyr 5 in magenta, Pro 66 in cyan, Leu 65 in red and Tyr 57 in pink. The distances between selected atoms, indicated by dashed yellow lines, are given in Angstroms.

NMR chemical shifts are highly sensitive to local environment. The chemical shifts of Pro 66 and its adjacent residues, as well as those of Tyr 5, would be expected to alter significantly if cohesin binding were to alter the conformation of Pro 66 or the isomeric state of the Leu 65 - Pro 66 peptide bond. We find that these dockerin chemical shifts are very similar in the absence or presence of 1.3 equivalents of cohesin, so this is good evidence that cohesin binding does not significantly alter the conformation of these residues (Table SR3). Our chemical shifts are also very similar to the values reported by (*6*) in the NMR database (BMRB file number: **25158**). In particular, the aliphatic $^{1}H$ signals of Leu 65, which are shifted strongly upfield due to the ring current effects of the Tyr 57 aromatic ring, are essentially the same in the absence or presence of 1.3 equivalents of cohesin (see Fig SR9):

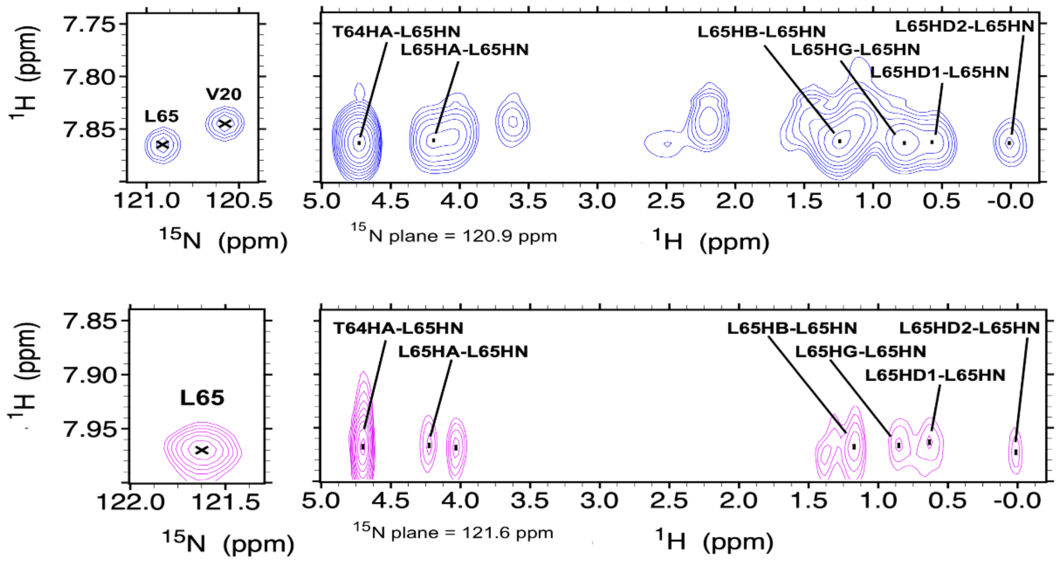

**Figure SR9** 2D $^{1}$H-$^{15}$N HSQC spectral region around L65 (small paneles on the left) of $^{15}$N-labeled dockerin without cohesin (blue) and with a 1.3 molar excess of unlabeled cohesin (magenta). Strips from 3D $^{1}$H-$^{15}$N-$^{1}$H HSQC·NOESY spectra (long panels on the right) showing NOESY crosspeaks to the HN of Leu65 of $^{15}$N-labeled dockerin in the absence (blue) and presence of a 1.3 molar excess of unlabeled cohesin (magenta).

Since such ring current effects are exquisitely sensitive to both angle and distance, this observation that Leu65's chemical shifts are so similar is particularly strong evidence that neither the local conformation of nearby residues nor the *cis* isomeric state of the Leu 65 - Pro 66 peptide bond are perturbed by cohesin binding.

| **Nucleus** | **δ Dockerin without Cohesin (ppm)** | **δ Dockerin with 1.3 eq Cohesin (ppm** | **δ Dockerin (ppm) from Chen et al.** |
|---|---|---|---|
| Lys 3 $^{1}$H, $^{15}$**N** | 8.30, **126.4** | 8.32, **126.4** | 8.31, **125.5** |
| Lys 3 $^{1}$Hα | 4.31 | 4.10 | 4.07 |
| Lys 3 $^{1}$Hβ | n.o.* | 1.31 | 1.34 |
| Lys 3 $^{1}$Hγ | n.o. | n.o. | 1.06, 0.87 |
| Lys 3 $^{1}$Hδ | n.o. | 1.31 | 1.35 |

| Nucleus | δ Dockerin without Cohesin (ppm) | δ Dockerin with 1.3 eq Cohesin (ppm | δ Dockerin (ppm) from Chen et al. |
|---|---|---|---|
| Lys 3 $^{1}$Hε | n.o. | n.o. | 2.72, 2.63 |
| Leu 4 $^{1}$H, $^{15}$**N** | 7.86, **127.8** | 7.81, **127.8** | 7.73, **127.4** |
| Leu 4 $^{1}$Hα | 4.49 | 4.49 | 4.48 |
| Leu 4 $^{1}$Hβ | 1.84, 1.24 | 1.83, 1.23 | 1.82, 1.21 |
| Leu 4 $^{1}$Hγ | 1.24 | 1.23 | 1.22 |
| Leu 4 $^{1}$Hδ | 0.83 | 0.82 | 0.87, 0.71 |
| Tyr 5 $^{1}$H, $^{15}$**N** | 7.99, 123.6 | 7.97, 1**23.6** | 7.82, **123.1** |
| Tyr 5 $^{1}$Hα | 3.58 | 3.57 | 3.62 |
| Tyr 5 $^{1}$Hβ | 2.41 | 2.40 | 2.45 |
| Gly 6 $^{1}$H, $^{15}$**N** | 8.90, **108.6** | 8.92, **108.7** | 8.92, **108.6** |
| Gly 6 $^{1}$Hα | 4.23, 3.16 | 4.23, 3.16 | 4.21, 3.17 |
| Asp 7 $^{1}$H, $^{15}$**N** | 8.16, **119.6** | 8.17, **120.4** | 7.98, **118.0** |
| Asp 7 $^{1}$Hα | 4.99 | 4.99 | 4.98 |
| Asp 7 $^{1}$Hβ | 3.09, 2.27 | 3.03, 2.19 | 3.00, 2.02 |
| Asp 63 $^{1}$H, $^{15}$**N** | 8.54, **119.6** | n.o. | 8.56, **119.7** |

| Nucleus | δ Dockerin without Cohesin (ppm) | δ Dockerin with 1.3 eq Cohesin (ppm | δ Dockerin (ppm) from Chen et al. |
|---|---|---|---|
| Asp 63 $^{1}H\alpha$ | 4.72 | 4.67 | 4.67 |
| Asp 63 $^{1}H\beta$ | 2.85 | 2.81 | 2.85, 2.67 |
| Thr 64 $^{1}H$, $^{15}$**N** | 7.40, **113.3** | 7.39, **113.2** | 7.41, **113.3** |
| Thr 64 $^{1}H\alpha$ | 4.73 | 4.70 | 4.71 |
| Thr 64 $^{1}H\beta$ | 4.06 | 4.03 | 4.04 |
| Thr 64 $^{1}H\gamma$ | 1.20 | 1.18 | 1.19 |
| Leu 65 $^{1}H$, $^{15}$**N** | 7.68, **121.0** | 7.97, **121.6** | 7.88, **121.0** |
| Leu 65 $^{1}H\alpha$ | 4.19 | 4.22 | 4.23 |
| Leu 65 $^{1}H\beta$ | 1.24, 0.58 | 1.20, 0.63 | 1.30, 0.52 |
| Leu 65 $^{1}H\gamma$ | 0.78 | 0.82 | 0.79 |
| Leu 65 $^{1}H\delta$ | 0.58, 0.02 | 0.63, 0.03 | 0.65, 0.01 |
| Pro 66 $^{1}H\alpha$ | 4.00 | 3.98 | 4.00 |
| Pro 66 $^{1}H\beta$ | 1.62 | 1.56 | 1.64, 1.58 |
| Pro 66 $^{1}H\gamma$ | n.o. | n.o. | 1.34, 1.43 |
| Pro 66 $^{1}H\delta$ | 3.32, n.o. | 3.37, n.o. | 3.29, 2.63 |
| Tyr 67 $^{1}H$, $^{15}$**N** | 8.66, **122.7** | 8.69, **112.6** | 8.66, **122.6** |

| Nucleus | δ Dockerin without Cohesin (ppm) | δ Dockerin with 1.3 eq Cohesin (ppm | δ Dockerin (ppm) from Chen et al. |
|---|---|---|---|
| Tyr 67 $^{1}$Hα | 4.57 | 4.63 | 4.66 |
| Tyr 67 $^{1}$Hβ | 2.84 | 2.79 | 2.91, 2.77 |
| Lys 68 $^{1}$H, $^{15}$**N** | 8.14, **126.2** | 8.15, 126.3 | 8.11, **126.2** |
| Lys 68 $^{1}$Hα | 4.28 | 4.26 | 4.27 |
| Lys 68 $^{1}$Hβ | 1.81, 1.43 | 1.41 | 1.78, 1.44 |
| Lys 68 $^{1}$Hγ | 1.17 | n.o. | 1.17 |
| Lys 68 $^{1}$Hδ | n.o. | n.o. | 1.53 |
| Lys 68 $^{1}$Hε | n.o. | n.o. | 2.84 |

**Table SR3**. Table of chemical shifts. *n.o.: not observed

**Supplementary results references**

1. M. Antonik, S. Felekyan, A. Gaiduk, C. A. Seidel, Separating structural heterogeneities from stochastic variations in fluorescence resonance energy transfer distributions via photon distribution analysis. *J Phys Chem B* **110**, 6970-6978 (2006).
2. S. Kalinin, S. Felekyan, A. Valeri, C. A. M. Seidel, Characterizing Multiple Molecular States in Single-Molecule Multiparameter Fluorescence Detection by Probability Distribution Analysis. *The Journal of Physical Chemistry B* **112**, 8361-8374 (2008).
3. W. Schrimpf, A. Barth, J. Hendrix, D. C. Lamb, PAM: A Framework for Integrated Analysis of Imaging, Single-Molecule, and Ensemble Fluorescence Data. *Biophysical Journal* **114**, 1518-1528 (2018).
4. E. Sisamakis, A. Valeri, S. Kalinin, P. J. Rothwell, C. A. Seidel, Accurate single-molecule FRET studies using multiparameter fluorescence detection. *Methods Enzymol* **475**, 455-514 (2010).
5. M. Wojciechowski *et al.*, Dual binding in cohesin-dockerin complexes: the energy landscape and the role of short, terminal segments of the dockerin module. *Scientific Reports* **8**, 5051 (2018).
6. C. Chen *et al.*, Revisiting the NMR solution structure of the Cel48S type-I dockerin module from Clostridium thermocellum reveals a cohesin-primed conformation. *J Struct Biol* **188**, 188-193 (2014).

### *Supplementary figures and Table*

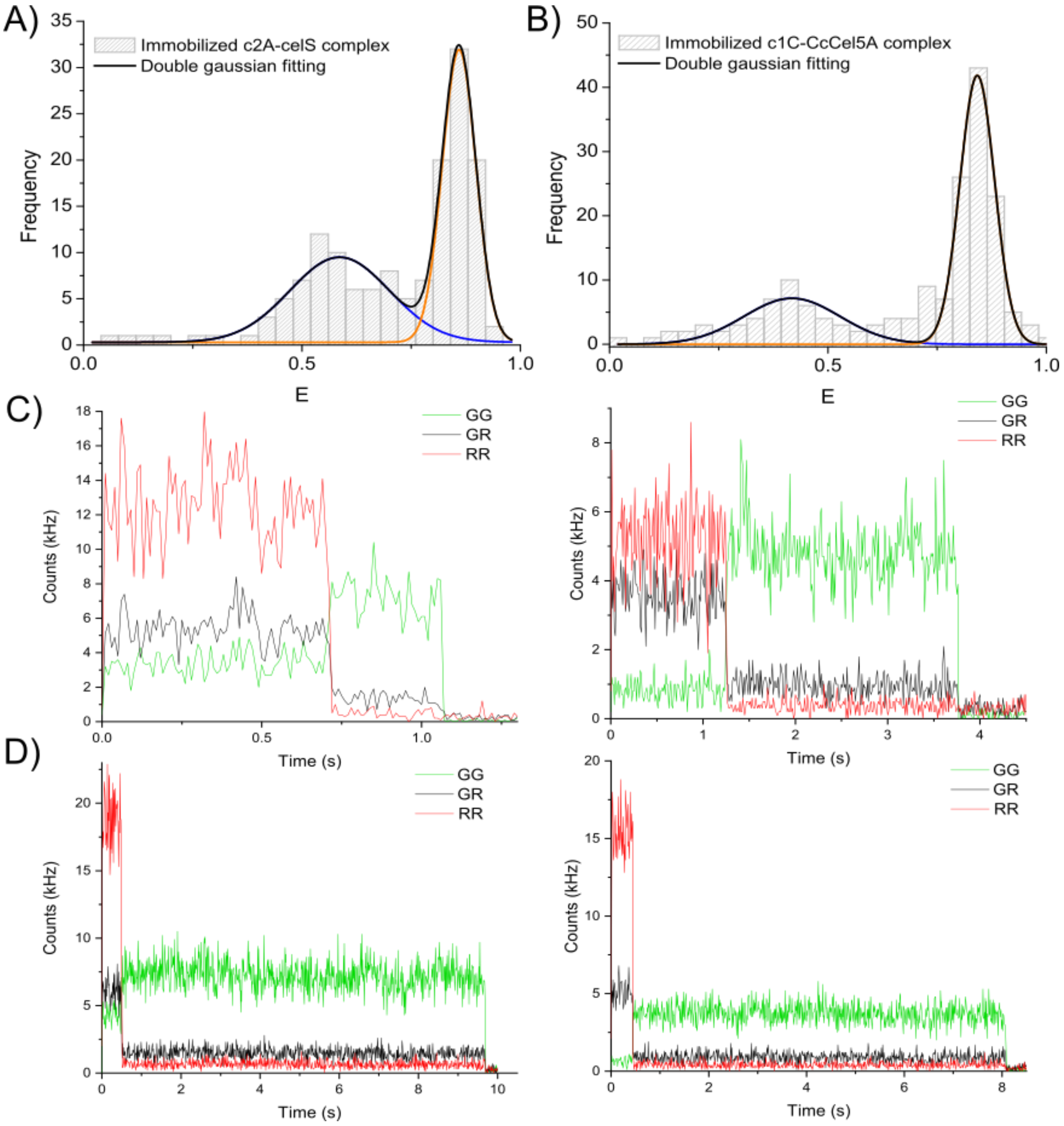

**Figure S1**. **Single binding mode detection in surface-immobilized complex**. The cohesin modules were selectively biotinylated at the N-terminus, and the whole cohesin-dockerin complex immobilized in neutravidin-functionalized surfaces. **A)** In good agreement with free diffusing molecules, the FRET histogram of immobilized c2A-celS complexes shows a bimodal distribution of low and high FRET populations as well. The bimodal distribution was fitted with two Gaussians (centered at blue: 0.58 ± 0.23 and orange: 0.86 ± 0.08). Error report for standard deviation. **B)** Both populations were also identified in surface-immobilized c1C-CcCel5A complexes. Black line represents Gaussian fitting of the bimodal distribution with two Gaussian functions (centered at blue: 0.42 ± 0.22 and orange: 0.84 ± 0.08). **C)** *C. thermocellum* c2A-celS traces. A low FRET trace is shown on the left, and a high FRET on the right. **D)** *C. cellulolyticum* c1C-Ccel5A single-molecule traces. Similarly, low and high FRET traces are represented on the left and right respectively. GG stands for green excitation and green emission channel, GR for green excitation and red emission channel, and RR for red excitation and red emission channel.

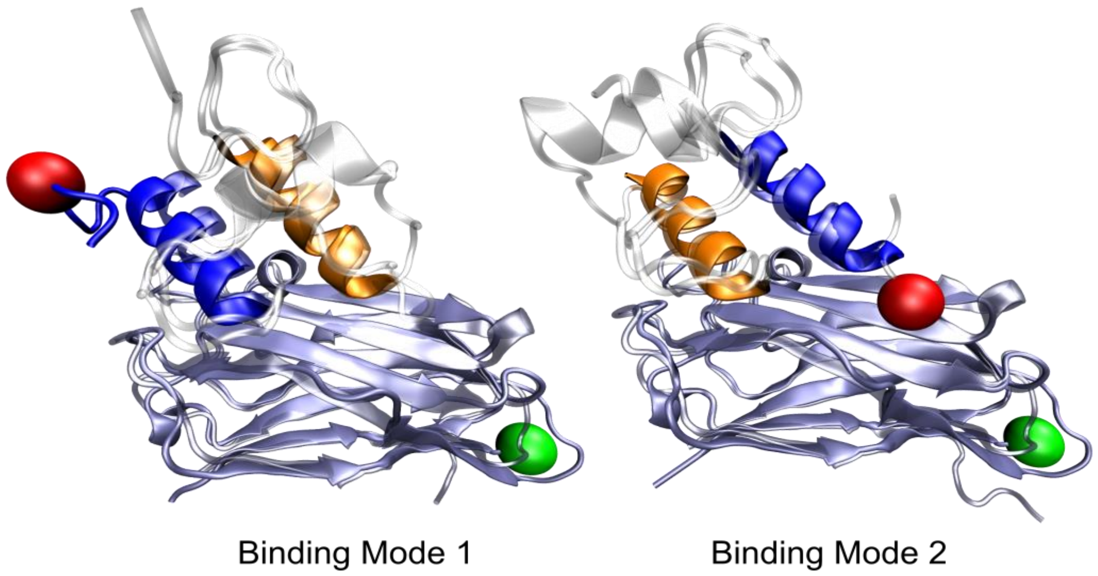

**Figure S2 Structural similarities between *C. thermocellum* and *C. cellulyticum* cohesin-dockerin** complexes. *C. thermocellum* and *C. cellulyticum* complexes follows the same color code as in the main text except the *C. thermocellum* complexes are shown in transparent colors. PDB codes for the *C. cellulyticum* structure in B1 modes is 2VN6, and 2VN5 for B2 mode. PDB codes for the *C. thermocellum* structure in B1 modes are 1OHZ, and 2CCL for B2 mode.

A)

```
              Dockerin CelS                Dockerin CelA

H1      stdavalkryvlr     H1      stdltmlkryllk
H3      stdlgiilkryilk    H3      ssdmtilkrylik
        ***   ****:*:             *:*:*:*****:*
```

B)

```
c2A    MDGVVVEIGKVTGSVGTTVEIPVYFRGVPSKGIANCDFVFRYDPNVLEIIGIDPGDIIVD 60
c7A    MGAVRIKVDTVNAKPGDTVRIPVRFSGIPSKGIANCDFVYSYDPNVLEIIEIEPGELIVD 60
       *..* :::..*... * **.*** * *:***********: ********* *:**::***

c2A    PNPTKSFDTAIYPDRKIIVFLFAEDSGTGAYAITKDGVFAKIRATVKSSAPG---YITFD 117
c7A    PNPTKSFDTAVYPDRKMIVFLFAEDSGTGAYAITEDGVFATIVAKVKSGAPNGLSVIKFV 120
       **********:*****.*****************:*****.* *.***.**.    *.*

c2A    EVGGFADNDLVEQKVSFIDGGVNV 141
c7A    EVGGFANNDLVEQKTQFFDGGVNV 144
       ******:*******..*:******
```

C)

```
c1C    -MDSLKVTVGTANGKPGDTVTVPVTFADVAKMKNVGTCNFYLGYDASLLEVVSVDAGPIV 59
c7C    MGKELKVAVGTASGKAGDTVTVPVTFADVATVGNVGTCNFYVTYDTNLLEVASVTPGSIV 60
         ..***:****.** **************.: ********: **:.****.**  * **

c1C    KNAAVNFSSSASNGTISFLFLDNTITDELITADGVFANIKFKLKSVTAKTTTPVTFKDGG 119
c7C    TNAAVNFSSSTSNGTISFLFLDNTITDQLIKTDGTFAEIKFKLKSVTAKTTTPVAFKDGG 120
       .*********:****************:**.:**.**:****************:*****

c1C    AFGDGTMSKIASVTKTNGSVTIDP 143
c7C    AFGDGTMAKIATVTKTNGSVTIDV 144
       *******:***:***********
```

D)

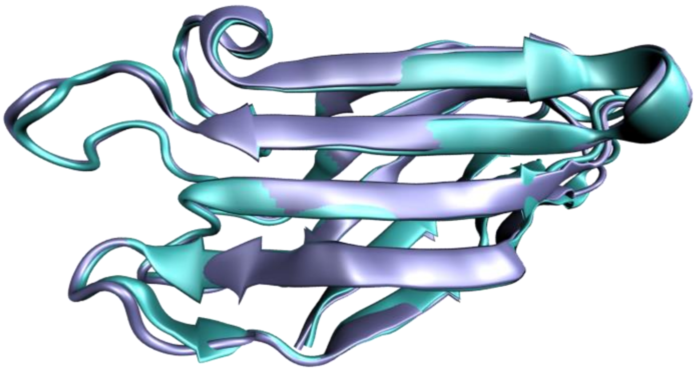

**Figure S3**. **Similarity between different cohesins and dockerins**. **A)** Sequence alignment between H1 and H3 helices of *C. thermocellum* celS and celA dockerins. The alignment shows the higher degree of similarity between helices in the case of celA dockerin, especially in the key positions 19, 20, 21 (H1) and 51, 52 and 53 (H3) that are highlighted in green. **B)** Sequence alignment of cohesin c2A and c7A from *C. thermocellum*. The sequence similarity between the cohesins is very high (73 % identity and 94 % similarity), particularly among those residues involved in contacts with the dockerin module (in green, 94 % identity and 100 % similarity). **C)** Sequence alignment of cohesin c1C and c7C from *C. cellulolyticum*. The sequence similarity between both cohesins is remarkably high (81 % identity and 96 % similarity), particularly among those residues involved in contacts with the dockerin module (in green, 100 % identity). **D)** Structural alignment of *C. thermocellum* cohesin c7A (dark blue, PDB code 1AOH) and c2A (cyan, PDB code 1OHZ). The alignment shows that the proteins' folds are highly similar.

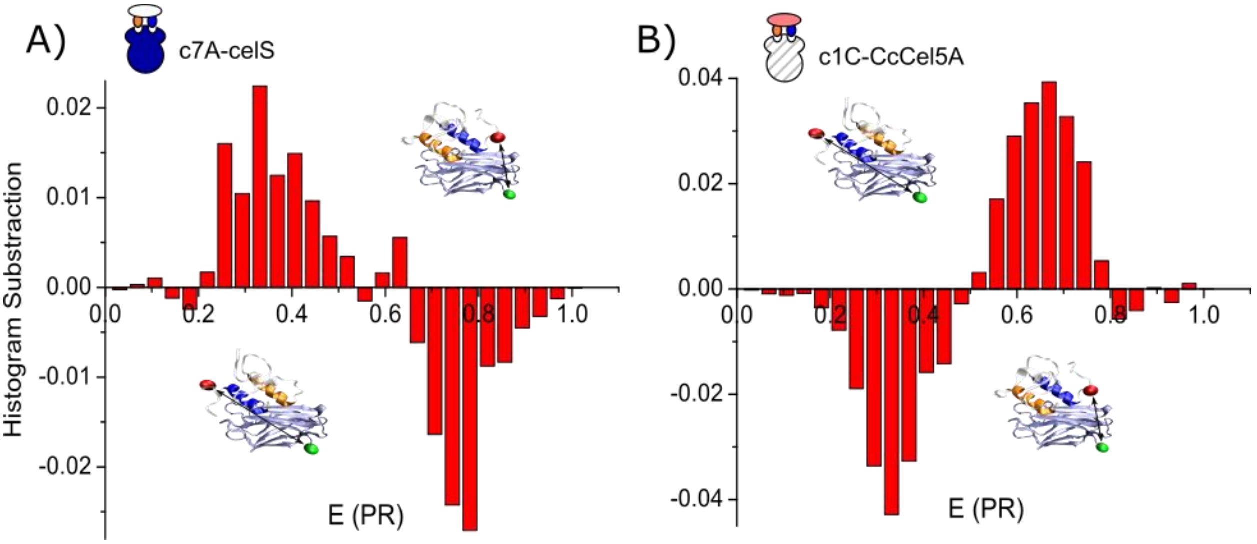

**Figure S4. Asymmetry analysis**. **A)** and **B)** different distribution of B1 and B2 subpopulations between cohesin-dockerin complexes. **A)** Plot showing the subtraction of a representative histogram of c7A-CelS complex (B1 fraction occupation of 0.58) and artificial population of single binding mode mutants displaying a B1 fraction of ≈ 0.50. **B)** Subtraction between c1C-Ccel5A and c7C-Ccel5A histograms showing the different distribution of binding modes among these *C. cellulolyticum* complexes. All histograms were area-normalized before the subtraction.

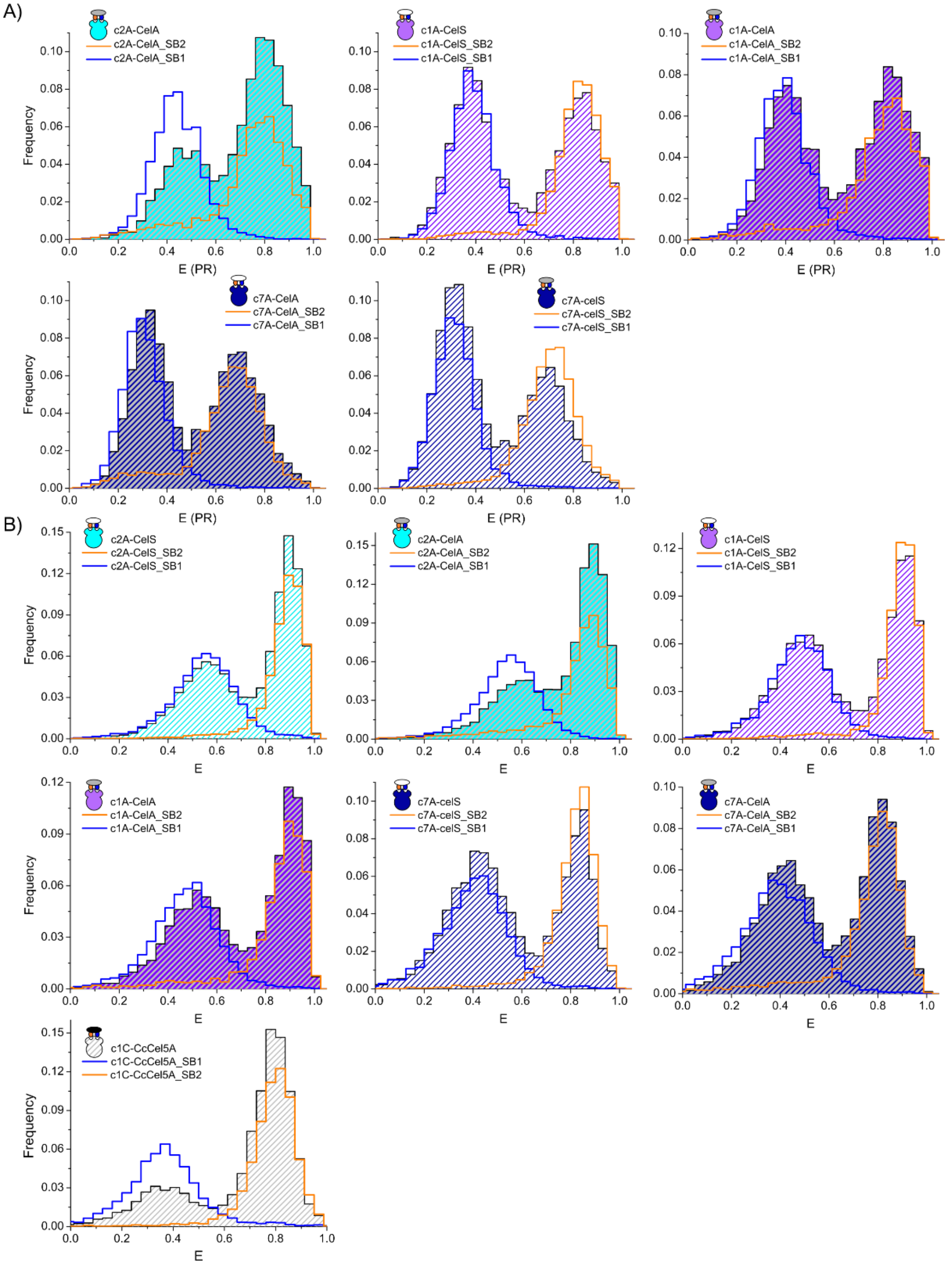

**Figure S5. smFRET histograms of cohesin-dockerin complexes**. **A)** Histograms of *C. thermocellum* complexes (striped histograms) are overlaid with those of single binding mode mutants (SB1 in blue and SB2 in orange), in order to confirm the identification of the low FRET and high FRET population as the B1 and B2 state. **B)** Fully corrected FRET histograms of *C. thermocellum* and *C. cellulolyticum* complexes. Overlaid on the complex histogram (striped histograms) the single binding mode mutants (SB1 in blue and SB2 in orange) are shown. Corrected and uncorrected histograms of *C. thermocellum´s* celA complexes show small structural deviations of SB mutants, but excellent agreement is observed for CelS and *C. cellulolyticum* c1C-CcCel5a complexes. All histograms were area-normalized to 1, SB mutants were normalized to a total area of 0.5 for comparison purposes.

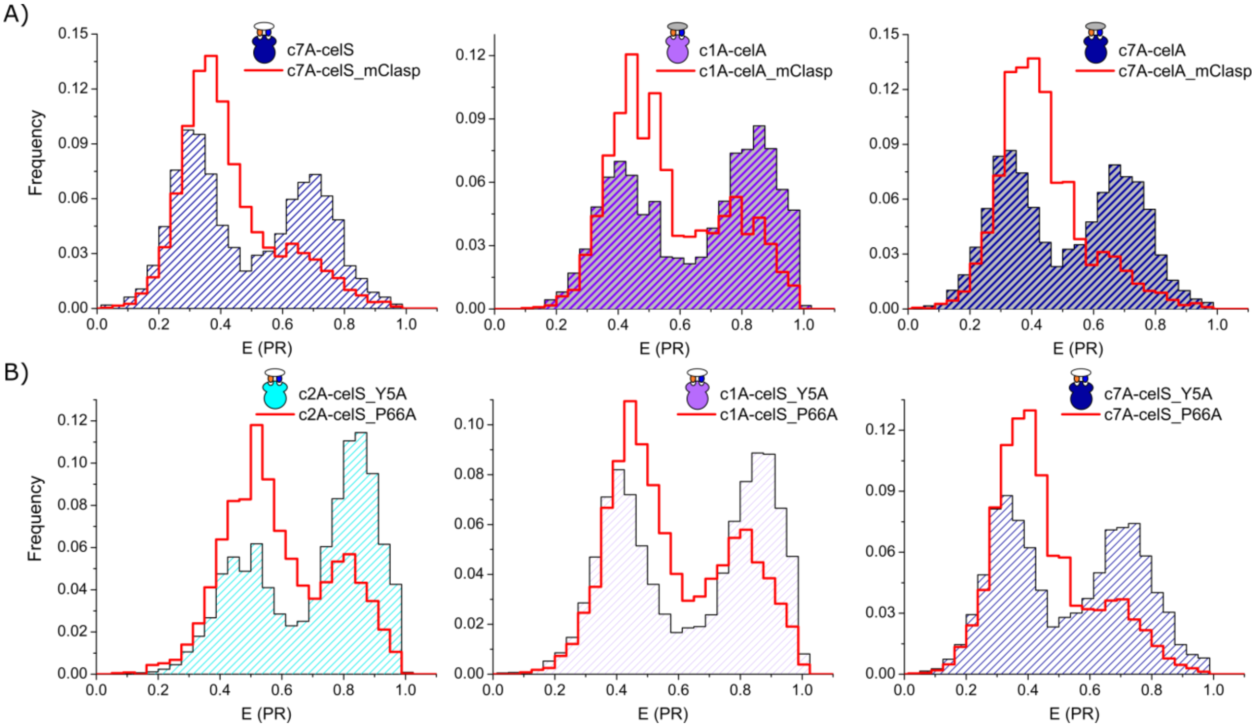

**Figure S6**. **FRET Histograms of *C. thermocellum* clasp mutant complexes**. **A)** A significant impact on the distribution of binding modes was observed for both dockerins, celS and celA and for all the complexes upon clasp mutation. Comparison of the clasp mutants (red solid line) with the wild type complexes (color-striped histograms) shows a consistent shift toward the B1 mode. **B)** The role of P66 in the clasp. In order to dissect the effect observed in *C. thermocellum* clasp mutants, we studied mutants of the individual amino/imino acid residues involved in the clasp (Tyr5 and Pro66). Surprisingly, complexes with a single mutation in Tyr5 (color-striped histograms) show a distribution of B1 and B2 subpopulation totally comparable with the wild type complexes (see Fig 2A, 3b and S5 and S6A)). In contrast, the mutant of Pro66 (red solid line) produces an obvious shift toward the B1 mode, even accounting for most of the shift observed in the clasp mutant for c2A and c7A complexes (see Fig 5C, and S6A)).

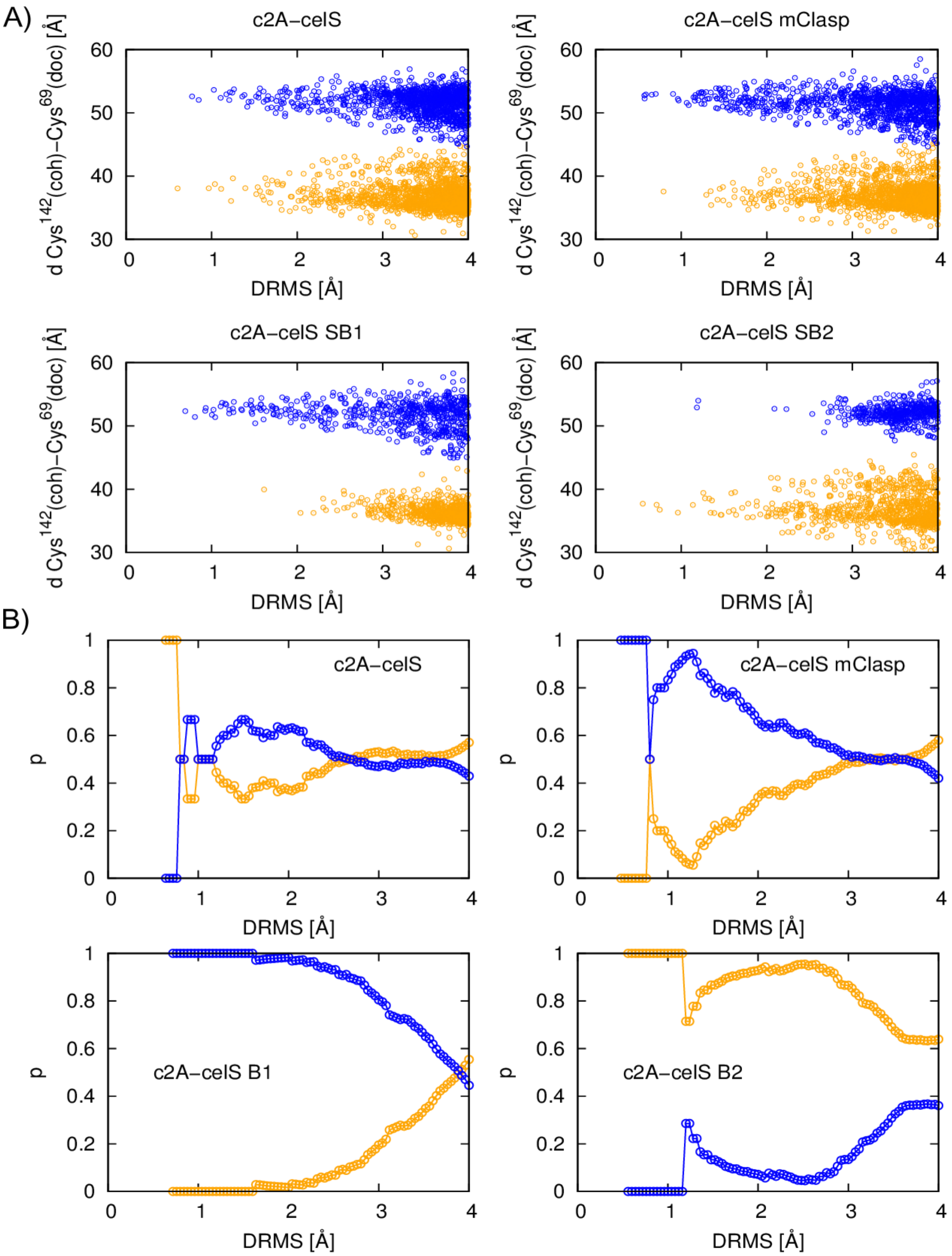

**Figure S7**. **Coarse-grained simulations**. **A)** The plots show the distance between the α-C atoms of $CYS^{142}$ in cohesin and $CYS^{69}$ in dockerin CelS *versus* DRMS(*S, $S_I$*) (blue data points) or DRMS(*S*, *$S_{II}$*) (orange). Each of the data points corresponds to a cohesin-dockerin structure *S* obtained from the CG simulations. Four systems are considered: the wild type, c2A-CelS, (top left), the clasp mutant, c2A-CelSmClasp, (top right), the SB1 mutant, c2A-CelS_SB1, (bottom left) and the SB2 mutant, c2A-CelS_SB2, (bottom right). **B)** Estimates for the probability of the cohesin binding the CelS dockerin in B1 mode (blue lines) or B2 mode (orange lines) as a function of the DRMS cutoff, d (see methods for a description of how this probability is calculated).

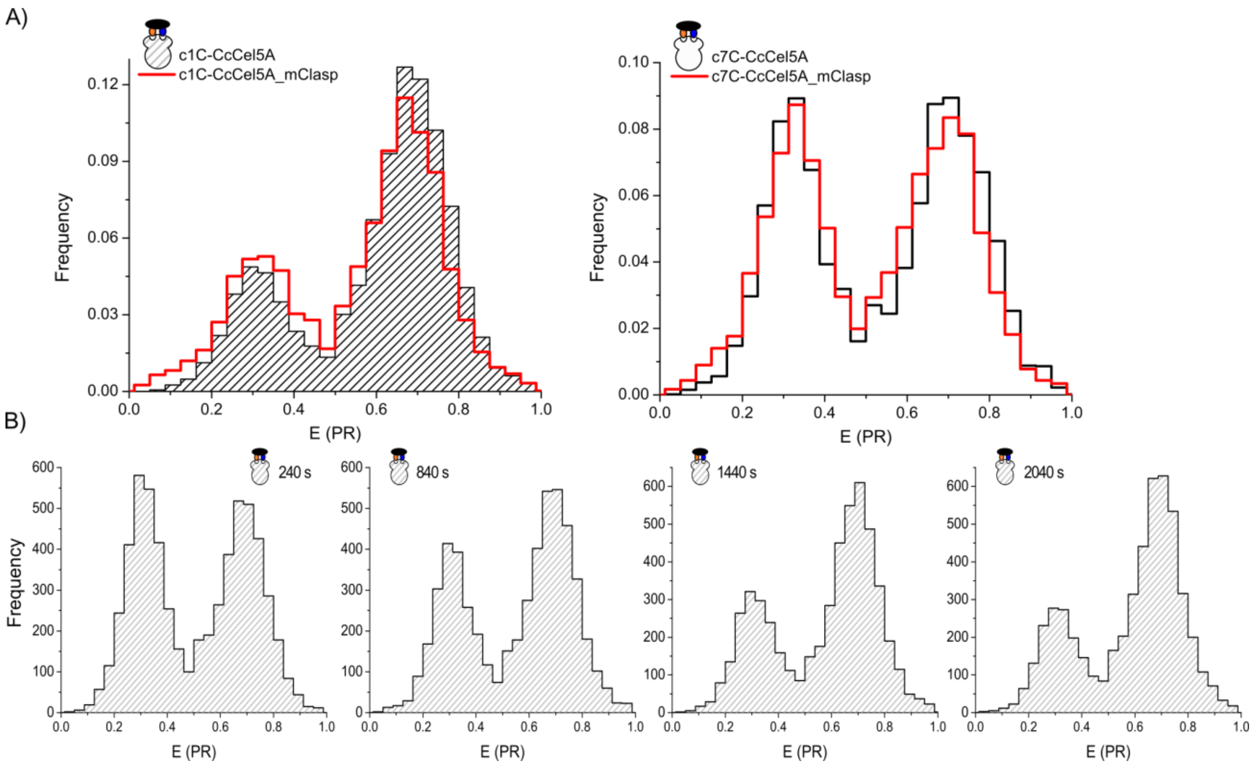

**Figure S8. Behavior of *C. cellulolyticum* complexes. A)** The effect in binding mode subpopulations for *C. cellulolyticum* clasp mutants. Although clasp mutation has a strong impact on the distribution of binding modes subpopulations in *C. thermocellum*, this is not the case for *C. cellulolyticum*. A mild enrichment in B1 was observed for c1C-CcCel5A complexes (B1 fraction of wild type 0.26 ± 0.02 *vs* 0.33 ± 0.03 of clasp mutant, left plot and Fig 5a), but no effect was present in the case of c7C-CcCel5A complexes (right, B1 fraction of wild type 0.45 ± 0.02 *vs* 0.5 ± 0.09 for clasp mutant). Clasp mutants are represented in red lines. **B)** Time series of wild type *C. cellulolyticum* smFRET histograms. Each histogram represents a time point on the evolution of the c1C-CcCel5A complex. The binding modes ratios evolved over time from a roughly equally distribution of binding modes subpopulations to the preeminence of B2 mode at equilibrium. All measurement were done at room temperature.

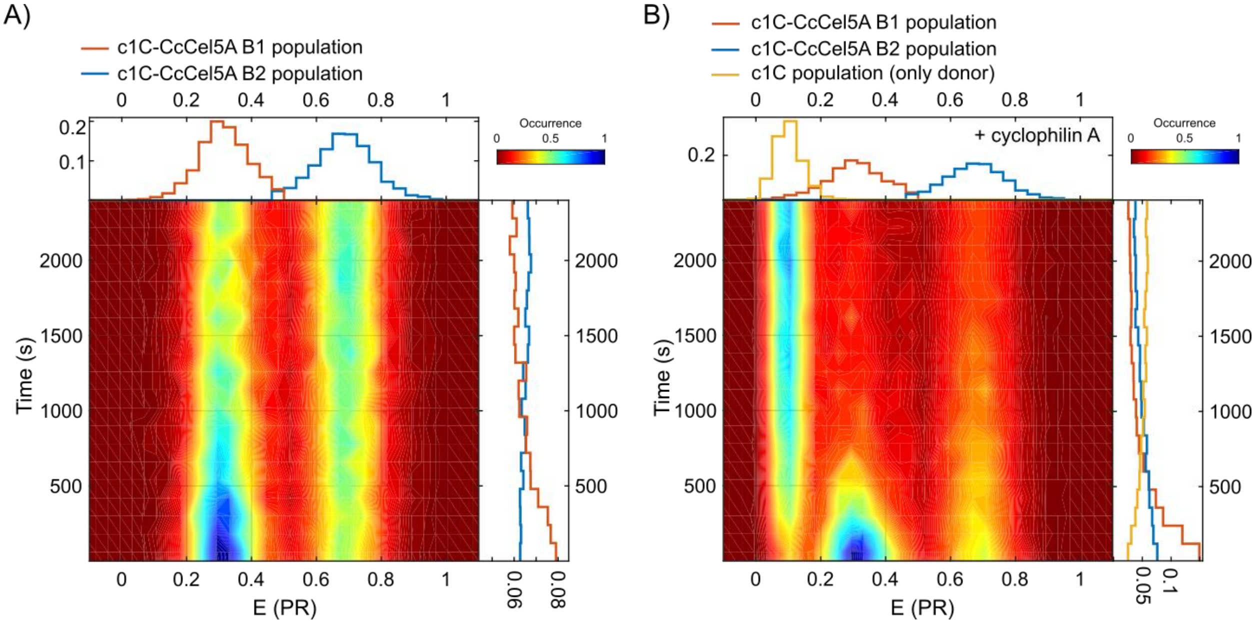

**Figure S9 Re-equilibrium process in c1C-Ccel5A samples**. 2D histograms of **A)** c1C-CcCel5A without cyclophilin and **B)** cyclophilin-treated c1C-CcCel5A complexes. FRET efficiency *vs* time is represented in the 2D histograms, on top of the 2D histogram, 1D FRET histogram is shown, and on the right the area-normalized frequency of detected bursts (each population normalized by 1). B1 (in blue), B2 (red) and only c1C cohesin population (yellow, only in **B)**) are plotted in the 1D histograms. The data shows that the kinetics behavior exhibit by the *C. cellulolyticum* complex is due to a re-equilibration process between binding modes that occurs upon dilution in the experimental conditions, rather than a direct interconversion between binding modes. Direct interconversion would produce a direct enrichment of the B2 population upon B1 depletion, which is not observed (see for example that the B1 and B2 population are independent during the first 800 seconds). Besides re-equilibration, with a visit to the pool of free molecules upon unbinding from one binding mode before binding in the second binding mode, fits well with the increase of free cohesin population that it is seen in the first moments of the dynamics (see first 800 s in **B)**).

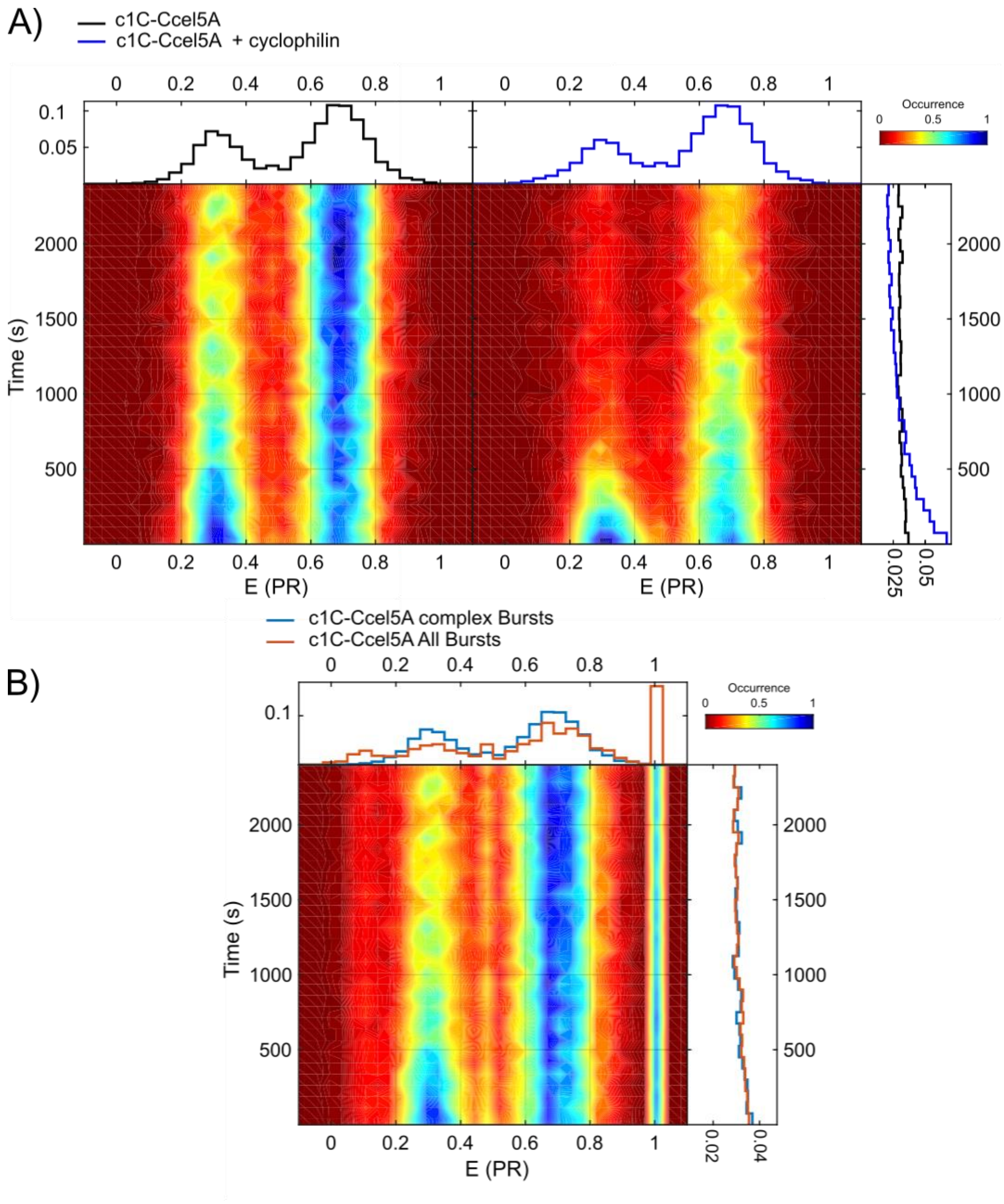

**Figure S10 Destabilization of c1C-Ccel5A complex upon proline isomerization**. 2D histograms of FRET efficiency *vs* time, on top of the 2D histogram the 1D FRET histogram is shown, and on the right the area-normalized frequency of detected bursts from all B1+B2 populations (each population normalized by 1). **A)** The total number of complex bursts rapidly decreased in the cyclophilin-treated samples (blue line), suggesting a general complex destabilization due to proline isomerization. It is important to notice that such a decay is not observed in the untreated samples (black line). **B)** Unspecific decay of c1C-Ccel5A complex. Frequency of all detected bursts (red line) and only complex bursts (blue line) are plotted. The slow population decay in c1C-Ccel5A complexes is undistinguishable from the decay observed in all bursts, suggesting that the decay is probably due to unspecific binding of the proteins to the chamber´s surfaces or sample photodamage.

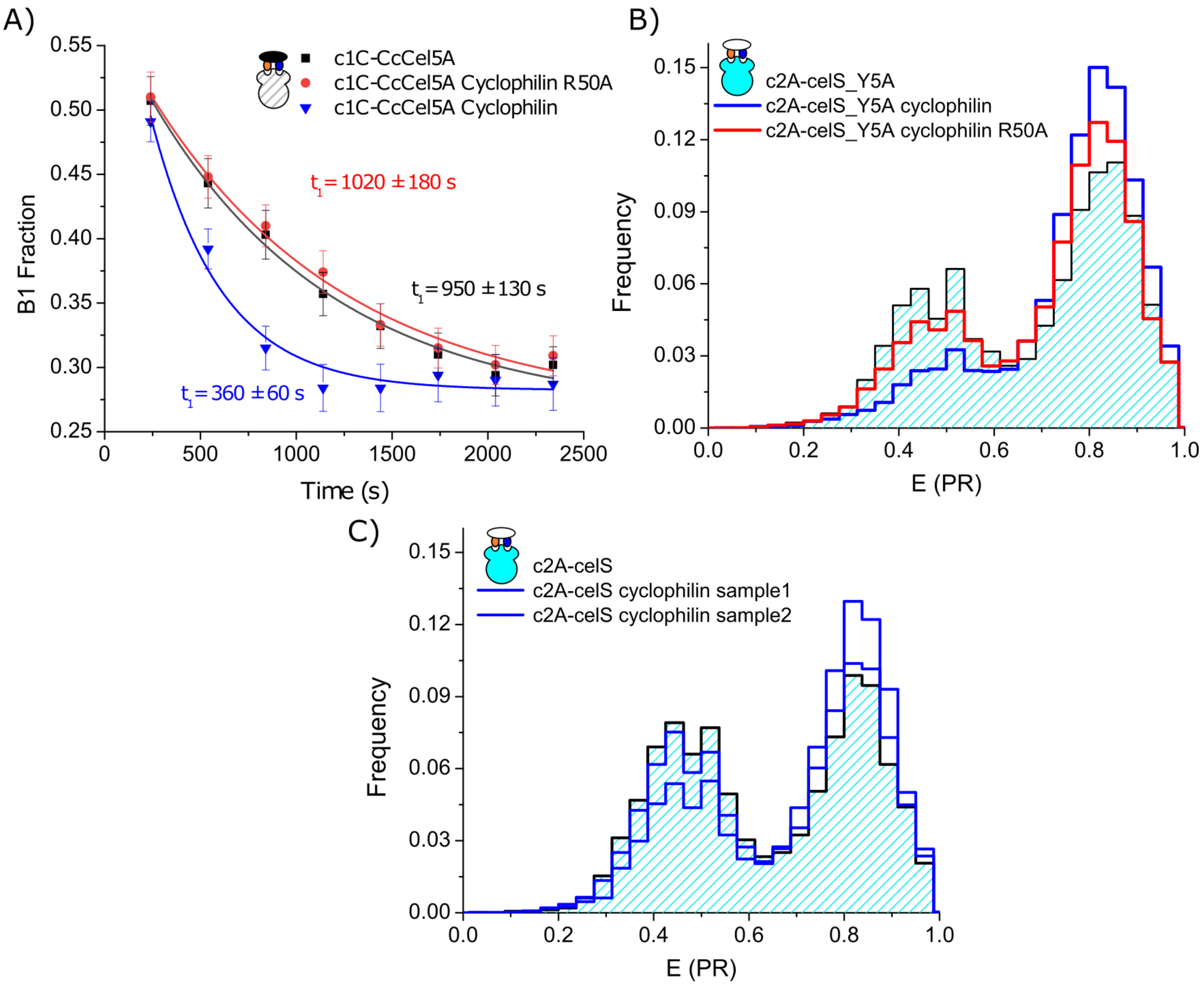

**Figure S11 Kinetics and population changes are due to proline isomerase activity. A)** In order to confirm the regulatory role of the *cis*/*trans* isomerization of proline´s clasp, we repeat our measurements using a cyclophilin mutant (R50A). This mutation is known to hinder the isomerase activity of the enzyme, but on the other hand does not affect substrate binding. a) Effect on the kinetics of *C. cellulolyticum* complex. The decay time of the samples incubated with cyclophilin R50A mutant is equivalent to the one of wild type sample. It must be noted the faster kinetics present in the case of the incubation with the wild type cyclophilin. B) Effect on *C. thermocellum* subpopulation ratios. The histograms show how attenuated the response of the system is when the sample is incubated with cyclophilin R50A (red line), especially when compared with the sample incubated with wild type cyclophilin (blue line). **C)** Effect of cyclophilin treatment on wild type c2A-celS complex. Two cyclophilin-treated c2A-celS samples are shown in blue, superimposed in top of a non-treated sample. After a few preliminary experiments, we observed that although the B1 population was systematically reduced on the samples treated with cyclophilin (see blue line histograms), the mild effect and the inherent variability of the samples made difficult to conclude an effect.

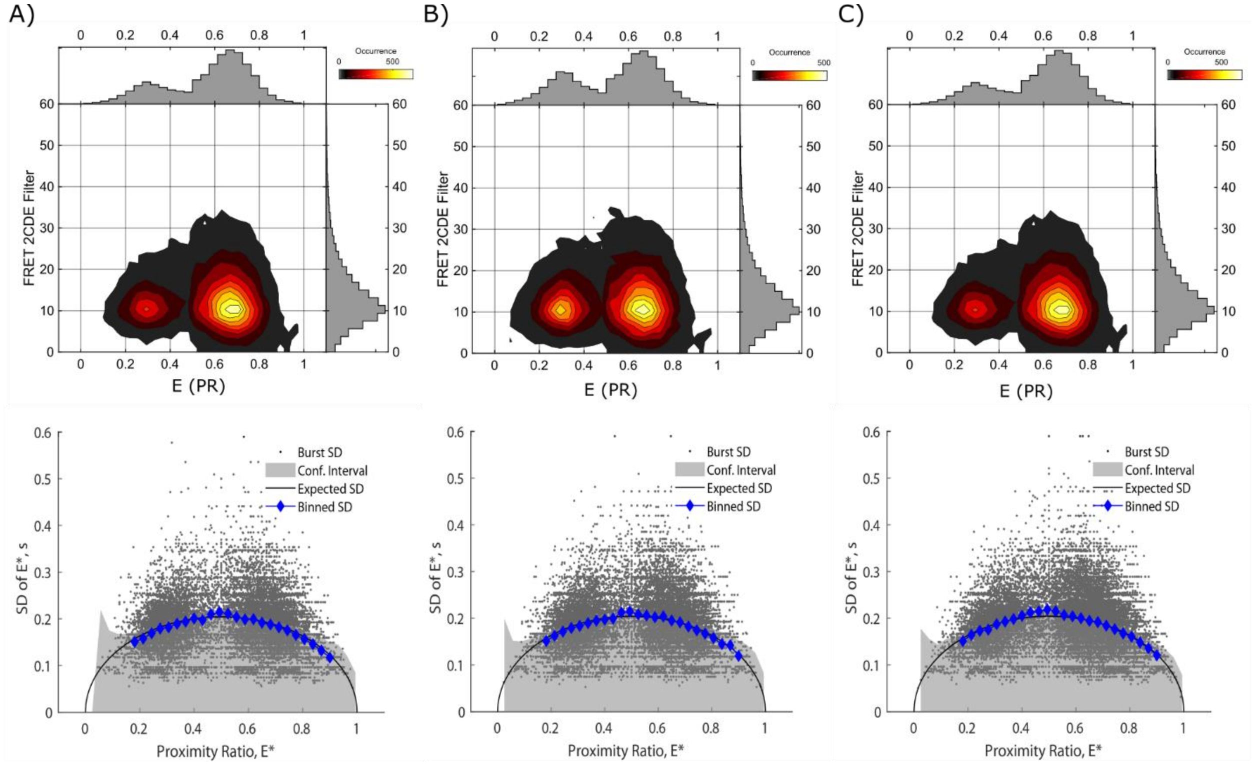

**Figure S12** *C. cellulolyticum* complexes are static in the burst duration scale. a), b) and c) show FRET 2CDE filter values and BVA analysis of c1C-CcCel5A, c1C-CcCel5A_mClasp and c1C-CcCel5A cyclophilin samples. In the top part the distribution of FRET 2CDE filter is represented, it can be seen that they peak at the value 10, as is expected for a static sample. The BVA analysis plots, displayed below, show that the standard deviation of the burst is always within the coefficient interval predicted for a static sample.

| Sample | R1 | Sigma1 | R2 | Sigma2 |
|---|---|---|---|---|
| c2A-celS | 67.35 ± 0.45 | 2.87 ± 1.27 | 48.94 ± 0.34 | 2.91 ± 0.55 |
| c2A-celA | 66.60 ± 0.90 | 3.52 ± 1.11 | 49.90 ± 0.51 | 3.63 ± 0.60 |
| c1A-celS | 70.46 ± 0.38 | 2.84 ± 1.23 | 48.56 ± 0.37 | 3.43 ± 0.56 |
| c1A-celA | 69.27 ± 0.41 | 3.06 ± 1.30 | 48.14 ± 0.41 | 4.32 ± 0.59 |
| c7A-celS | 74.16 ± 0.39 | 2.75 ± 1.42 | 54.10 ± 0.37 | 2.88 ± 0.49 |
| c7A-celA | 73.98 ± 0.49 | 3.02 ± 2.11 | 54.60 ± 0.38 | 3.19 ± 0.47 |
| c2A-celS_Swap | 67.20 ± 0.35 | 3.65 ± 0.57 | 49.32 ± 1.67 | 1.56 ± 2.00 |
| c1A-celS_ Swap | 71.23 ± 0.46 | 3.61 ± 1.00 | 50.36 ± 0.60 | 4.04 ± 0.81 |
| c7A-celS_ Swap | 73.25 ± 0.44 | 3.67 ± 0.98 | 54.69 ± 0.69 | 2.67 ± 0.73 |
| c1C-CcCel5A | 76.60 ± 0.81 | 4.15 ± 3.48 | 55.68 ± 0.34 | 2.59 ± 0.50 |
| c1C-CcCel5A_mClasp | 76.96 ± 1.96 | 5.46 ± 3.23 | 56.20 ± 0.40 | 2.92 ± 0.49 |
| c7C-CcCel5A | 74.27 ± 0.41 | 3.64 ± 1.87 | 53.78 ± 0.25 | 2.52 ± 0.37 |
| c7C-CcCel5AmClasp | 76.70 ± 2.46 | 3.64 ± 3.18 | 56.56 ± 1.02 | 2.52 ± 1.32 |
| c2A-celS_Y5A | 66.32 ± 0.85 | 3.49 ± 1.18 | 48.99± 0.33 | 2.85 ±0.50 |

**Supplementary Table 1**. Fitting parameters for all PDA fittings.
